# Enrichment analyses identify shared associations for 25 quantitative traits in over 600,000 individuals from seven diverse ancestries

**DOI:** 10.1101/2021.04.20.440612

**Authors:** Samuel Pattillo Smith, Sahar Shahamatdar, Wei Cheng, Selena Zhang, Joseph Paik, Misa Graff, Christopher Haiman, T.C. Matise, Kari E North, Ulrike Peters, Eimear Kenny, Chris Gignoux, Genevieve Wojcik, Lorin Crawford, Sohini Ramachandran

## Abstract

Since 2005, genome-wide association (GWA) datasets have been largely biased toward sampling European ancestry individuals, and recent studies have shown that GWA results estimated from self-identified European individuals are not transferable to non-European individuals due to various confounding challenges. Here, we demonstrate that enrichment analyses which aggregate SNP-level association statistics at multiple genomic scales—from genes to genomic regions and pathways—have been underutilized in the GWA era and can generate biologically interpretable hypotheses regarding the genetic basis of complex trait architecture. We illustrate examples of the robust associations generated by enrichment analyses while studying 25 continuous traits assayed in 566,786 individuals from seven diverse self-identified human ancestries in the UK Biobank and the Biobank Japan, as well as 44,348 admixed individuals from the PAGE consortium including cohorts of African-American, Hispanic and Latin American, Native Hawaiian, and American Indian/Alaska Native individuals. We identify 1,000 gene-level associations that are genome-wide significant in at least two ancestry cohorts across these 25 traits, as well as highly conserved pathway associations with triglyceride levels in European, East Asian, and Native Hawaiian cohorts.

## Introduction

Over the past two decades, funding agencies and biobanks around the world have made enormous investments to generate large-scale datasets of genotypes, exomes, and whole-genome sequences from diverse human ancestries that are merged with medical records and quantitative trait measurements ^1–8^. However, analyses of such datasets are usually limited to the application of standard genome-wide association (GWA) SNP-level association analyses, in which SNPs are tested one-by-one for significant association with a phenotype ^9–11^ (Table 1). Yet, even in the largest available multi-ancestry biobanks, GWA analyses fail to offer a comprehensive view of genetic trait architecture among human ancestries.

**Table 1:** The three genomic scales and corresponding association tests used in this study. The models of genetic trait architecture corresponding to each genomic scale and statistical method that have been previously invoked in the literature (including relevant examples cited in the last column). These nested genomic scales should routinely be leveraged in multi-ancestry GWA studies to generate biologically interpretable hypotheses of trait architecture across ancestries.

| Genomic Scale | Association Test | Model of Genetic Trait Architecture | Relevant Example |
| --- | --- | --- | --- |
| SNPs | Standard univariate genome-wide association (GWA) test | The true mutation-level trait architecture is the same for all individuals. | Many inflammatory bowel disease mutations replicate across ancestries<sup>49</sup>. |
| SNP-Sets/Genes | Gene-level association tests (e.g., gene-ε<sup>38</sup>, SKAT<sup>40</sup>) | Core genes are the same across all ancestries, with potentially varying causal SNPs. | Late-onset Alzheimer’s disease risk from *ApoE4* allele is lower in African ancestry individuals<sup>89</sup>. |
| Pathways/Networks | Pathway analysis and network propagation (e.g., Hierarchical HotNet<sup>31</sup>, RSS<sup>60</sup>). | Core genes differ across ancestries, but are all in the same annotated pathway. | Skin pigmentation architecture in the same pathway differs between African and European ancestry individuals<sup>2</sup>. |

**Table 2:** Effect size homogeneity in variants identified as significant in the European or East Asian cohorts. In each of the 25 traits analyzed in this study a majority of variants that are significant in at least the European or East Asian cohorts had the same direction of effect in the other ancestry cohort.

| Trait | Number of significant SNPs in at least the European or East Asian cohort | Number of SNPs with same direction of effect | Percentage of SNPs with same direction of effect |
| --- | --- | --- | --- |
| BMI | 6,374 | 4,456 | 69.91 |
| Basophil | 884 | 520 | 58.82 |
| CRP | 694 | 463 | 66.71 |
| Cholesterol | 3,451 | 2,379 | 68.94 |
| DBP | 1,962 | 1,320 | 67.28 |
| EGFR | 7,129 | 5,038 | 70.67 |
| Eosinophil | 5,890 | 3,608 | 61.26 |
| HBA1C | 3,030 | 2,206 | 72.81 |
| HDL | 5,995 | 4,077 | 68.01 |
| Height | 33,577 | 22,090 | 65.79 |
| Hematocrit | 5,382 | 3,602 | 66.93 |
| Hemoglobin | 5,280 | 3,601 | 68.20 |
| LDL | 2,521 | 1,929 | 76.56 |
| Lymphocyte | 2,214 | 1,237 | 55.87 |
| MCH | 6,763 | 4,674 | 69.11 |
| MCHC | 1,760 | 1,124 | 63.86 |
| MCV | 7,489 | 5,208 | 69.54 |
| Monocyte | 3,929 | 2,565 | 65.28 |
| Neutrophil | 5,431 | 3,850 | 70.89 |
| PLC | 11,014 | 7,161 | 65.02 |
| RBC | 9,211 | 6,263 | 67.99 |
| SBP | 1,807 | 1,182 | 65.41 |
| Triglyceride | 3,743 | 2,686 | 71.76 |
| WBC | 6,017 | 4,105 | 68.22 |
| Urate | 5,864 | 3,787 | 64.58 |

SNP-level GWA results are difficult to interpret across multiple human ancestries due to a litany of confounding variables, including: (*i*) ascertainment bias in genotyping ^2, 5^, (*ii*) varying linkage disequilibrium (LD) patterns ^12, 13^, (*iii*) variation in allele frequencies due to different selective pressures and unique population histories ^13–17^, and (*iv*) the effect of environmental factors on phenotypic variation ^18–21^. These confounders and the observed low transferability of GWA results across ancestries ^2, 22, 23^ have generated an important call for increasing GWA efforts focused on populations of diverse, non-European ancestry individuals.

We also note, as other studies have ^6, 24^, that the GWA SNP-level test of association is rarely applied to non-European ancestry individuals ^25^. There are two likely explanations for leaving non-European ancestry individuals out of analyses: (*i*) researchers are electing to not analyze diverse cohorts due to a lack of statistical power and concerns over other confounding variables (recently covered in Ben-Eghan et al. ^24^); or (*ii*) the analyses of non-European cohorts yield no genome-wide significant SNP-level associations. In either case, valuable information is being ignored in GWA studies or going unreported in resulting publications ^24–26^. Even when diverse ancestries are analyzed, GWA studies usually condition on GWA results identified using European ancestry cohorts to detect and give validity to other SNP-level associations ^6^. In our own analysis of abstracts of publications between 2012 and 2020 using UK Biobank data, we found that only 33 out of 166 studies (19.87%) reported genome-wide significant associations in any non-European ancestry cohort (Figure S1 - Figure S2). Focusing energy and resources on increasing GWA sample sizes without intentional focus on sampling of non-European populations will thus likely perpetuate an already troubling history of leaving non-European ancestry samples out of GWA analyses of large-scale biobanks such as the UK Biobank ^24^ (but see Sinnott-Armstrong et al. ^27^). However, we note that non-European ancestry GWA studies have — and will continue to have — smaller sample sizes than existing and emerging European-ancestry GWA cohorts, limiting the precision of effect size estaimates in these studies. What has received less attention than the need to improve GWA study design is the potential of enrichment analyses to characterize genetic trait architecture in multi-ancestry datasets while accounting for variable statistical power to detect, estimate, and replicate genetic associations among cohorts.

In this analysis, we illustrate that focusing solely on *p*-values from the standard GWA framework is insufficient to capture the genetic architecture of complex traits. Specifically, we propose that expansion of association analyses to the genomic scale of genes and pathways generates robust and interpretable hypotheses about trait architecture in multi-ancestry cohorts. These enrichment analyses can increase the power to detect genes through the aggregation of SNPs of small effect (which explain the majority of the heritability of most traits ^28, 29^). Mathieson ^30^ recently highlighted the pattern of homogeneity of direction of effect in multi-ancestry studies even when individual SNPs are not categorized as genome-wide significant in multiple ancestries. Gene and pathway level analyses can prioritize biological regions where there is homogeneity in the direction of effect, generating biologically interpretable hypotheses for the genetic architecture of complex traits in multiple ancestry cohorts.

In this study of 25 quantitative traits and more than 600,000 diverse individuals from the UK Biobank (UKB), BioBank Japan (BBJ), and the PAGE study data (Table S1 - Table S9), we detail biological insights gained from the application of gene and pathway level enrichment analyses to seven diverse ancestry cohorts. We perform genetic association tests for SNPs, genes, and pathways across multiple ancestry groups with a trait of interest. We test for significantly mutated subnetworks of genes using known protein-protein interaction networks and the Hierarchical HotNet software ^31^. Enrichment analyses do not require generating additional information beyond standard GWA inputs (or outputs, for methods that take in GWA summary statistics). We demonstrate that moving beyond SNP-level associations allows for a biologically comprehensive prioritization of shared and ancestry-specific mechanisms underlying genetic trait architecture.

## Materials and Methods

### Data overview

We performed statistical tests of association at the SNP, gene, and pathway-level for 25 quantitative traits. These analyses were performed on data seven ancestry cohorts drawn from the UK Biobank, BioBank Japan (BBJ), and PAGE consortia (Table S3). Descriptions of the samples studied, including sample size, and number of SNPs for each ancestry cohort are given in Table S1 and Table S4 - Table S9. For an extensive description of each cohort from the three biobanks that we analyze in this study, see the Supplemental Information.

### SNP-level GWA analyses

In the European, African, and South Asian ancestry cohorts from the UK Biobank, we performed GWA studies for each ancestry-trait pair in order to test whether the same SNP(s) are associated with a given quantitative trait in different ancestries. SNP-level GWA effect sizes were calculated using plink and the --glm flag ^32^. Age, sex, and the first twenty principal components were included as covariates for all traits analyzed ^7^. Principal component analysis was performed using flashpca 2.0^33^ on a set of independent markers derived separately for each ancestry cohort using the plink command --indep-pairwise 100 10 0.1. Summary statistics for the 25 quantitative traits in the Biobank Japan, as well as available ancestry-trait pairs in the PAGE study data, were then compared with the results from the association analyses in the UK Biobank cohorts (same traits as listed in Table S4). In each analysis of an ancestry-trait pair, a separate Bonferonni-corrected significance threshold was calculated using the number of SNPs tested in that particular ancestry-trait pair (Table S10).

### Comparison of SNP level fine-mapping methods and effect size direction

We compared the results of two fine-mapping methods, SuSie ^34^ and PESCA ^35^, when applied to SNP-level summary statistics in the European (UKB) and East Asian (BBJ) cohorts. SuSie is an iterative Bayesian stepwise selection method that identifies a credible set of SNPs that contribute to a phenotype of interest ^34^. Using the effect sizes and standard errors generated from the standard GWA framework for each trait in each ancestry, we applied SuSiE in order to identify probable sets of causal SNPs. We then found the correlation between the posterior inclusion probabilities of each SNP in the European and East Asian cohorts (Table S11 - Table S13).

Unlike SuSiE, the PESCA framework is explicitly designed for identifying shared SNP-level association signals between multiple ancestry cohorts versus ancestry-specific associations ^35^. In addition to GWA summary statistics, PESCA uses information about the correlation structure between SNPs (i.e., LD) to identify SNPs that are likely to be causal in two cohorts of interest. Shi et al. ^35^ analyzed seven continuous traits in the European (UKB) and East Asian (BBJ) cohorts using PESCA and produced posterior probabilities that individual SNPs were: (*i*) associated with a phenotype in both the both cohorts, (*ii*) associated with the trait of interest only in the European cohort, or (*iii*) associated with the trait of interest only in the East Asian cohort. For each trait, we calculated the number of SNPs that were nominally significant (*p*-value *<* 10*^−^*5, as in the original PESCA analysis) in the standard GWA framework in both the European and East Asian cohorts and had a PESCA posterior probability of being associated in both ancestries *>* 0.8 (see Table S14). We also found the number of SNPs that had a PESCA posterior probability of being associated in both ancestries *>* 0.8 that were only identified as significant in one ancestry using the GWA framework.

Finally, we explored the recent proposition of Mathieson ^30^ that the direction of effect for SNP-level summary statistics might be conserved among ancestry cohorts even if those variants are not genome-wide significant in either cohort. To that end, for each of the 25 traits that we analyzed, we compared the direction of SNP effect sizes between the European and East Asian ancestry cohorts. We were only able to carry this analysis out for variants that were genotyped in both cohorts (Table S14). For each remaining nominally significant variant, we stored the direction of the effect size and checked the direction of effect size in the other ancestry. When zero was included within the range of the effect size plus or minus one standard deviation, we assumed the SNP did not have the same direction in both cohorts. The results of our comparison are shown in Table S14. We note that this test may be confounded by the precision of effect size estimation and warrants further exploration, including an analysis of local false sign discovery rates (see ^36, 37^).

### Gene-level association tests

In order to test aggregated sets of SNP-level GWA effect sizes for enrichment of associated mutations with a given quantitative trait, we applied gene-*ε*^38^ to each ancestry cohort we studied for each trait of interest, resulting in 125 sets of gene-level association statistics (Table S3, Table S15). The gene-*ε* method takes two summary statistics as input: (*i*) SNP-level GWA marginal effect size estimates ***β***^^^ derived using ordinary least squares and (*ii*) an LD matrix **Σ** empirically estimated from external data (e.g., directly from GWA study genotype data, a matrix estimated from a population with similar genomic ancestry to that of the samples analyzed in the GWA study). It is well-known that SNP-level effect size estimates can be inflated due to various correlation structures among genome-wide genotypes. gene-*ε* uses its inputs to derive regularized effect size estimates through elastic net penalized regression.

In practice, gene-*ε* and other enrichment analyses ^39–41^ can be applied to any user-specified set of genomic regions, such as regulatory elements, intergenic regions, or gene sets. These gene-level association tests enable identifying traits in which genetic architecture may be heterogeneous among individuals at the SNP-level across individuals; applying gene-*ε* in multiple ancestry cohorts allows researchers to further test whether genes associated with a trait of interest are the same, or vary, across ancestries. gene-*ε* takes as input a list of boundaries for all regions to be tested for enrichment of associations. In our study, we applied gene-*ε* to all genes and transcriptional elements defined in Gusev et al. ^42^ for human genome build 19. Throughout this study, we refer to the resulting effect size estimates produced by gene-*ε* as “gene-level association statistics”.

In our gene-level analysis, SNP arrays included both genotyped and high-confidence imputed SNPs (information score *≥* 0.8) for each ancestry-trait pair. To compute the LD matrix, we first pruned highly linked SNPs so that the number of SNPs included for any chromosome was less than 35,000 SNPs — the computational limit of gene-*ε* due to the size of the LD matrix — using the plink command --indep-pairwise 100 10 0.5. For the UK Biobank European, South Asian, and African ancestry cohorts, we then derived empirical LD estimates between each pair of SNPs for each chromosome in each cohort using plink flag --r square applied to the empirical genotype and high-confidence imputed data. The ancestry-specific SNP arrays were then used to calculate 23,603 gene-level association statistics for the European ancestry cohort, 23,671 gene-level association statistics for the South Asian ancestry cohort, and 23,575 gene-level association statistics for the African ancestry cohort.

To calculate gene-level association statistics using Biobank Japan summary statistics, we first found the intersection between SNPs included in the analysis of each trait (Table S4) and SNPs included in the 1000 Genomes Project phase 3 data for the sample of 93 JPT individuals. Note, this intersection was different among some traits as the genotype data in the Biobank Japan were from different studies, which in turn used different genotyping arrays. We then pruned highly linked markers for each trait separately using the plink flag --indep-pairwise 100 10 X where X was determined by finding the highest possible value that led to less than 35,000 SNPs being included on each chromosome for the trait. Because of the increased density of SNPs with effect size estimates for Height, X was set to prune more conservatively at X = 0.15. For all other traits, X was set to 0.5. The number of regions for which a gene-*ε* gene-level association statistic was calculated for each trait is given in Table S4.

GWA summary statistics for the five cohorts in the PAGE study data were used as input to gene-*ε* for each available ancestry-trait combination. The array of markers for each ancestry cohort in the PAGE study data was pruned using plink flag --indep-pairwise 100 10 X. X was set to the maximum value in each ancestry that ensured no chromosome contained more than 35,000 markers: X was set to 0.05 for the African-American cohort, 0.08 for the Hispanic and Latin American and AIAN cohorts, and 0.25 for the Native Hawaiian cohorts. Finally, for each ancestry-trait combination, genes that passed the Bonferroni corrected *p*-value (*p* = 0.05/number of genes tested) were labeled as “significant” throughout this study (see Table S15 for specific thresholds).

### Pathway analysis and network propagation using Hierarchical HotNet

We tested for shared and divergent gene-level association results among interacting genes for each trait of interest using network propagation of gene-*ε* gene-level association statistics as input to Hierarchical HotNet ^31^. Hierarchical HotNet identifies significantly altered subnetworks using gene-level scores as input; in this study, these gene scores were set to *−*log_10_-transformed *p*-values of gene-*ε* gene-level association test statistics (see also Nakka et al. ^41, 43^). For each ancestry-trait combination, we assigned *p*-values of 1 to genes with *p*-values greater than 0.1 to make the resulting networks both sparse and more interpretable (again see Nakka et al. ^41, 43^). In addition, ancestry-trait pairs in which less than 25 genes produced gene-*ε p*-values less than 0.1 were discarded as there were an insufficient number of gene-level statistics to populate the protein-protein interaction networks.

We used three protein-protein interaction networks: ReactomeFI 2016^44^, iRefIndex 15.0^45^, and HINT+HI ^46, 47^. For the ReactomeFI 2016 interaction network, interactions with confidence scores less than 0.75 were discarded. The HINT+HI interaction network consists of the combination of all interactions in HINT binary, HINT co-complex, and HuRI HI interaction networks. We ran Hierarchical HotNet (10^3^ permutations) on the thresholded -log_10_-transformed gene-level *p*-values for each ancestry-trait combination. We restricted our further investigation to the largest subnetwork identified in each significant ancestry-trait-interaction network combination (*p <* 0.05).

## Results

### SNP-level replication of GWA results among ancestries is the exception, not the rule

Multiple recent studies have interrogated the extent to which SNP-level associations for a given trait replicate across ancestries, both empirically and under a variety of models (see Wojcik et al. ^6^, Durvasula and Lohmueller ^22^, Shi et al. ^35^, Carlson et al. ^48^, Liu et al. ^49^, Eyre-Walker ^50^, Shi et al. ^51^). To extensively compare variant-level associations among the seven ancestry cohorts that we analyzed, we first examined the number of genome-wide significant SNP-level associations that replicated exactly based on chromosomal position and rsID in multiple ancestries (see Figure S3a and Figure S3c, with Bonferroni-corrected thresholds provided in Table S10). Exact replication of at least one SNP-level association across two or more ancestries occurs in all 25 traits that were studied. The C-reactive protein (CRP) trait had the highest proportion of replicated SNP associations in multiple ancestries, with 18.95% replicating using the standard GWA framework in at least two ancestries, but has a relatively low number of unique GWA significant SNPs (2,734) when compared to other traits (Figure 1). This is likely because the genetic architecture of CRP is sparse and highly conserved across ancestries, as is shown in Figure 2. We note that the concordance of genome-wide significant SNP-level association statistics for CRP among five ancestry cohorts is exceptional. In the other 24 traits we analyzed, we did not observe any SNP-level replication among five cohorts. C-reactive protein, which is encoded by the gene of the same name located on chromosome 1, is synthesized in the liver and released into the bloodstream in response to inflammation. In our standard GWA analysis of SNP-level association signal in each ancestry cohort with CRP, rs3091244 is genome-wide significant in a single ancestry cohort. rs3091244 has been functionally validated as influencing C-reactive protein levels ^52, 53^ and is linked to genome-wide significant SNPs in the other two ancestries for which genotype data is available. Interestingly, all GWA significant SNP-level associations for CRP in the Native Hawaiian ancestry cohort replicate in both the African-American (PAGE) and the Hispanic and Latin American cohorts (these three cohorts were all genotyped on the same array).

**Figure 1:**
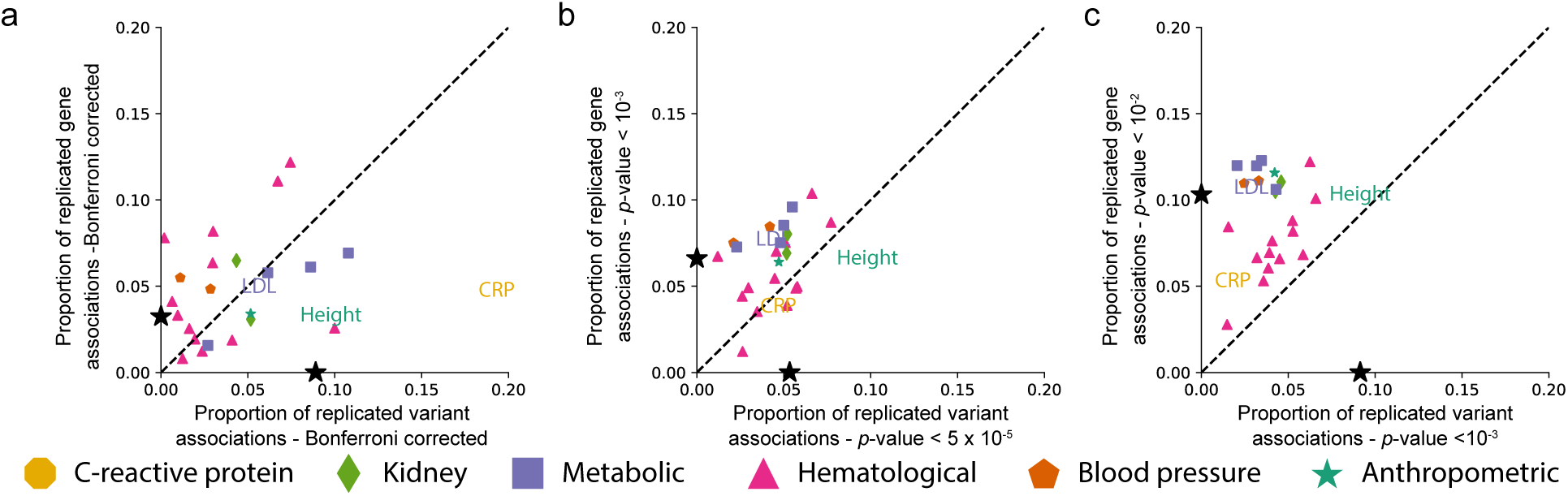
Less stringent significance thresholds lead to a decrease in the proportion of replicated SNP-level associations and an increase in the proportion of gene-level associations among ancestries for each of the 25 traits analyzed. **a.** Proportion of all SNP-level Bonferroni-corrected genome-wide significant associations in any ancestry that replicate in at least one other ancestry is shown on the x-axis (see Table S10 for ancestry-trait specific Bonferroni corrected *p*-value thresholds). On the y-axis we show the proportion of significant gene-level associations that were replicated for a given phenotype in at least two ancestries (see Table S15 for Bonferroni corrected significance thresholds for each ancestry-trait pair). The black stars on the x- and y-axes represent the mean proportion of replicates in SNP and gene analyses, respectively. C-reactive protein (CRP) contains the greatest proportion of replicated SNP-level associations of any of the phenotypes. **b.** The x-axis indicates the proportion of SNP-level associations that surpass a nominal threshold of *p*-value *<* 10*^−^*^5^ in at least one ancestry cohort that replicate in at least one other ancestry cohort. The y-axis indicates the proportion of gene-level associations that surpass a nominal threshold of *p*-value *<* 10*^−^*^3^ in at least one ancestry cohort and replicate in at least one other ancestry cohort. Nominal *p*-value thresholds tend to decrease the proportion of replicated SNP-level associations and tend to increase the proportion of replicated gene-level associations. The number of unique SNPs and genes that replicated in each cohort is given in Figure S15. **c.** The x-axis indicates the proportion of SNP-level associations that surpass a nominal threshold of *p*-value *<* 10*^−^*^3^ in at least one ancestry cohort that replicate in at least one other ancestry cohort. The y-axis indicates the proportion of gene-level associations that surpass a nominal threshold of *p*-value *<* 10*^−^*^2^ in at least one ancestry cohort and replicate in at least one other ancestry cohort. The number of unique SNPs and genes that replicated in each cohort is given in Figure S16. As shown in panel b, nominal *p*-value thresholds tend to decrease the proportion of replicated SNP-level associations and tend to increase the proportion of replicated gene-level associations. Expansion of three letter trait codes are given in Table S2 and a version of this plot with all trait names displayed as text is shown in Figure S14. Figure S14 shows the same set of plots with all traits represented as text.

**Figure 2:**
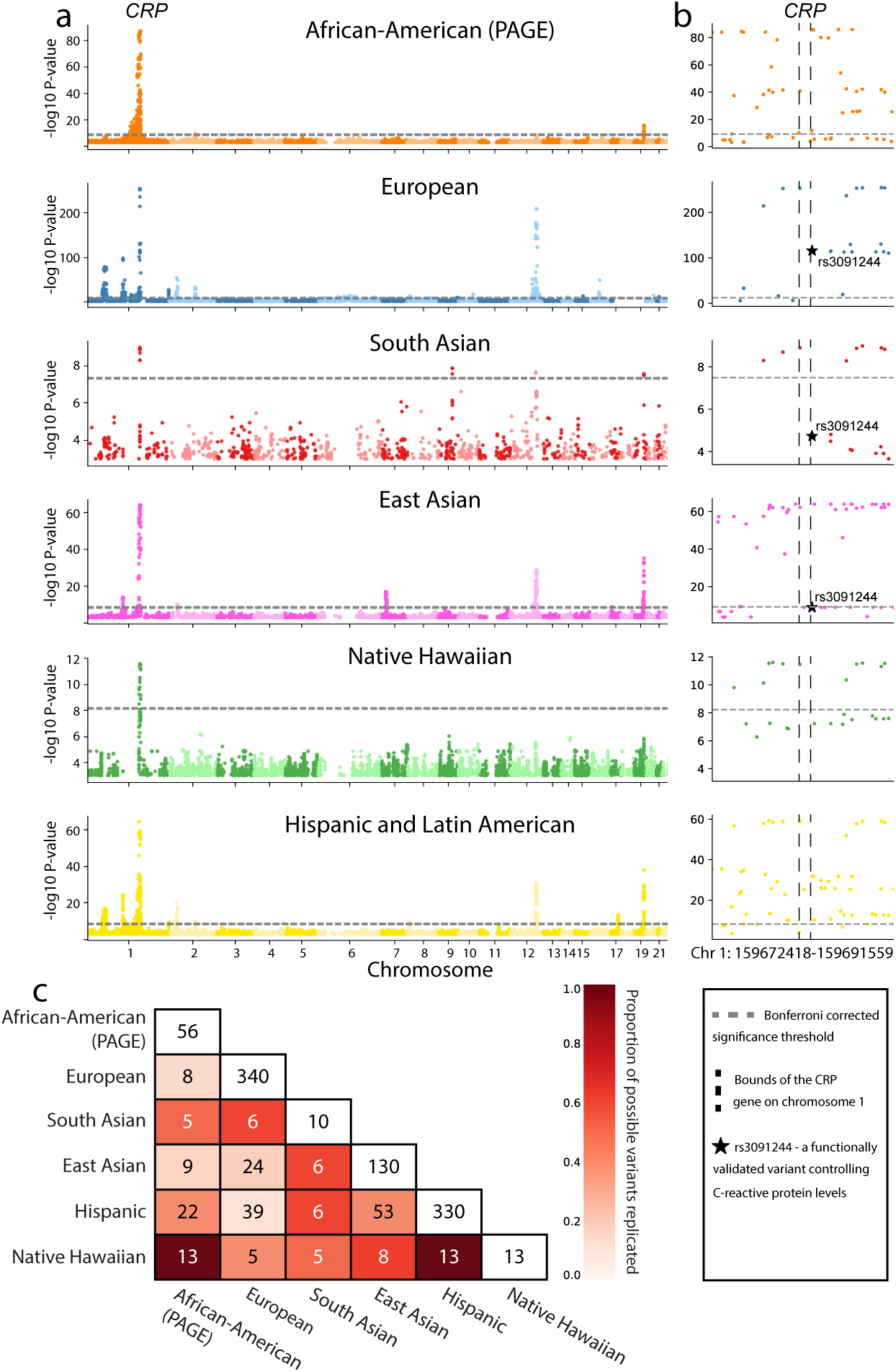
C-reactive protein is an exceptional trait where standard GWA analyses may be sufficient to identify shared genetic architecture among ancestry cohorts. **a.** Manhattan plot for SNP-level associations with C-reactive protein levels. Ancestry-specific Bonferroni-corrected significance thresholds are shown with dashed horizontal grey lines and listed in Table S10. Note that the scale of the -log_10_- transformed *p*-values on the y-axis is different for each ancestry for clarity. **b.** Manhattan plot of SNP-level associations around the *CRP* gene located on chromosome 1 for each ancestry (zoomed in from panel **a**). Boundaries of the *CRP* gene are shown with vertical dashed black lines. All six ancestries contain genome-wide significant SNPs in the region. Black stars in the European, South Asian, and East Asian plots represent rs3091244, a SNP that has been functionally validated as contributing to serum levels of C-reactive protein ^52, 53^. **c.** Heatmap of Bonferroni-corrected significant genotyped SNPs replicated between each pair of ancestries analyzed. Here, we focus on SNPs in the 1MB region surrounding the *CRP* gene. Entries along the diagonal represent the total number of SNP-level associations in the 1MB region surrounding the *CRP* gene for each ancestry. The color of each cell is proportional to the percentage of SNP-level associations replicated out of all possible replications in each ancestry pair (i.e., the minimum of the diagonal entries between the pairs being considered). For example, the maximum number of genome-wide significant SNPs that can possibly replicate between the Hispanic and East Asian is 25, and 20 replicate resulting in the cell color denoting 80% replication. A similar matrix, computed including imputed SNPs is shown in Figure S17.

**Figure 3:**
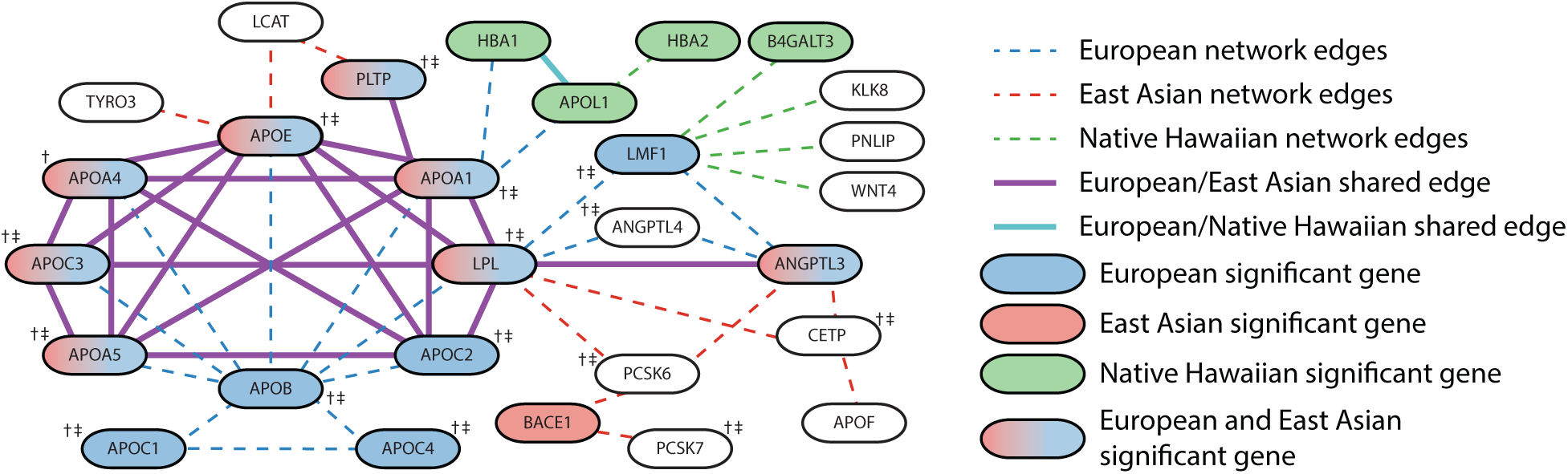
A subnetwork of apolipoprotein genes is significantly enriched for mutations in European, East Asian, and Native Hawaiian ancestries associated with triglyceride levels. The largest significantly altered subnetwork (*p*-value *<* 0.05) for triglyceride levels contains overlapping gene subnetworks for each of the European, East Asian, and Native Hawaiian ancestries when analyzed independently with Hierarchical HotNet ^31^. Each node in the network represents a gene. The shading of each node indicates the statistical significance of the association of that gene with triglyceride levels in a particular cohort. Two genes are connected if their protein products interact based on the ReactomeFI 2016^44^ (European, East Asian) or iRefIndex 15.0^45^ (Native Hawaiian) protein-protein interaction networks. Several genes from the apolipoprotein gene family are significantly associated with triglyceride levels in both the European and East Asian cohorts (see Data Availability). Additionally, the interactions between them form a highly connected subnetwork. Smaller subnetworks identified in the Native Hawaiian cohort are distal modules that are connected to the subnetwork detected in the European cohort. Not all genes in the largest significantly altered subnetwork for the Native Hawaiian ancestry group are shown for visualization purposes (127 not pictured here). Genes that contain SNPs previously associated to triglyceride levels in a European cohort in the GWAS catalog are indicated with. Similarly, genes that contain SNPs previously associated with triglyceride levels in a non-European cohort in the GWAS Catalog are indicated with. The studies identifying these associations are given in Table S16.

In the other 24 traits, the proportion of genome-wide SNP-level replications was below 10% (Figure 1a). For polygenic traits, replication of SNP-level GWA results is challenging to interpret considering the large number of GWA significant associations for the trait overall. For example, height contains the largest number of replicated SNP-level associations in our multi-ancestry analysis — but these only represent 8.90% of all unique SNP-level associations with height discovered in any ancestry cohort. A more comprehensive discussion of previously associated SNPs is available for both height and CRP in the Supplemental Information.

### Fine-mapping methods have variable efficacy in identification of SNP level associations among ancestry cohorts

Often, replication of GWA results across cohorts is tested using genomic regions centered on a SNP. Scans across the region surrounding the SNP of interest are usually defined arbitrarily — using physical windows (or “clumps”) to smooth over ascertainment bias and varying LD across cohorts or ancestries instead of using regions that are biologically annotated such as genes or transcriptional elements. While clumping presents an easy way to scan for regional replication of a given GWA finding, the corresponding results are not readily interpretable when prioritizing GWA results for downstream validation. We performed clumping using windows of size 1Mb centered around significant SNP-level associations (see Materials and Methods). Height had the largest proportion of windows that contain a SNP-level association that replicated in at least two ancestries (Figure S3b and Figure S3e). In the three traits with the greatest proportion of windows containing SNP-level replications — height (77.09% of clumps), urate (65.89%), and low density lipoprotein (54.40%) — we then recorded the number of genes and transcriptional elements within the window that contained GWA significant SNP-level associations. We found that for all three traits, the vast majority of 1Mb windows that were used to clump SNP-level associations contained multiple genes and transcriptional elements with significantly associated SNPs: height (94.04%, 17.93 genes in clump (mean) *±* 15.71 (standard deviation)), urate (97.47%, 18.44 *±* 13.72), and low density lipoprotein (99.12%, 14.85 *±* 12.89). Thus, we find window-based clumping does not easily produce biologically interpretable hypotheses for downstream validation.

Recent analyses of multi-ancestry GWA cohorts have also tested for effect size heterogeneity ^6, 34, 51, 54, 55^. We applied the fine-mapping method SuSiE ^34^ to identify signals of effect size heterogeneity in the three ancestry cohorts for which we had access to raw genotype data (UK Biobank European ancestry, African ancestry, and South Asian ancestry individuals; see Table S1). We find little evidence of correlated SuSiE effect size estimates among ancestry cohorts, including among independent subsamples of the UK Biobank European ancestry individuals Table S11 - Table S13. In addition, we applied PESCA (a method developed by Shi et al. ^51^) to the results of our SNP-level analysis to understand how the modeling of LD to affected the power to identify probably causal SNPs shared in the European and East Asian ancestry cohorts. PESCA improves upon standard clumping approaches by modeling the LD in a region to identify SNPs that are likely to be causal for the same trait in multiple ancestries. In a comparison with the results from seven continuous traits analyzed in the original study ^51^, we found that the vast majority of SNPs identified by PESCA as causal (posterior probability *>* 0.8) in both ancestries were also nominally significant in our SNP-level association results (see Table S14). Both SuSiE ^34^ and PESCA ^51^ demonstrate the utility of modeling variation in LD structure among ancestries when conducting multi-ancestry GWA studies.

Recently, Mathieson ^30^ proposed the hypothesis that the direction of effect sizes is the same among ancestries, even when the effects are not genome-wide significant. To test this, we compared the direction of effect in SNPs that were significant in either the European or East Asian ancestry cohort to the direction of the effect in the other ancestry where the SNP was tested using the standard GWA framework. We limit the comparison to the European and East Asian cohorts due to their large sample sizes which increases the precision of effect size estimates. Table S14 shows the number of variants that were significantly associated with each trait in at least one of the European and East Asian ancestry cohorts, and also displays the number of those variants that have the same direction of effect as the significant variant in the other ancestry. In the 25 traits that we analyzed, the direction of effect was conserved in both the European and East Asian ancestry cohorts (between effect direction concordance from 55.87% and 76.56% of SNPs across 25 traits). The remaining SNPs where the direction of the effect size was not conserved represent those SNPs that: (*i*) had different direction of effect size, (*ii*) were not tested in both ancestry cohorts, or (*iii*) had effect size estimates within one standard error of zero (Table S14). The observed conservation of effect size direction in multiple ancestry cohorts, even when SNPs are non-significant in one or more cohorts, is a primary assumption of regional enrichment methods and supports Mathieson ^30^ ’s hypothesis and findings. This suggests that regional enrichment methods, which are sensitive to shared patterns of effect size direction among cohorts, are a natural approach to apply to GWA summary statistics even in the absence of replication SNP-level GWA signals among cohorts.

### Valuing biological mechanism over statistical significance

Enrichment analyses aggregate SNP-level association statistics using predefined SNP sets, genes, and pathways to identify regions of the genome enriched for trait associations beyond what is expected by chance. Published enrichment analyses have demonstrated the ability to identify trait associations that go unidentified when using standard SNP-level GWA analysis ^38, 40, 56–60^. The standard GWA method is known to have a high false discovery rate (FDR) ^36, 61^, which enrichment analyses can mitigate. Our analyses in Figure S4 and Figure S5 illustrate that two methods — regression with summary statistics (RSS) ^60^, a fully Bayesian method; and gene-*ε*^38^ — control FDR particularly well both in the presence and absence of population structure. Enrichment methods also increase power for identifying biologically interpretable trait associations in studies with smaller sample sizes than present-day GWA studies. For example, Nakka et al. ^41^ identified an association between *ST3GAL3* and attention deficit hyperactivity disorder (ADHD) using methods that aggregated SNP-level signals across genes and networks. ADHD is a trait with heritability estimates as high as 75% which had no known genome-wide significant SNP-level associations at the time; Nakka et al. ^41^ studied genotype data from just 3,319 individuals with cases, 2,455 controls and 2,064 trios ^62^. A study by Demontis et al. ^63^ later found a SNP-level association in the *ST3GAL3* gene, but was only able to do so with a cohort an order of magnitude larger (20,183 individuals diagnosed with ADHD and 35,191 controls, totaling 55,374 individuals).

Because non-European GWA ancestry cohorts usually have much smaller sample sizes compared to studies with individuals of European ancestry, enrichment analyses offer a unique opportunity to boost statistical power and identify biologically relevant genetic associations with traits of interest using multiancestry datasets. In a simulation study using synthetic phenotypes generated from the European and African ancestry cohorts in the UK Biobank, we show that gene-*ε* is able to identify significantly associated genes even in smaller cohorts (*N* = 10, 000 and *N* = 4, 967 in the European and African ancestry cohorts, respectively) without the inflated false discovery rate that is often exhibited by the standard GWA framework (Figure S6 and Figure S7). Additionally, in these simulations, gene-*ε* correctly identifies “causal” genes that are commonly associated in both cohorts (Figure S8 and Figure S9). These simulations illustrate the utility of modeling LD (and in the case of gene-*ε*, additionally shrinking inflated effect sizes) information to identify enrichment of SNP-level associations in predefined SNP sets.

In an analysis performed by Ben-Eghan et al. ^24^ on 45 studies analyzing UK Biobank data, the second most commonly stated reason for omitting non-European cohorts in applied GWA analyses was due to lack of power for identifying SNP-level GWA signals. We tested for gene-level associations in each of the 25 complex traits in each ancestry cohort for which we had data (Table S1 - Table S9), and identified associations in genes and transcriptional elements shared across ancestries for every trait. All of our analyses discussed here used gene-*ε* (see performance comparison with other enrichment analyses in Cheng et al. ^38^ and Figure S6 - Figure S9), an empirical Bayesian approach that aggregates SNP-level GWA summary statistics, where *p*-values for each gene are derived by constructing an empirical null distribution based on the eigenvalues of a gene-specific partitioning of the LD matrix (for more details, see Cheng et al. ^38^). Our analyses show that several hematological traits have a higher rate of significant gene-level associations that replicate across multiple ancestry cohorts than SNP-level associations that replicate across ancestry cohorts (Figure 1b). These include platelet count (PLC), mean corpuscular hemoglobin (MCH), mean corpuscular hemoglobin concentration (MCHC), hematocrit, hemoglobin, mean corpuscular volume (MCV), red blood cell count (RBC), and neutrophil count (Figure S3f). Focusing on platelet count as an example, we identify 65 genes that are significantly enriched for associations in multiple ancestries when tested using gene-*ε* (see Table S15 for details on Bonferroni thresholds used to correct for the number of genes tested) ^38^. Fifty-five of these genes are significantly associated in both the European and East Asian ancestry cohorts, and the remaining ten all replicate in other pairs of ancestry cohorts. Overall, each of the six ancestry cohorts in our analysis share at least one significant gene with another ancestry cohort, as shown in Figure S10.

Results from gene-level enrichment analyses can be further propagated on protein-protein interaction networks to identify interacting genes enriched for association signals ^64^. Often, studies use network propagation as a way to incorporate information from multiple “omics” databases in order to identify significantly mutated gene subnetworks or modules contributing to a particular disease ^65^. An unexplored extension of network propagation is how it can be used with multi-ancestry GWA datasets to identify significantly mutated subnetworks that are shared or ancestry-specific ^43^.

To conduct network propagation of gene-level association results in our analyses, we applied the Hierarchical HotNet method ^31^ to gene-*ε* gene-level association statistics for each trait-ancestry data set. In 3, we display the significant (*p*-value *<* 0.01) network results for triglyceride levels in three ancestry cohorts: European, East Asian, and Native Hawaiian (networks separated by ancestry are available in Figure S11). In both the European and East Asian cohorts, we identify enrichment of mutations in a highly connected subnetwork of genes in the apolipoprotein family. In addition, we identify a gene subnetwork enriched for mutations in the East Asian and Native Hawaiian cohorts that interacts with the significantly mutated sub-network identified in both the European and East Asian cohorts. For instance, beta-secretase 1 (*BACE1*), is a genome-wide significant gene-level association in the East Asian cohort but does not contain SNPs previously associated with triglycerides in any ancestry cohort in the GWAS catalog. Additionally, both *APOL1* and *HBA1* were identified as significantly associated with triglycerides using gene-*ε* in our analysis of the Native Hawaiian ancestry cohort, and both genes were part of significant subnetworks identified by Hierarchical HotNet in the European and Native Hawaiian ancestry cohorts. Details on replicated SNP-level and gene-level associations among ancestries for triglyceride levels are shown in Figure S12 and Figure S13, respectively. SNP-level and gene-level association results are further discussed for both platelet count and triglyceride levels in the Supplemental Information.

## Discussion

Many recent studies have proposed changes to multi-ancestry GWA study design ^2, 5, 6, 22, 24–27, 66, 67^. In this analysis, we have focused on the potential of *methods* to increase the insight gained into complex trait architecture from multi-ancestry GWA datasets via the generation of biologically interpretable hypotheses. We demonstrate the potential gains of moving beyond standard SNP-level GWA analysis using 25 quantitative complex traits among seven human ancestry cohorts in three large biobanks: BioBank Japan and the UK Biobank, and the PAGE consortium database (Table S1 - Table S9). Ultimately, we believe that complex traits demand analysis across multiple genomic scales and ancestries in order to gain biological insight into complex trait architecture and ultimately achieve personalized medicine.

As has been previously noted ^5, 25^, non-European ancestry cohorts are often excluded from GWA analyses of multi-ancestry biobanks; complementing the analyses of Ben-Eghan et al. ^24^, we find that 80.13% of UK Biobank studies over the last 9 years only report significant SNP-level associations in the white British cohort (Figure S1 - Figure S2), despite the tens of thousands of individuals of non-European ancestry sampled in that data set. Unless this practice is curbed by the biomedical research community, it will exacerbate already existing disparities in healthcare across diverse communities. There are undoubted benefits from increased sampling in a given ancestry for association mapping using the standard GWA framework, but it is still unknown the extent to which results from larger GWA and fine-mapping studies using European-ancestry genomes will generalize to the entire human population ^2, 20^.

Here, we have not addressed the downstream consequences of using self-identified ancestry to define cohorts in large-scale GWA studies (but see Urbut et al. ^37^, Willer et al. ^68^, Lin et al. ^69^, Yang et al. ^70^). Each sample we analyzed has also experienced environmental exposures that may influence the statistical detection of genetic associations, and some of those environmental exposures may be correlated with genomic ancestry ^13, 71–73^. Interrogation of the influence of gene by environment interactions on complex traits must be done with highly controlled experiments, which can in turn help prioritize traits in which association studies will be interpretable and useful. Increasing sample size in GWA studies alone will not resolve these fundamental biological questions: the proportion of phenotypic variance explained by associations discovered as sample sizes increase in GWA studies has largely reached diminishing returns ^39^, and gene by environment interactions are increasingly influential, and estimable, in large biobanks with cryptic relatedness ^74, 75^.

Many recent methodological advances that leverage GWA summary statistics have focused on: testing the co-localization of causal SNPs (e.g., fine mapping ^34, 54, 76, 77^); the non-additive effects of SNP-level interactions (i.e., epistasis ^78, 79^); and multivariate GWA tests ^37, 79–81^. While these methods can be extended and applied to multi-ancestry GWA analyses, they still focus on SNP-level signals of genetic trait architecture (see also Brown et al. ^82^, Galinsky et al. ^83^). Unlike the traditional GWA method, enrichment analyses increase statistical power by aggregating SNP-level signals of genetic associations and allowing for genetic heterogeneity in SNP-level trait architecture across samples, as well as offering the opportunity for immediate insights into trait architecture using existing datasets. However, these methods have been comparatively underused in multi-ancestry GWA studies.

While many studies note that differences in LD across ancestries affect transferability of effect size estimates ^6, 48, 84–86^, recent studies in population genetics have additionally debated how various selection pressures and genetic drift may hamper transferability of GWA results across ancestries (see for example, Edge and Rosenberg ^15, 16^, Novembre and Barton ^18^, Harpak and Przeworski ^20^, Durvasula and Lohmueller ^22^, Mostafavi et al. ^23^). Future GWA studies should be coupled with approaches from studies of how evolutionary processes shape the genetic architecture of complex traits ^20, 28, 50, 87^.

Two open questions must be tackled when studying complex trait architecture in the multi-ancestry biobank era: (*i*) to what extent is the true genetic trait architecture (causal SNPs and/or their effects on a trait of interest) heterogeneous across cohorts? ^6, 88^ and (*ii*) which components of GWA results (e.g. *p*-values, estimated effect size, direction of effect sizes) are transferable across ancestries, at any genomic scale? Continued application of the standard SNP-level GWA approach will not answer these questions. However, enrichment methods that aggregate SNP-level effects, test for effect size heterogeneity, leverage genomic annotations and gene interaction networks offer opportunities to directly test these fundamental questions. Methods can and should play an important role as biomedical research shifts current paradigms to extend the benefits of personalized medicine beyond people of European ancestry.

Additionally, biomedical researchers should continue to pressure both funding agencies and institutions to diversify their sampling efforts in the name of inclusion and addressing—instead of exacerbating—genomic health disparities. In addition to those efforts, we believe existing and new methods can increase the return on investment in multi-ancestry biobanks, ensure that every bit of information from these datasets is studied, and prioritize biological mechanism above SNP-level statistical association signals by identifying associations that are robust across ancestries.

## Appendices

Detailed information about the SNP-level results for both C-reactive protein and height, as well as, gene and pathway level associations for platelet count and triglyceride levels can be found in the Appendix.

## Declaration of interests

C.G. owns stock in 23andMe. E.E.K. and C.G. are members of the scientific advisory board for Encompass Bioscience. E.E.K. consults for Illumina.

## Acknowledgments

We thank Kirk Lohmueller and Alicia R. Martin for helpful comments on an earlier version of this manuscript, as well as the Crawford and Ramachandran Labs for helpful discussions. This research was conducted in part using computational resources and services at the Center for Computation and Visualization at Brown University as well as, using the UK Biobank Resource under Application Number 22419. The Population Architecture Using Genomics and Epidemiology (PAGE) program is funded by the National Human Genome Research Institute (NHGRI) with co-funding from the National Institute on Minority Health and Health Dis-parities (NIMHD). The WHI program is funded by the National Heart, Lung, and Blood Institute, National Institutes of Health, U.S. Department of Health and Human Services through contracts 75N92021D00001, 75N92021D00002, 75N92021D00003, 75N92021D00004, 75N92021D00005. The HCHS/SOL study was carried out as a collaborative study supported by contracts from the National Heart, Lung and Blood Institute (NHLBI) to the University of North Carolina (N01-HC65233), University of Miami (N01-HC65234), Albert Einstein College of Medicine (N01-HC65235), Northwestern University (N01-HC65236) and San Diego State University (N01-HC65237). S.P.S. is a trainee supported under the Brown University Predoctoral Training Program in Biological Data Science (NIH T32 GM128596). L.C. acknowledges the support of an Alfred P. Sloan Research Fellowship and a David & Lucile Packard Fellowship for Science and Engineering. This work was also supported by US National Institutes of Health R01 GM118652 to S.R., and S.R. acknowledges additional support from National Science Foundation CAREER Award DBI-1452622.

## Web Resources

The methods applied in this paper include: PESCA (https://github.com/huwenboshi/pesca), RSS(https://github.com/stephenslab/rss), gene-*ε* (https://github.com/ramachandran-lab/genee), and Hierarchical HotNet (https://github.com/raphael-group/hierarchical-hotnet).

## Data and code availability

All scripts, publicly available data, and outputs from GWA, gene, and pathway association tests are available at https://github.com/smithsap/redefining_replication. Results from PESCA analyses were provided through personal correspondence with Huwenbo Shi.

## Supplemental Figures

**Figure S1:**
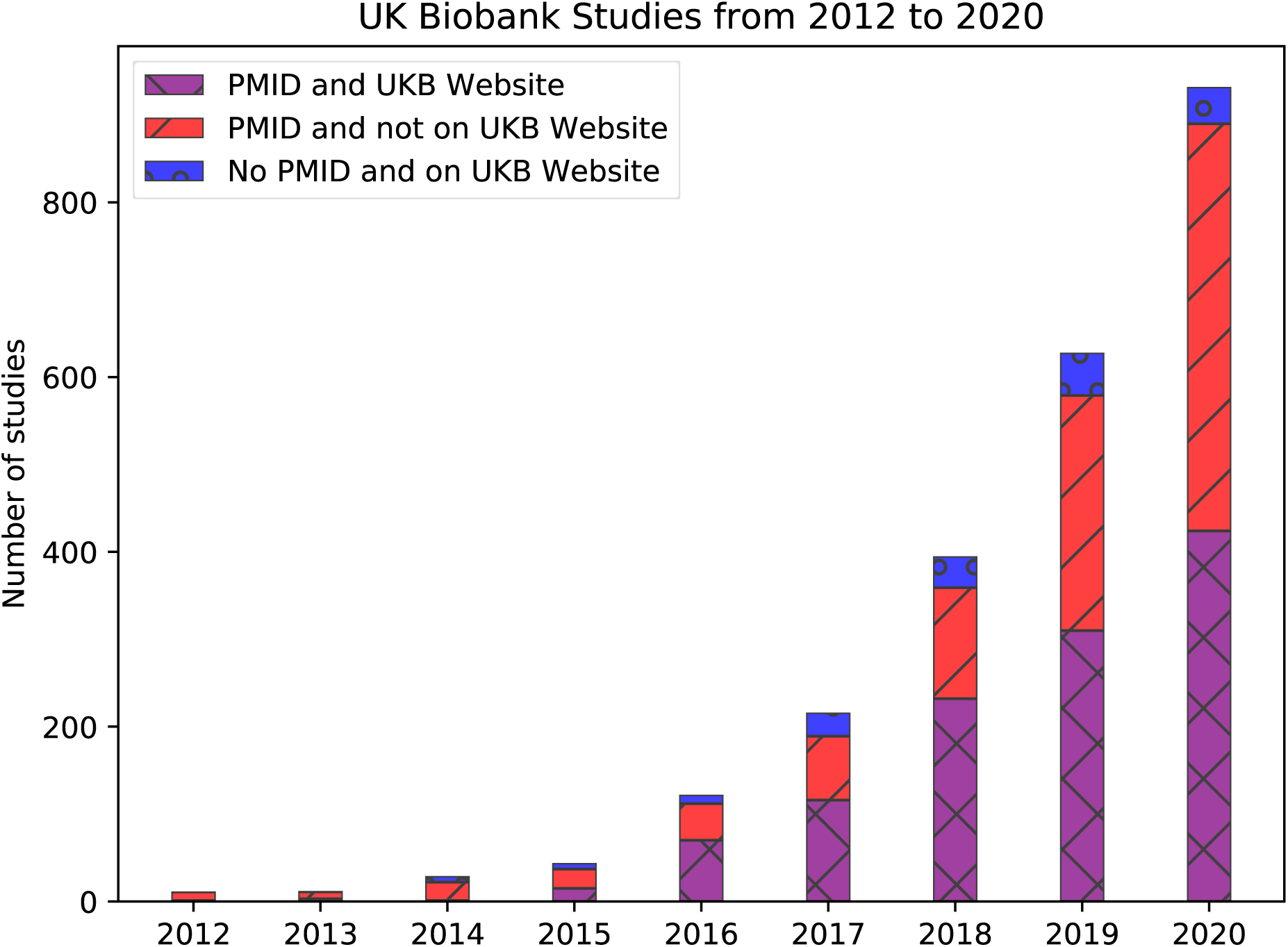
Number of publications we identified using UK Biobank data from 2012 to 2020. Studies identified using PMIDs as described in the Supplemental Information. Studies that are displayed on the UK Biobank website (https://www.ukbiobank.ac.uk/) and identified on PubMed are shown in purple. Studies listed on the UK Biobank website but do not have a PMID are shown in blue, and studies only identified using PubMed but not listed on the UK Biobank website are shown in red. The protocols for identifying studies both on PubMed and the UK Biobank website are detailed in the Supplemental Information. Data from both the UK Biobank website and PubMed were accessed on January 12, 2021.

**Figure S2:**
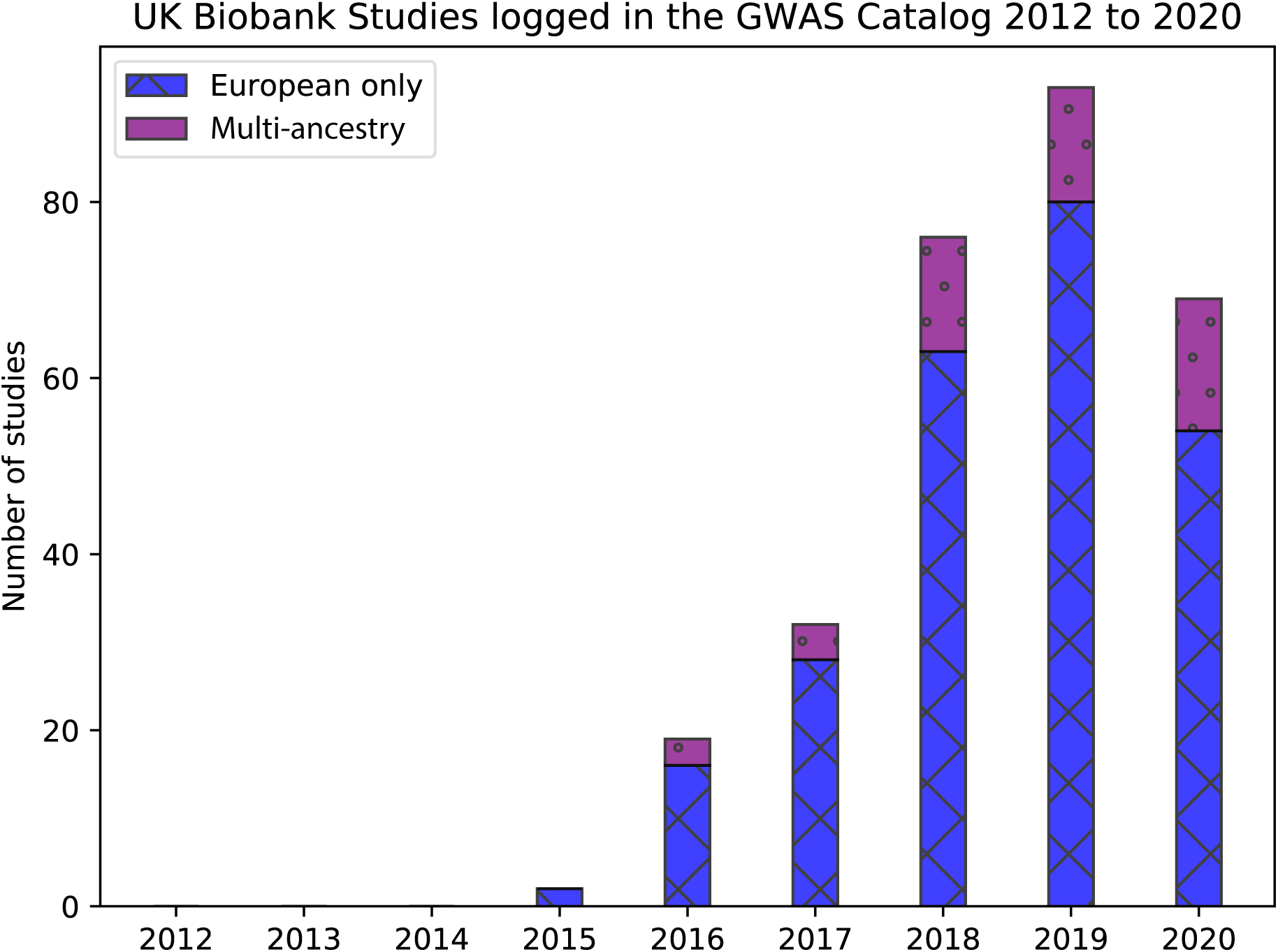
Number of studies published using UK Biobank data from 2012 to 2020 that have available metadata in the GWAS Catalog. Our protocols for identifying studies from the GWAS Catalog are detailed in the Supplemental Information. Multi-ancestry studies are shown in purple and include those that list samples of more than one ancestral group in the GWAS catalog (as defined according to the protocol using Popejoy and Fullerton ^25^, available on the GitHub page https://github.com/ramachandran-lab/redefining_replication). Studies that only list samples of European ancestry in the GWAS catalog are shown in blue. Every multi-ancestry analysis includes samples of European ancestry and of at least one other ancestry. GWAS Catalog data was accessed on January 10, 2021 from the website https://www.ebi.ac.uk/gwas/docs/file-downloads using the final release file of 2020 (see file named gwas catalog v1.0.2-associations e100 r2020-12-15.tsv).

**Figure S3:**
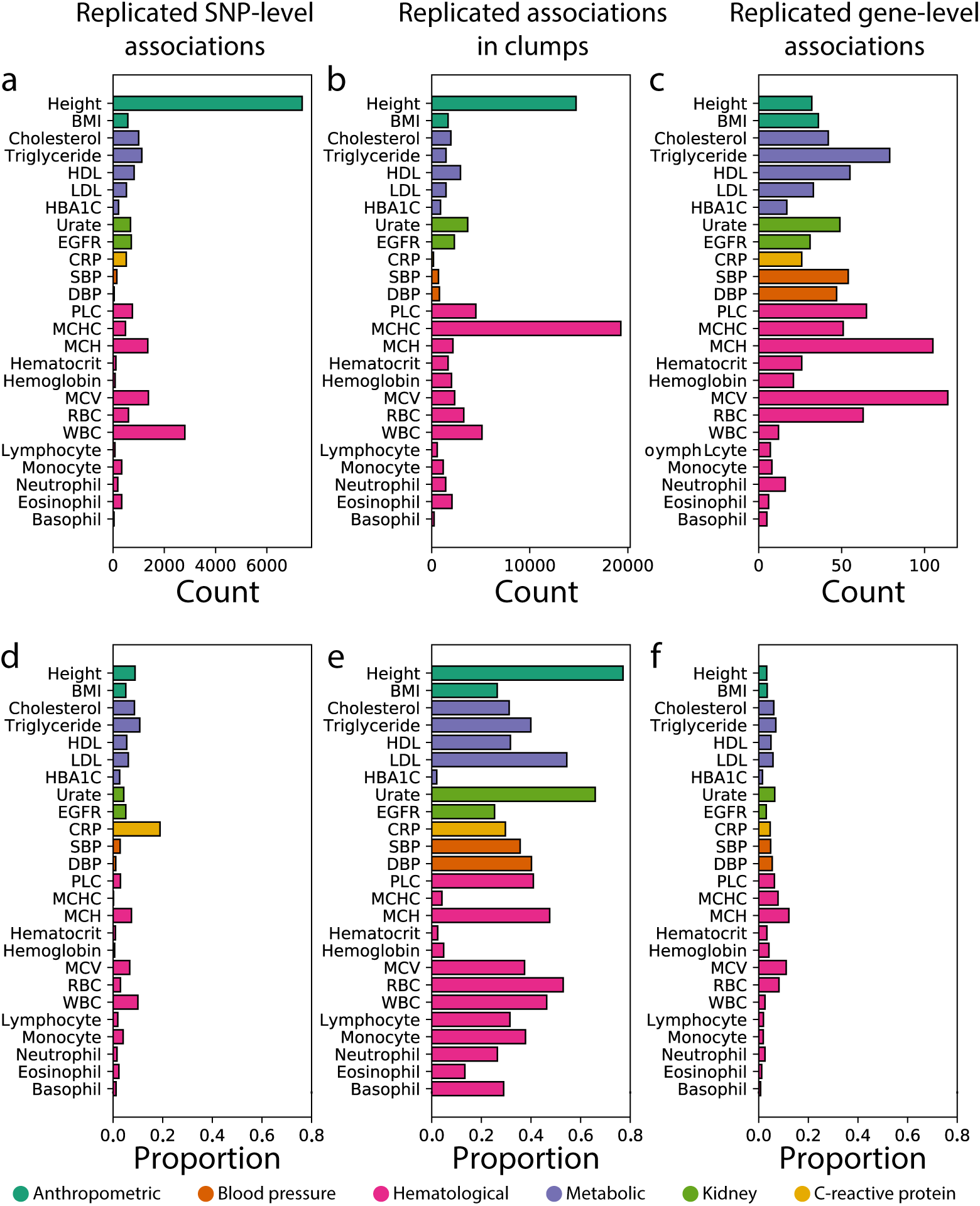
Summaries of replicated associations at multiple genomic scales among ancestry cohorts for all 25 traits analyzed using Bonferroni-corrected thresholds. Expansion of three letter trait codes are given in Table S2. (**a**) Number and (**d**) proportion of genome-wide significant SNPs associated with a phenotype in at least one ancestry cohort that were replicated in at least two ancestry cohorts. In all 25 traits, genome-wide significant SNPs replicate in at least two ancestry cohorts. Height contains over 7,000 replicated SNPs among the seven ancestry cohorts analyzed, illustrating its highly polygenic architecture. For many traits across all categories, with the exception of other biochemical (i.e., CRP), the replication rate of genes is higher in gene-level associations than at the SNP-level. (**b**) Number and (**e**) proportion of 1Mb windows, or “clumps”, that contain at least one genome-wide significant SNP-level associations for a given phenotype in at least two ancestry cohorts. (**c**) Number and (**f**) proportion of genome-wide significant gene-level associations that replicate among ancestry cohorts. Replicated associations in hematological are common at the gene-level in hematological and metabolic traits. For instance, in three of the four cohorts with mean corpuscular hemoglobin (MCH) measurements *HBA1* and *HBA2* were identified as significant associated with MCH in the African, European, and East Asian ancestry cohorts Table S3. The denominator of the proportion is calculated as the total number of unique SNPs, clumps, or genes that are significantly associated with a trait in at least on ancestral cohort. Note that **d** and **f** correspond to Figure 1**a** and **b**, with an altered x-axis upper limit of 0.8 in this figure.

**Figure S4:**
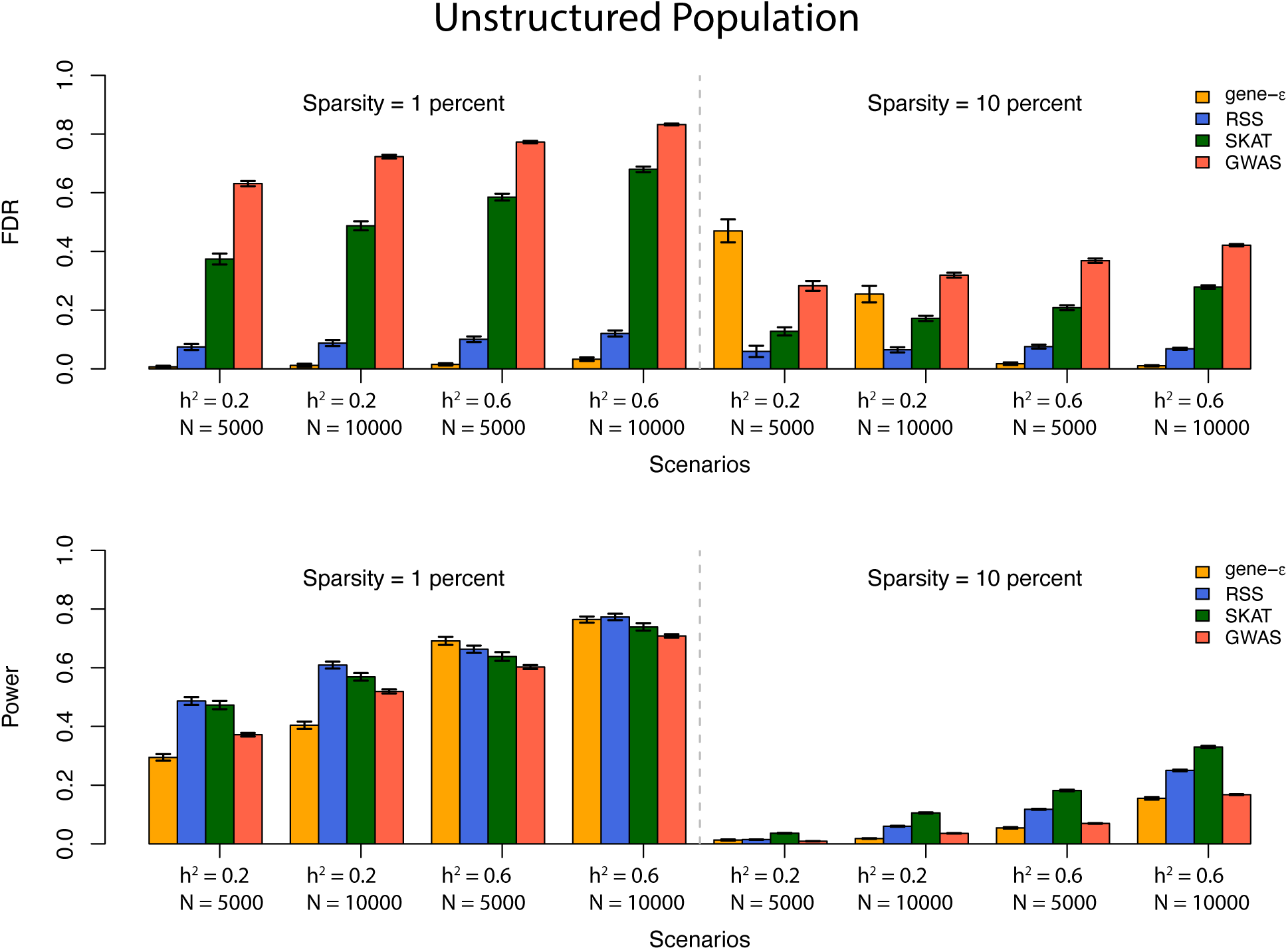
gene-*ε* outperforms and controls false discovery rate (FDR) better than other association methods in simulations with varying heritability and sample size. Simulations were designed to assess gene versus SNP-level association false discovery rate (FDR) and power in an unstructured population as described by the protocols in the Supplemental Information. The top and bottom panels show the FDR and power of four different association methods on 100 simulated datasets, respectively. We compared performance of three gene-level association test methods (gene-*ε*^38^, RSS ^60^, SKAT^40^) with outputs from the standard GWA association test under different simulation parameters (sample size *N*, narrow-sense heritability *h*^2^, and sparsity). We define sparsity as the proportion of SNPs that are ground-truth causal. Standard errors across the simulated replicates are shown using black whisker plots. Simulation protocol is described in the Supplemental Information.

**Figure S5:**
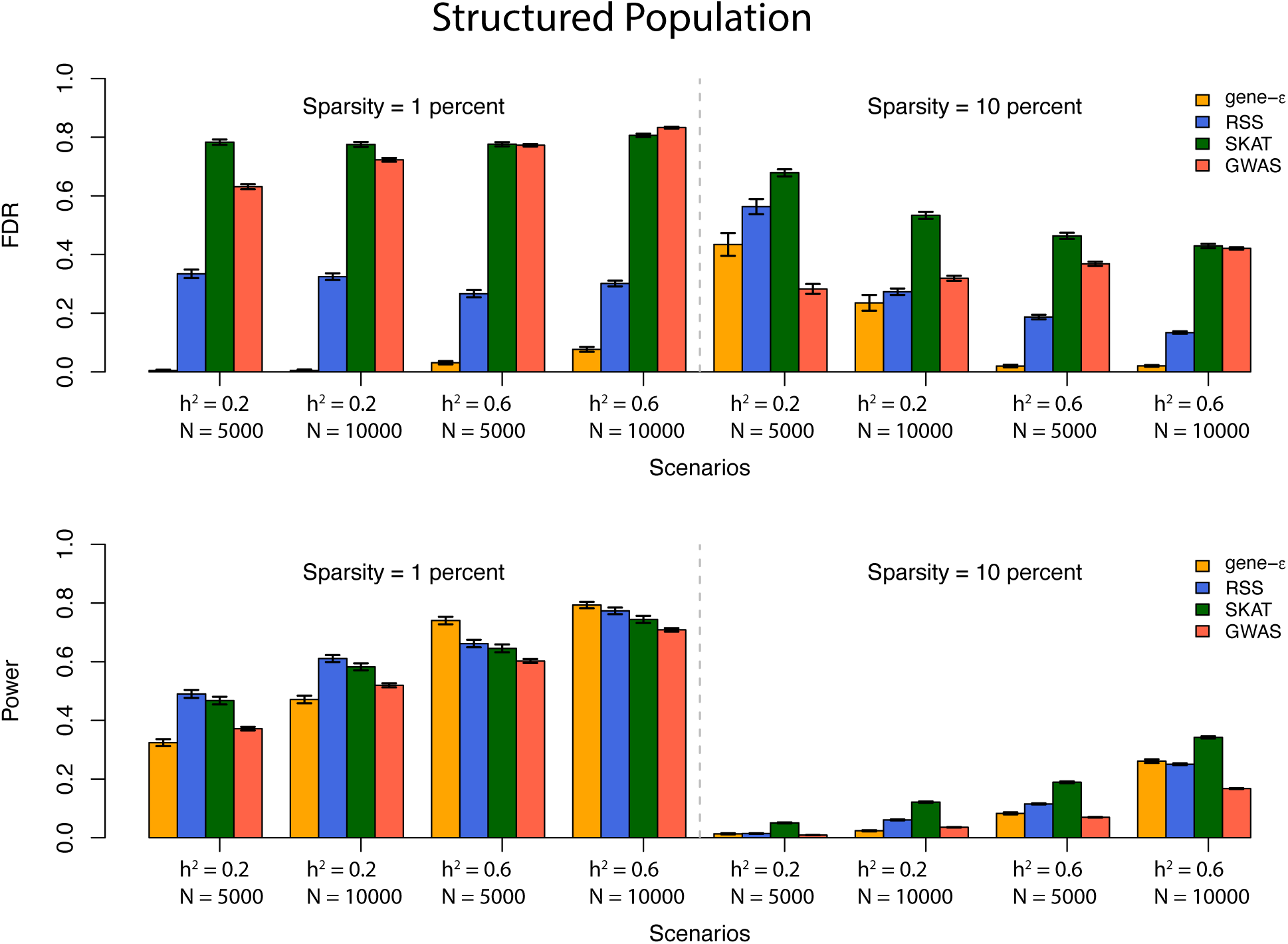
gene-*ε* outperforms and controls false discovery rate (FDR) better than other association methods in simulations with varying heritability and sample size. Simulations are designed to assess gene versus SNP-level association false discovery rate (FDR) and power in an structured population as described by the protocols in the Supplemental Information. The top and bottom panels show the FDR and power of four different association methods on 100 simulated datasets, respectively. We compared performance of three gene-level association test methods (gene-*ε*^38^, RSS ^60^, SKAT^40^) with outputs from the standard GWA association test under different simulation parameters (sample size *N*, narrow-sense heritability *h*^2^, and sparsity). We define sparsity as the proportion of SNPs that are designated to be causal. Standard errors across the simulated replicates are shown using black whisker plots.

**Figure S6:**
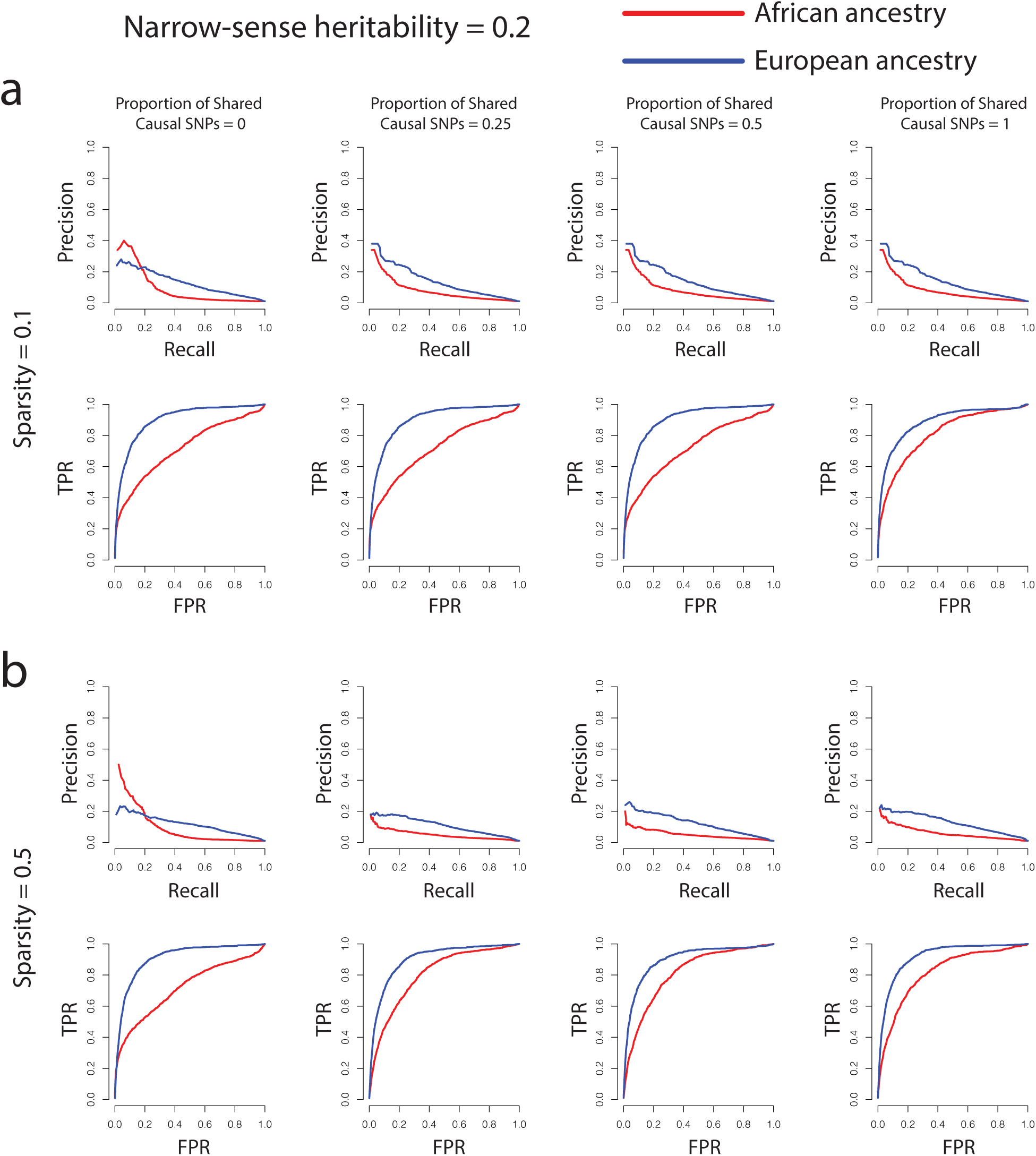
gene-*ε* identifies associated genes in two simulated ancestry cohorts under a variety of genetic architectures with low narrow sense heritability. **a.** Precision-recall (top row) and receiver operating curves (bottom row) for gene-*ε* analysis of cohorts simulated using genotypes from individuals of European (*N* = 10,000; blue line) and African (*N* = 4,967; red line) ancestry, respectively. Narrow-sense heritability was set to *h*^2^ = 0.2 in each simulation. Sparsity of causal SNPs was set to a proportion of 0.1 and the proportion of causal SNPs shared was tested at different values. 50 replicates of each set of simulations under each parameter were performed. **b.** Precision-recall (top row) and receiver operating curves (bottom row) for gene-*ε* analysis of 50 replicated simulations of a European and African cohort using a causal SNP sparsity of 0.5.

**Figure S7:**
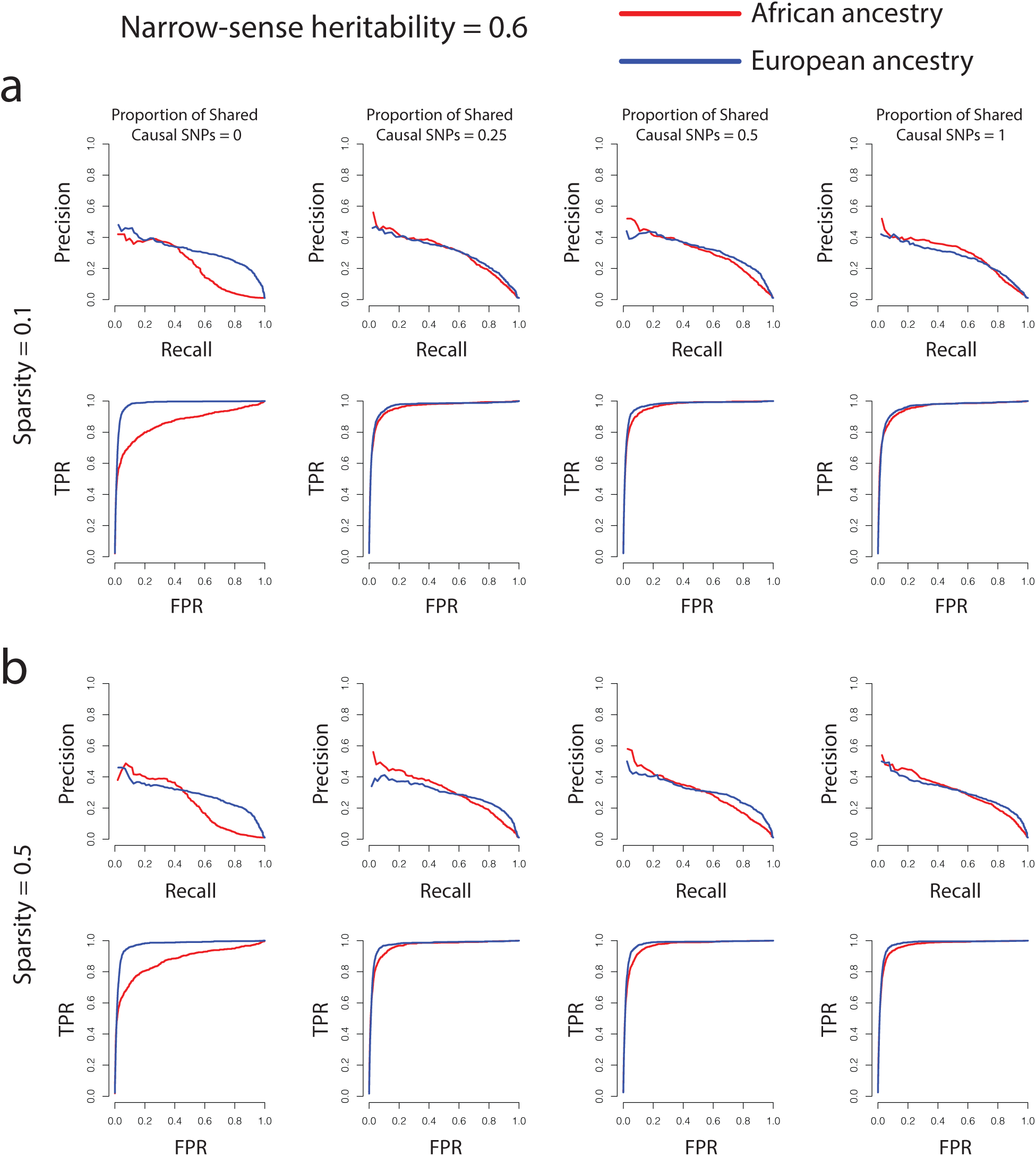
gene-*ε* identifies associated genes in two simulated ancestry cohorts under a variety of genetic architectures with high narrow sense heritability. **a.** Precision-recall (top row) and receiver operating curves (bottom row) for gene-*ε* analysis of cohorts simulated using genotypes from individuals of European (*N* = 10,000; blue line) and African (*N* = 4,967; red line) ancestry, respectively. Narrow-sense heritability was set to *h*^2^ = 0.6 in each simulation. Sparsity of causal SNPs was set to a proportion of 0.1 and the proportion of causal SNPs shared was tested at different values. 50 replicates of each set of simulations under each parameter were performed. **b.** Precision-recall (top row) and receiver operating curves (bottom row) for gene-*ε* analysis of 50 replicated simulations of a European and African cohort using a causal SNP sparsity of 0.5.

**Figure S8:**
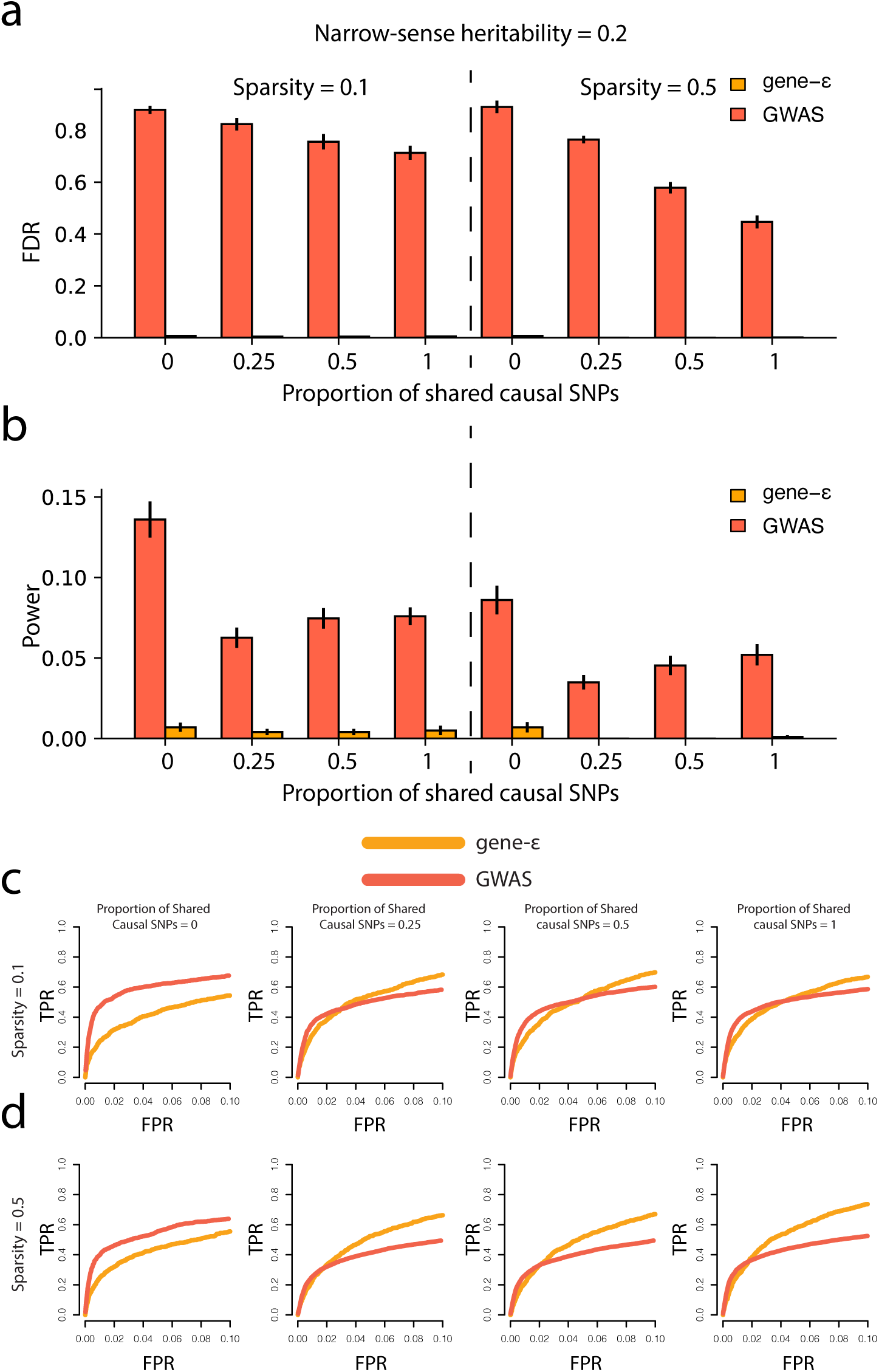
gene-*ε* (orange) has a lower false discovery rate for identification of shared genetic determinants between cohorts than the standard GWA framework (red). Narrow-sense heritability (percent variance explained by the genotype matrix) was set to *h*^2^ = 0.2 for all simulations. **a.** False discovery rate of shared genetic determinants between two ancestry cohorts using varying levels of causal SNP sparsity and proportion of shared causal SNPs between the cohorts. **b.** Power of gene-*ε* and the standard GWA framework to detect shared genetic determinants between two cohorts. Error bars were calculated using the results from 50 simulations of each parameter set of sparsity and proportion of shared causal SNPs for both FDR(a) and Power(b). **c.** Receiver operating curves corresponding to simulations of genetic architecture when causal SNP sparsity is equal to 0.1 (corresponding to the left-hand panels of **a** and **b**). **d.** Receiver operating curves corresponding t2o8 simulations of genetic architecture when causal SNP

**Figure S9:**
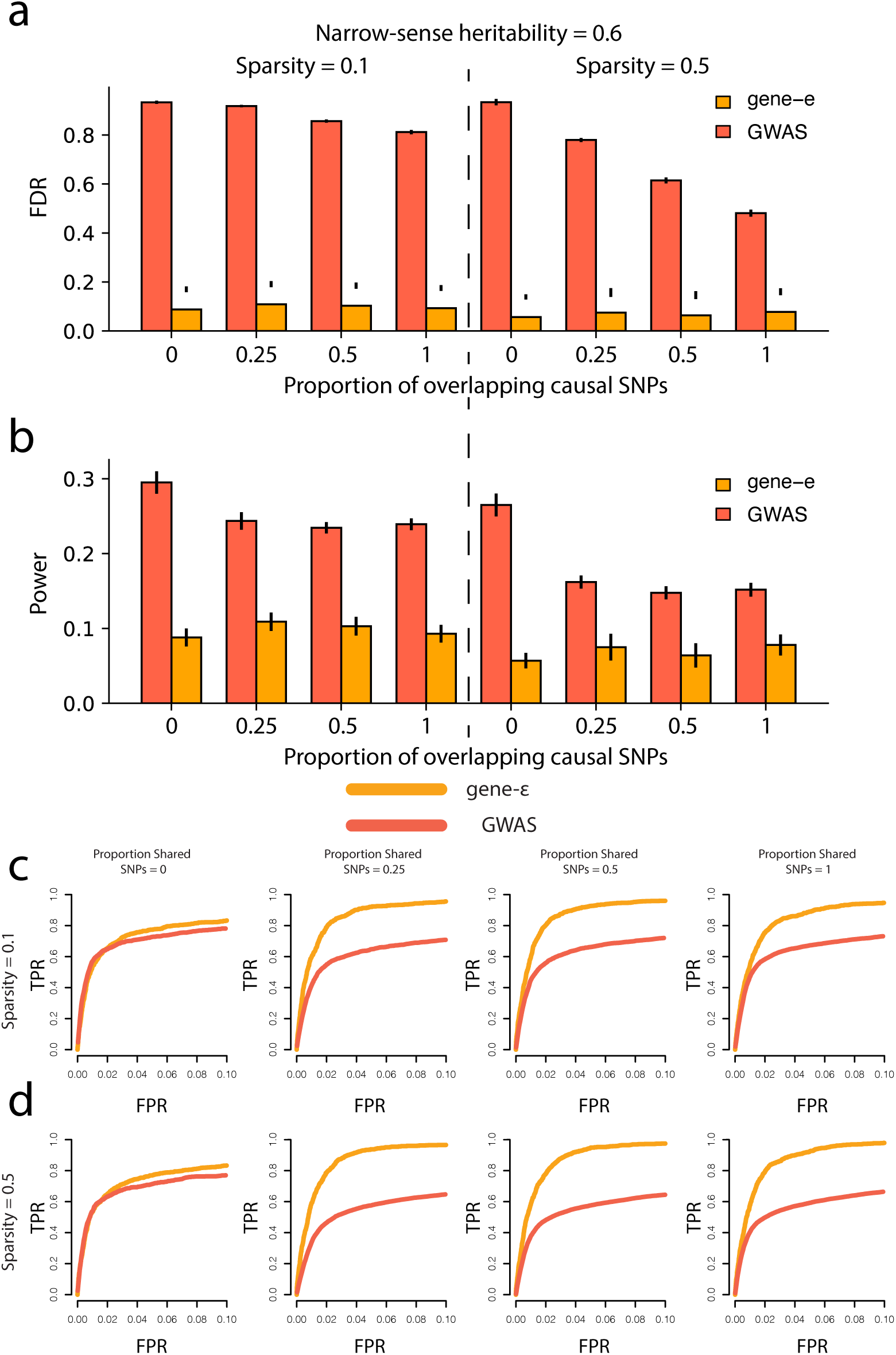
gene-*ε* (orange) has a lower false discovery rate for identification of shared genetic determinants between cohorts than the standard GWA framework (red). Narrow-sense heritability (percent variance explained by the genotype matrix) was set to *h*^2^ = 0.6 for all simulations. **a.** False discovery rate of shared genetic determinants between two ancestry cohorts using varying levels of causal SNP sparsity and proportion of shared causal SNPs between the cohorts. **b.** Power of gene-*ε* and the standard GWA framework to detect shared genetic determinants between two cohorts. Error bars were calculated using the results from 50 simulations of each parameter set of sparsity and proportion of shared causal SNPs for both FDR(a) and Power(b). **c.** Receiver operating curves corresponding to simulations of genetic architecture when causal SNP sparsity is equal to 0.1 (corresponding to the left-hand panels of **a** and **b**). **d.** Receiver operating curves corresponding to simulations of genetic architecture when causal SNP

**Figure S10:**
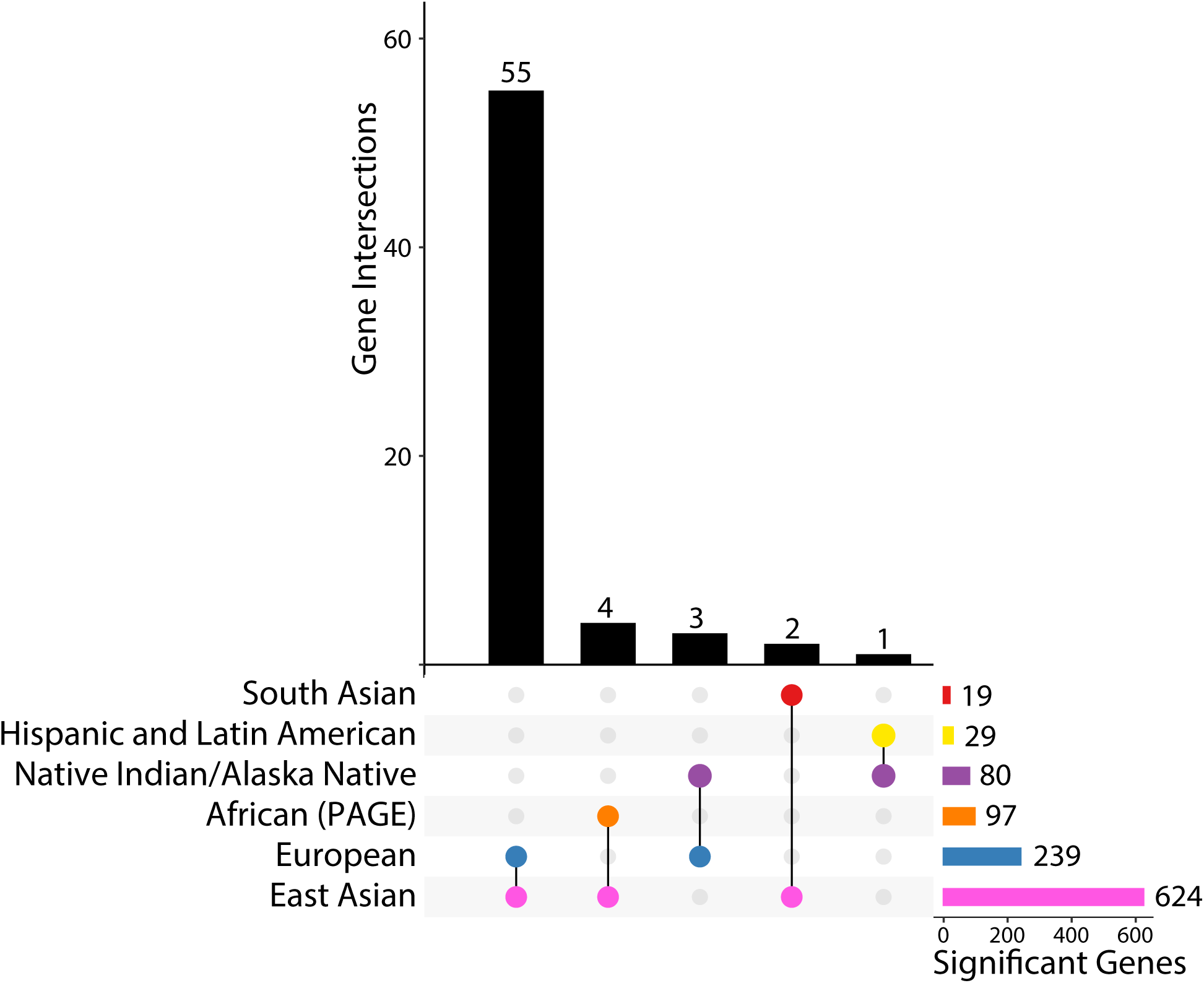
All six ancestries have gene-level associations with platelet count that replicate in at least one other ancestry. Total number of genome-wide significant genes in each ancestry, after correcting for total number of regions tested, are given in the bar plot located in the bottom right (significance thresholds are given in Table S15, sample sizes are given in Table S5 - Table S8). Shared gene-level association statistics between pairs of ancestries are shown in the vertical bar plot; the pair of ancestries represented by each bar can be identified using the dots and links below the barplot. Of the 65 genome-wide significant gene-level association statistics that replicate in at least two ancestry cohorts, only 25 contain SNPs that have been previously associated with platelet count in at least one ancestry in at least one study in the GWAS catalog (https://www.ebi.ac.uk/gwas/home) This plot was generated using the UpSetR package ^90^.

**Figure S11:**
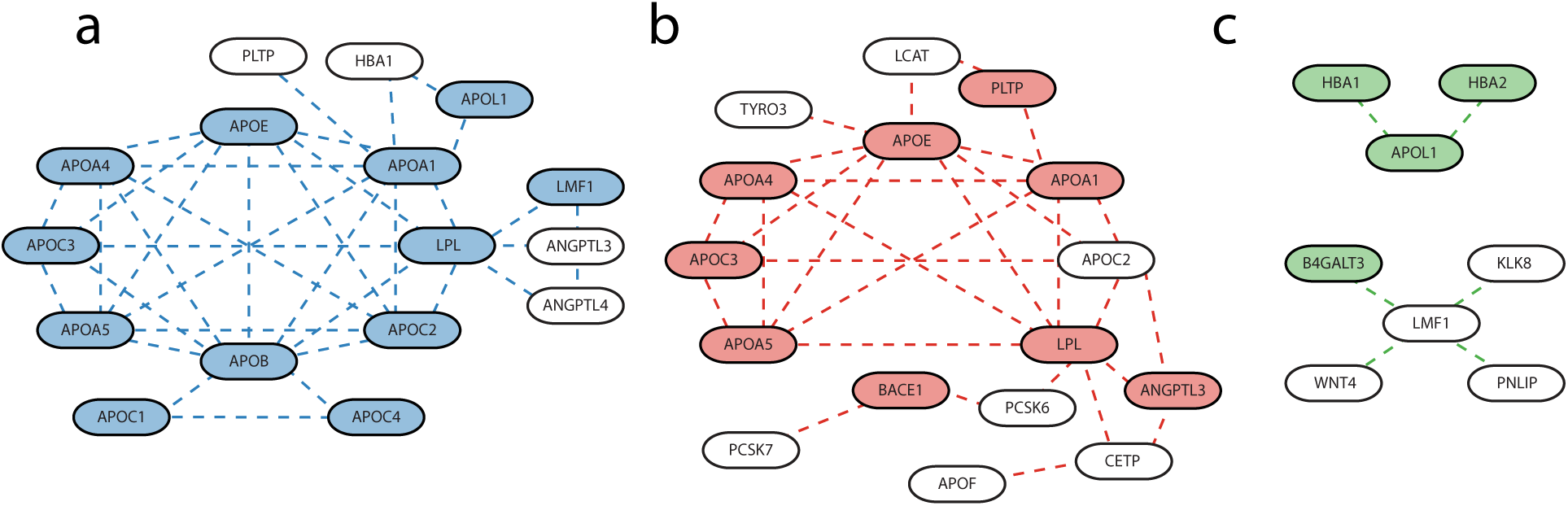
Significantly mutated subnetworks associated with triglyceride levels identified in the (a) European, (b) East Asian, and (c) Native Hawaiian ancestry cohorts. Significantly mutated subnetworks were identified using the Hierarchical HotNet method ^31^. Genes that were identified in each ancestry as significantly associated with triglyceride levels using the gene-*ε* method are shaded using ancestry-specific color coding (also used in Figure 3, European—blue, East Asian—pink, Native Hawaiian— green). Significantly mutated subnetworks in the (**a**) European and (**b**) East Asian cohorts were identified using the ReactomeFI ^44^ protein-protein interaction network, and the significantly mutated subnetwork in the (**c**) the Native Hawaiian cohort was identified using the iRefIndex 15.0^45^ protein-protein interaction networks. Genes that are present in any of the significantly mutated subnetworks that contain SNPs previously associated with triglyceride levels in the GWAS Catalog are listed with corresponding citations in Table S16.

**Figure S12:**
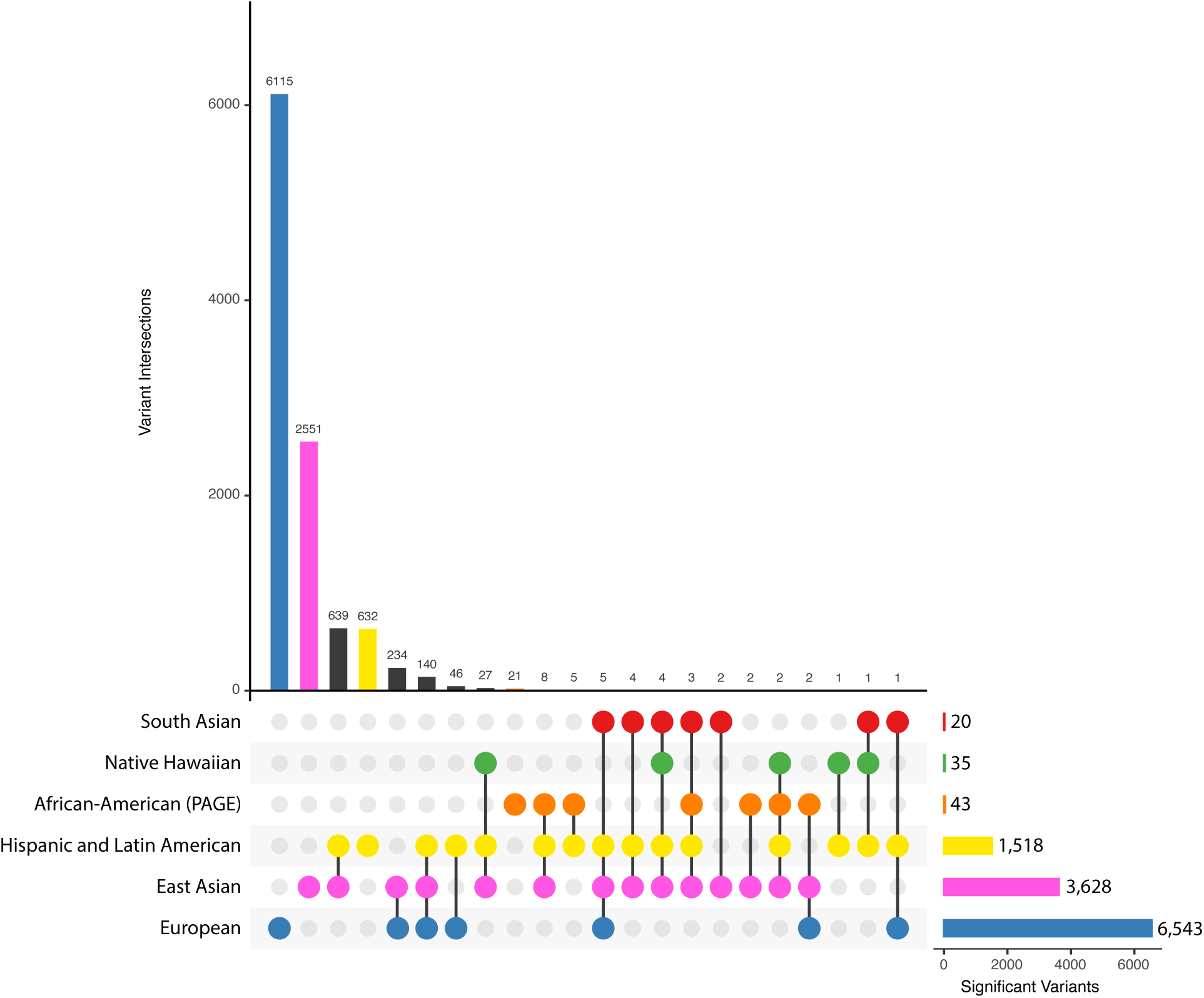
Shared SNP-level associations with triglyceride levels in six ancestral cohorts. Total number of genome-wide significant genes in each ancestry, after correcting for total number of regions tested, are given in the bar plot located in the bottom right (significance thresholds are given in Table S10 and sample sizes are given in Table S5 - Table S8). Shared SNP-level association statistics between pairs of ancestries are shown in the vertical bar plot. The pair of ancestries represented by each bar can be identified using the dots and links below the vertical barplot. This plot was generated using the UpSetR package ^90^.

**Figure S13:**
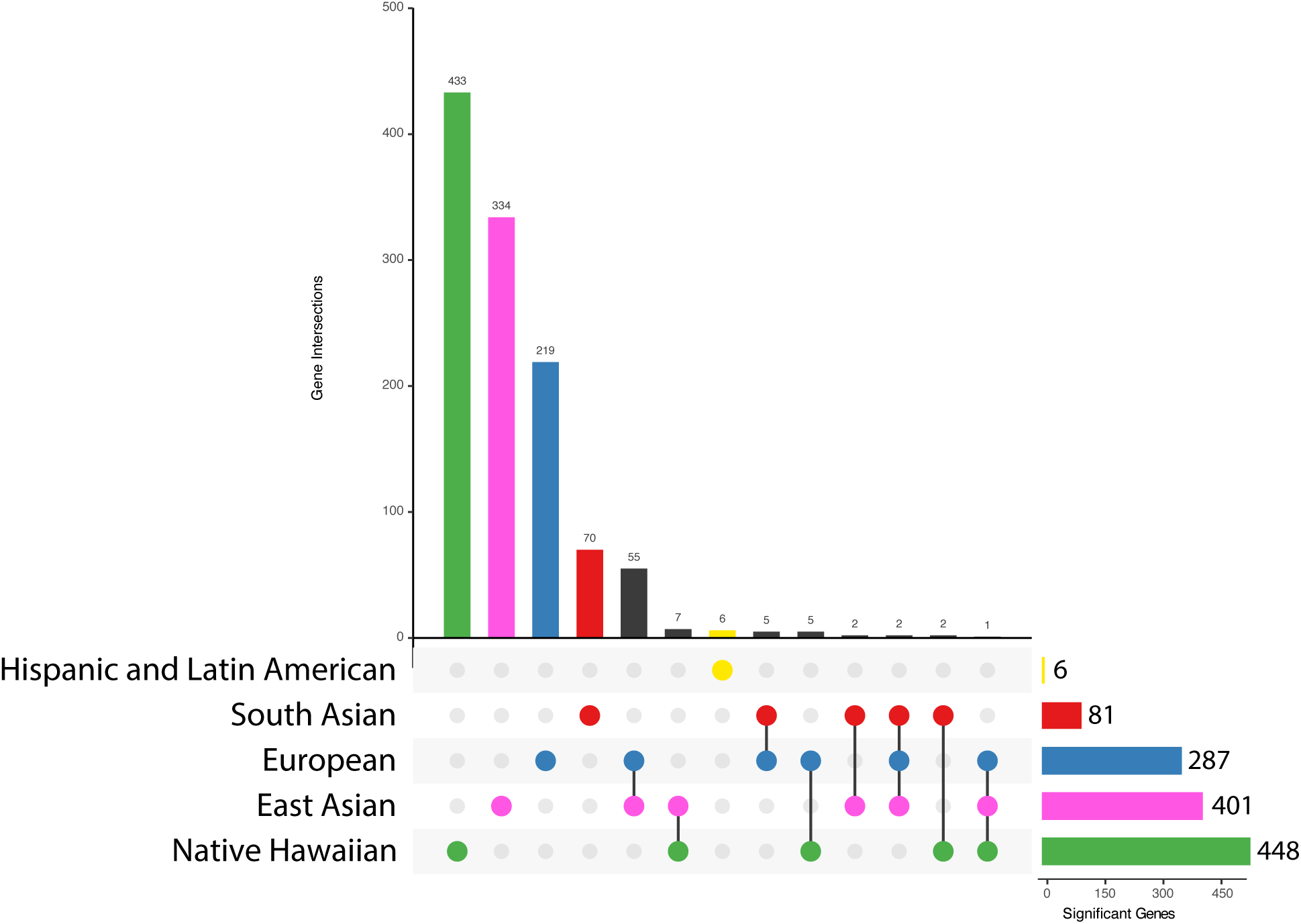
Shared gene-level associations with triglyceride levels in five ancestral cohorts. Total number of genome-wide significant genes in each ancestry, after correcting for total number of regions tested, are given in the bar plot located in the bottom right (significance thresholds are given in Table S15 and sample sizes are given in Table S5 - Table S8). Shared gene-level association statistics between pairs of ancestries are shown in the vertical bar plot. The pair of ancestries represented by each bar can be identified using the dots and links below the vertical barplot. This plot was generated using the UpSetR package ^90^.

**Figure S14:**
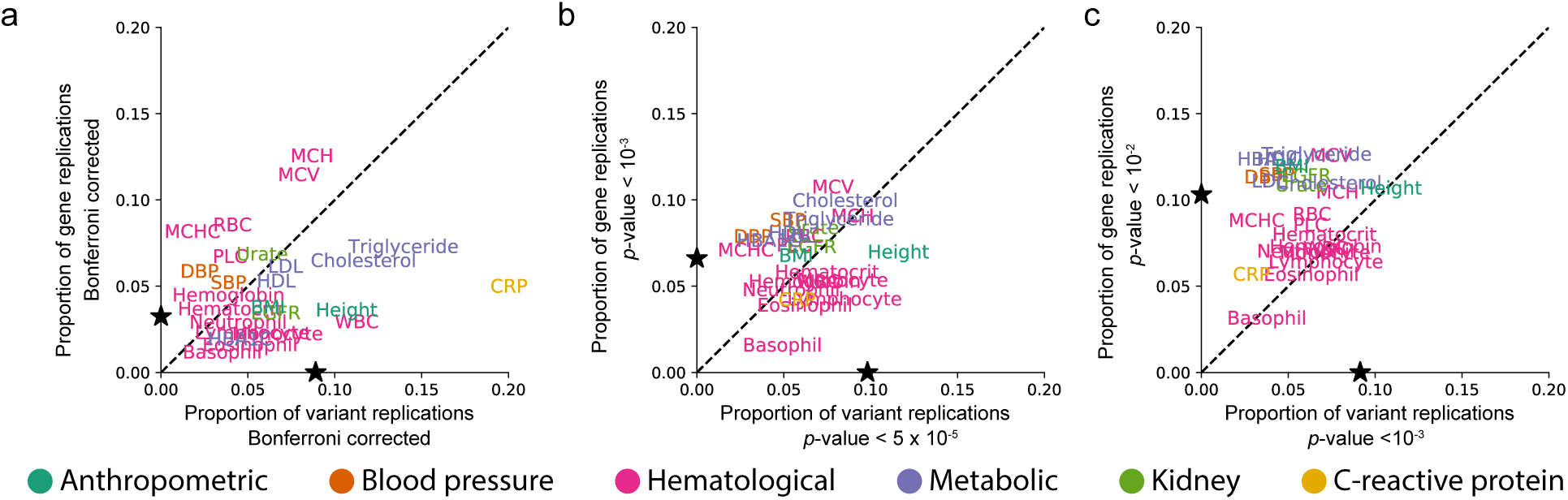
Less stringent significance thresholds lead to a decrease in the proportion of replicated SNP-level associations and an increase in the proportion of gene-level associations among ancestries for each of the 25 traits analyzed. **a.** Proportion of all SNP-level Bonferroni-corrected genome-wide significant associations in any ancestry that replicate in at least one other ancestry is shown on the x-axis (see Table S10 for ancestry-trait specific Bonferroni corrected *p*-value thresholds). On the y-axis we show the proportion of significant gene-level associations that were replicated for a given phenotype in at least two ancestries (see Table S15 for Bonferroni corrected significance thresholds for each ancestry-trait pair). The black stars on the x- and y-axes represent the mean proportion of replicates in SNP and gene analyses, respectively. C-reactive protein (CRP) contains the greatest proportion of replicated SNP-level associations of any of the phenotypes. **b.** The x-axis indicates the proportion of SNP-level associations that surpass a nominal threshold of *p*-value *<* 10*^−^*^5^ in at least one ancestry cohort that replicate in at least one other ancestry cohort. The y-axis indicates the proportion of gene-level associations that surpass a nominal threshold of *p*-value *<* 10*^−^*^3^ in at least one ancestry cohort and replicate in at least one other ancestry cohort. Nominal *p*-value thresholds tend to decrease the proportion of replicated SNP-level associations and tend to increase the proportion of replicated gene-level associations. The number of unique SNPs and genes that replicated in each cohort is given in Figure S15. **c.** The x-axis indicates the proportion of SNP-level associations that surpass a nominal threshold of *p*-value *<* 10*^−^*^3^ in at least one ancestry cohort that replicate in at least one other ancestry cohort. The y-axis indicates the proportion of gene-level associations that surpass a nominal threshold of *p*-value *<* 10*^−^*^2^ in at least one ancestry cohort and replicate in at least one other ancestry cohort. The number of unique SNPs and genes that replicated in each cohort is given in Figure S16. As shown in panel **b**, nominal *p*-value thresholds tend to decrease the proportion of replicated SNP-level associations and tend to increase the proportion of replicated gene-level associations. Expansion of three letter trait codes are given in Table S2.

**Figure S15:**
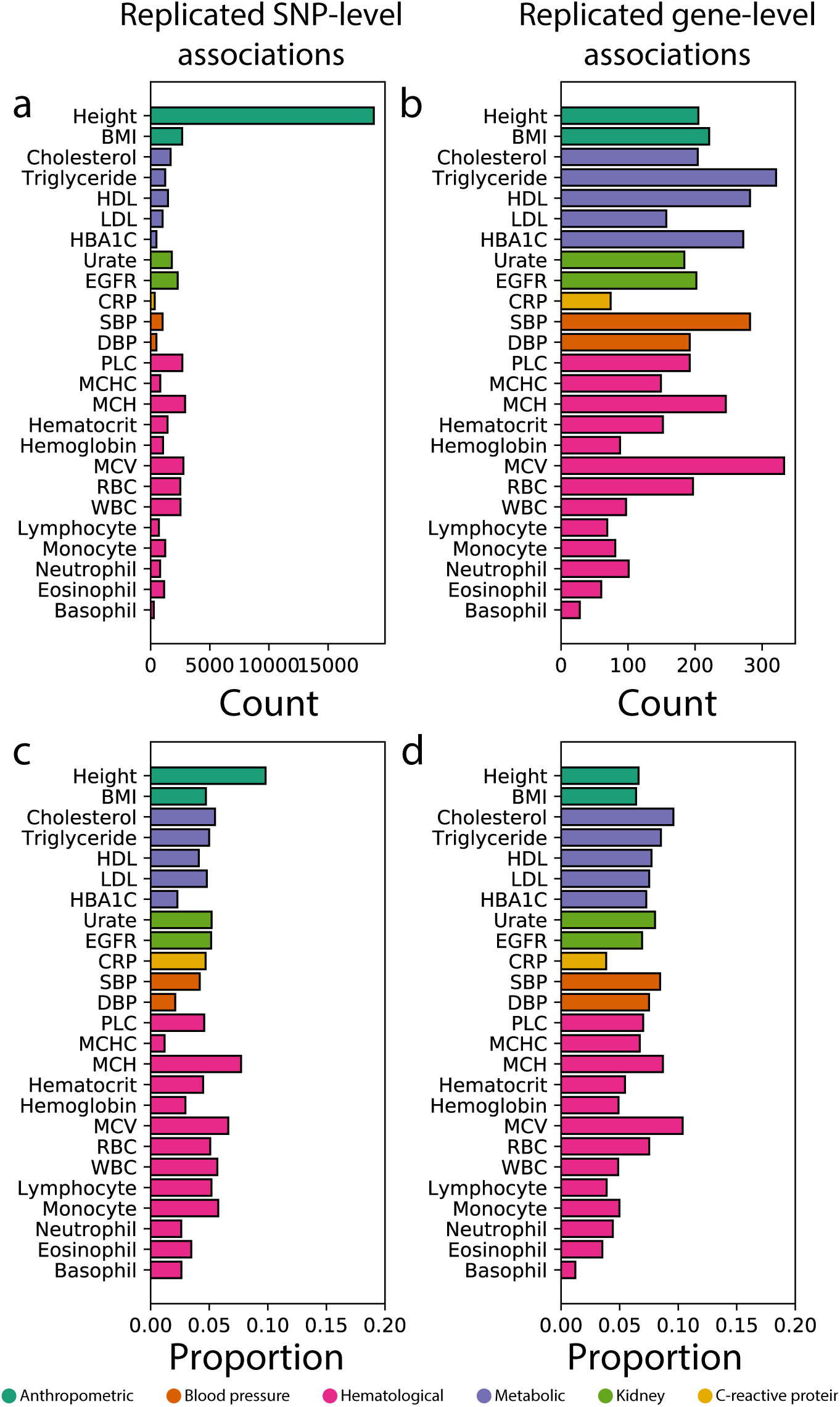
Summaries of replicated associations at multiple genomic scales among ancestry cohorts for all 25 traits analyzed using nominal *p*-value thresholds. (**a**) Number and (**c**) proportion of genome-wide significant SNPs associated with a phenotype in at least one ancestry cohort that were replicated in at least two ancestry cohorts using a nominal *p*-value threshold of 5 10*^−^*^5^. (**b**) Number and (**d**) proportion of genome-wide significant gene-level associations that replicate among ancestry cohorts using a nominal *p*-value threshold of 10*^−^*^3^. Expansion of three letter trait codes are given in Table S2.

**Figure S16:**
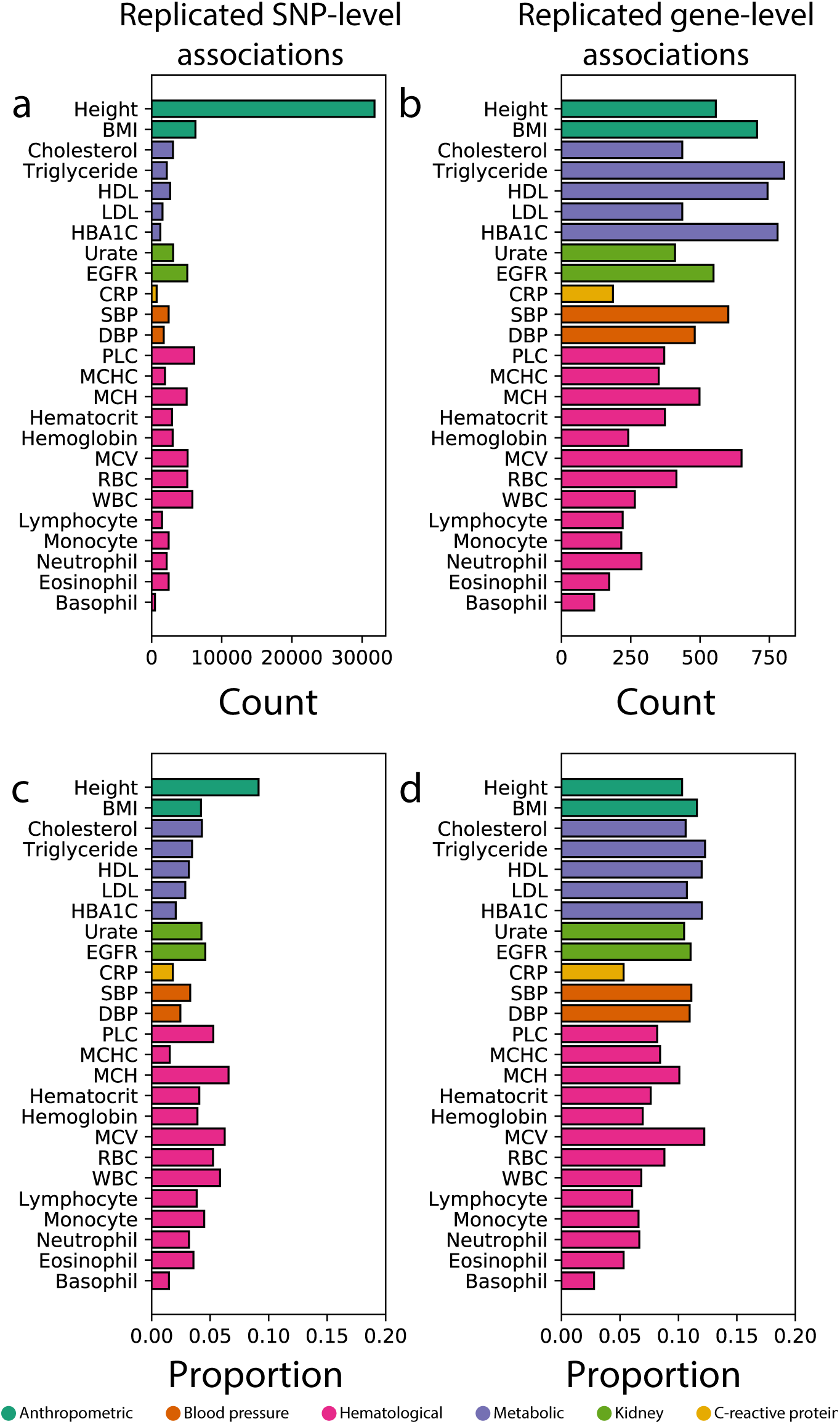
Summaries of replicated associations at multiple genomic scales among ancestry cohorts for all 25 traits analyzed using nominal *p*-value thresholds. (**a**) Number and (**c**) proportion of genome-wide significant SNPs associated with a phenotype in at least one ancestry cohort that were replicated in at least two ancestry cohorts using a nominal *p*-value threshold of 10*^−^*^3^. (**b**) Number and (**d**) proportion of genome-wide significant gene-level associations that replicate among ancestry cohorts using a nominal *p*-value threshold of 10*^−^*^2^. Expansion of three letter trait codes are given in Table S2. Expansion of three letter trait codes are given in Table S2.

**Figure S17:**
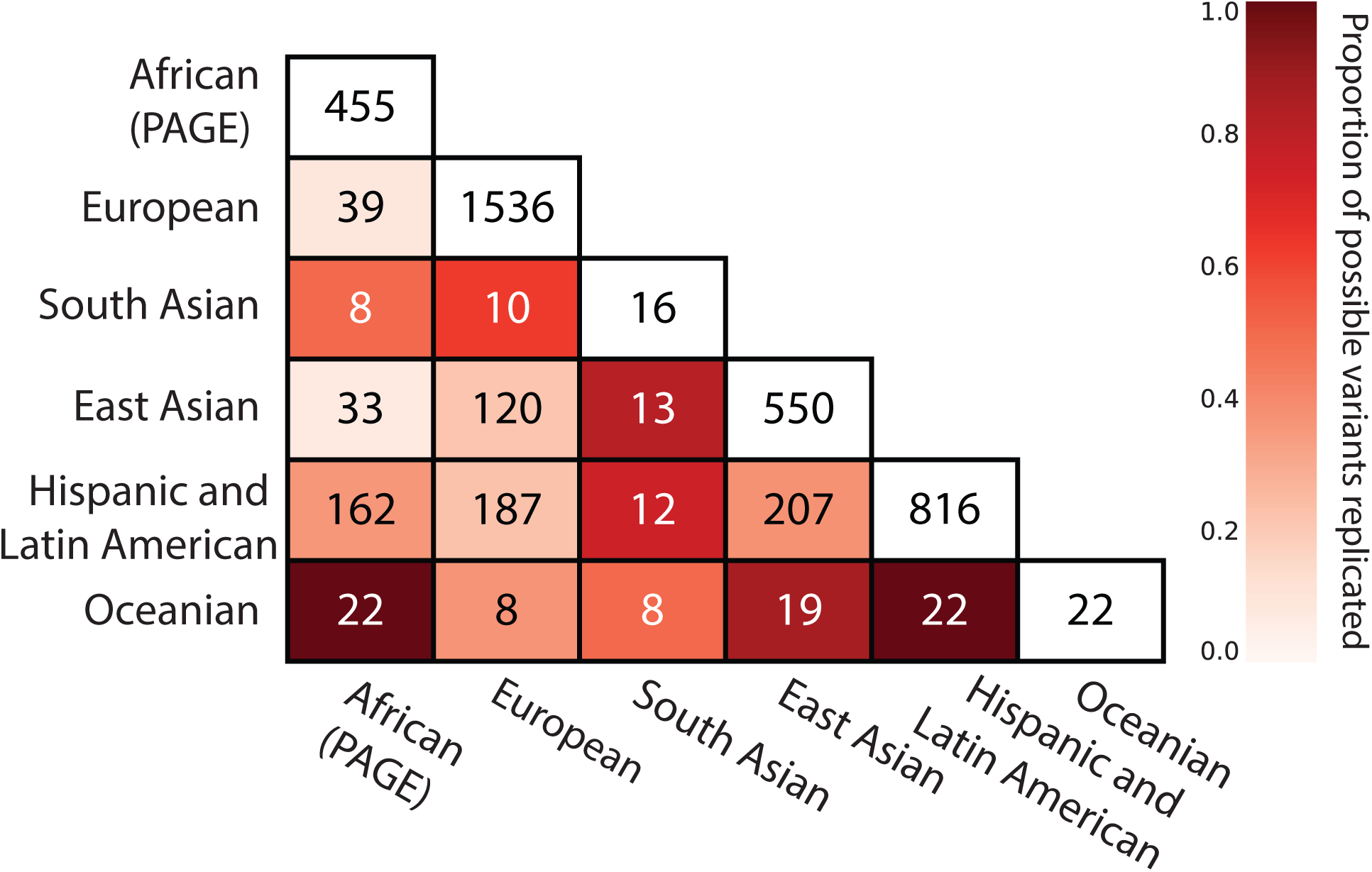
Pairwise replication of SNP-level association signals for C-reactive protein in six ancestral cohorts using genotype and imputed data. Imputed data was available and included in this analysis for each of the six cohorts. The inclusion of imputed SNPs in GWA analysis of C-reactive protein increases both the number of significant SNPs in each ancestry (along the diagonal) as well as the number of replicating significant SNP-level associations among pairs of ancestry cohorts (lower triangular). The corresponding analysis of pairwise SNP-level replication using only genotype data from each cohort is shown in Figure 2**c**.

**Figure S18:**
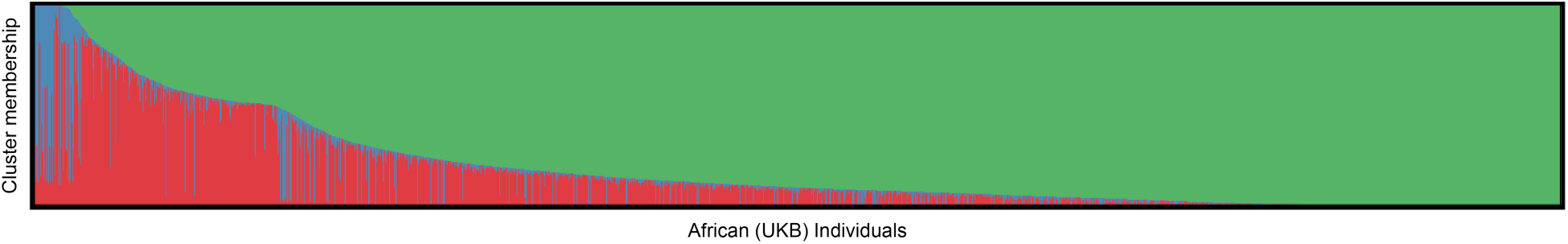
ADMIXTURE results from self-identified African individuals in the UK Biobank. We performed 10 runs of unsupervised ADMIXTURE ^91^ setting *K* = 3 on 4,967 self-identified African individuals in the UK Biobank. We included YRI and CEU individuals from the 1000 Genomes Project ^92^ to identify European and African ancestry components in the ADMIXTURE results. We then filtered out all individuals with less than 5% membership in the African component (identified as the component shown in green). The AIAN and European ancestry components are shown in blue and red, respectively. Our ADMIXTURE pipeline used the same protocol described in Bitarello and Mathieson ^84^. All scripts for ADMIXTURE runs as well as filtration steps are available on the GitHub page given in the Data Availability section.

**Figure S19:**
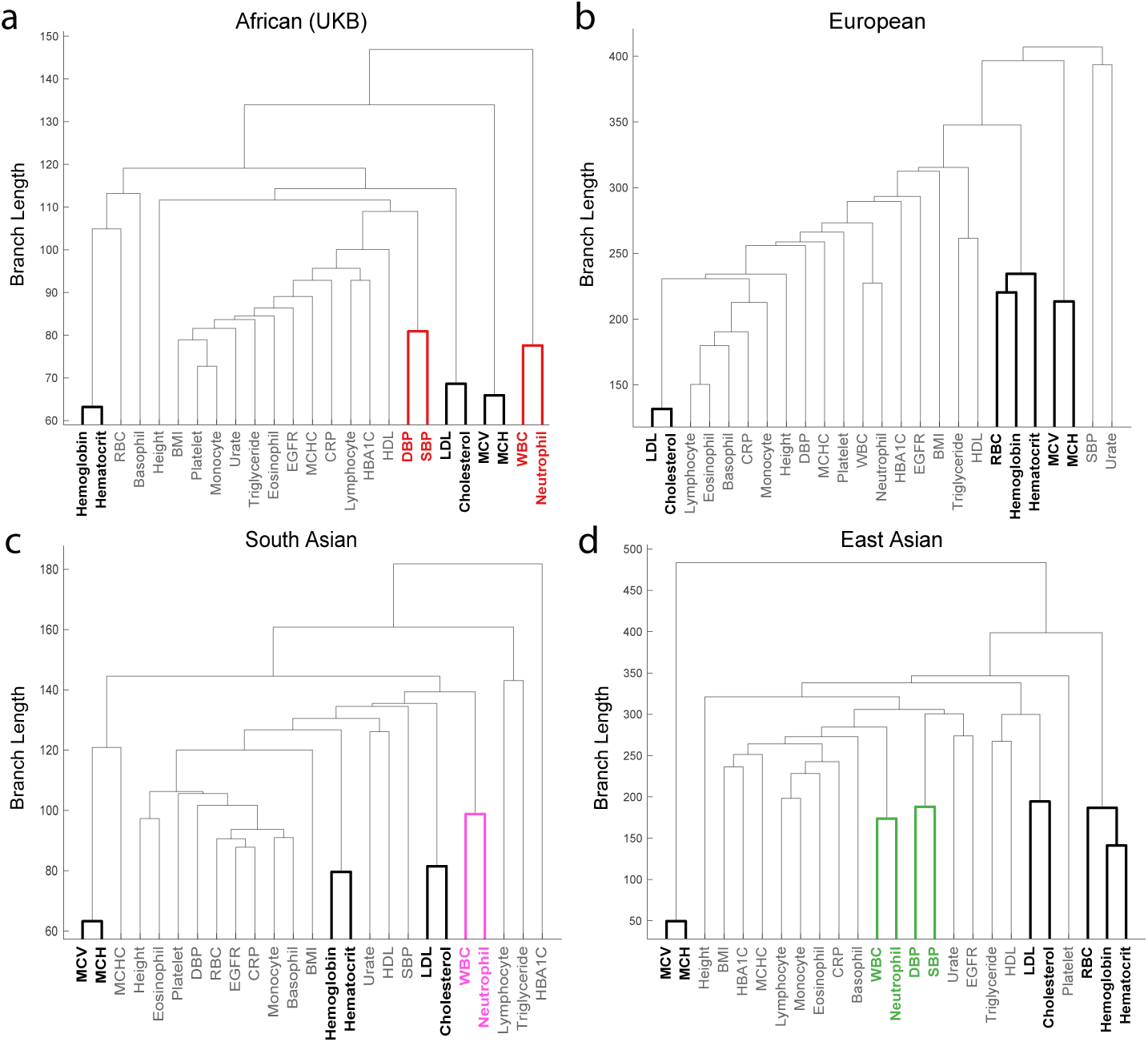
Multiple prioritized trait clusters with shared core genetic trait architecture replicate in the (a) African (UKB), (b) European, (c) South Asian, and (d) East Asian ancestry cohorts. The WINGS algorithm identified prioritized phenotype clusters in each of these ancestry cohorts, denoted in each dendrogram as clades with emboldened lines. Three clusters of phenotypes were found in all ancestries (shown and labeled in black), comprising: mean corpuscular volume (MCV) and mean corpuscular hemoglobin (MCH), hemoglobin and hematocrit, and the metabolic traits low-density lipoprotein (LDL) and cholesterol levels. In both the European and East Asian ancestry cohorts, red blood cell count (RBC) was also a member of the hemoglobin and hematocrit phenotype cluster. Two other phenotype clusters were identified in at least two ancestry cohorts. One of these clusters contains white blood cell count (WBC) and neutrophil count, and the other contains diastolic and systolic blood pressure (DBP and SBP). These two clusters are color-coded according to the ancestry cohorts in which they are prioritized. The WINGS algorithm was applied to traits from each ancestry cohort separately as described in the Supplemental Information.

## Supplemental Tables

**Table S1:** Ancestry cohorts analyzed in this study. In studies where GWA summary statistics were available to us, sample size and number of SNPs differ due to original study design. The specific sample size and number of SNPs for each trait in studies from Biobank Japan and PAGE are provided in Table S4-Table S9.

| Ancestry cohort label in this study | Study | Label from original study | Sample Size | Number of SNPs |
| --- | --- | --- | --- | --- |
| European | UK Biobank | Self-identified white british | 349,411 | 1,933,096 |
| East Asian | Biobank Japan | East Asian | 19,190 - 206,692 | 4,823,101 - 6,628,005 |
| Hispanic and Latin American | PAGE | Admixed Hispanic and Latin American | 15,522 - 21,955 | 8,576,622 - 8,822,607 |
| African-American (PAGE) | PAGE | African-American | 3,750 - 17,280 | 12,107,345 - 12,274,127 |
| South Asian | UK Biobank | Self-identified South Asian | 5,716 | 958,375 |
| African (UKB) | UK Biobank | African | 4,967 | 578,320 |
| Native Hawaiian | PAGE | Native Hawaiian | 1,777 - 3,938 | 6,656,996 - 6,966,169 |
| American Indian/Alaska Native | PAGE | AIAN | 574 - 645 | 3,970,247 - 8,504,923 |

**Table S2:**
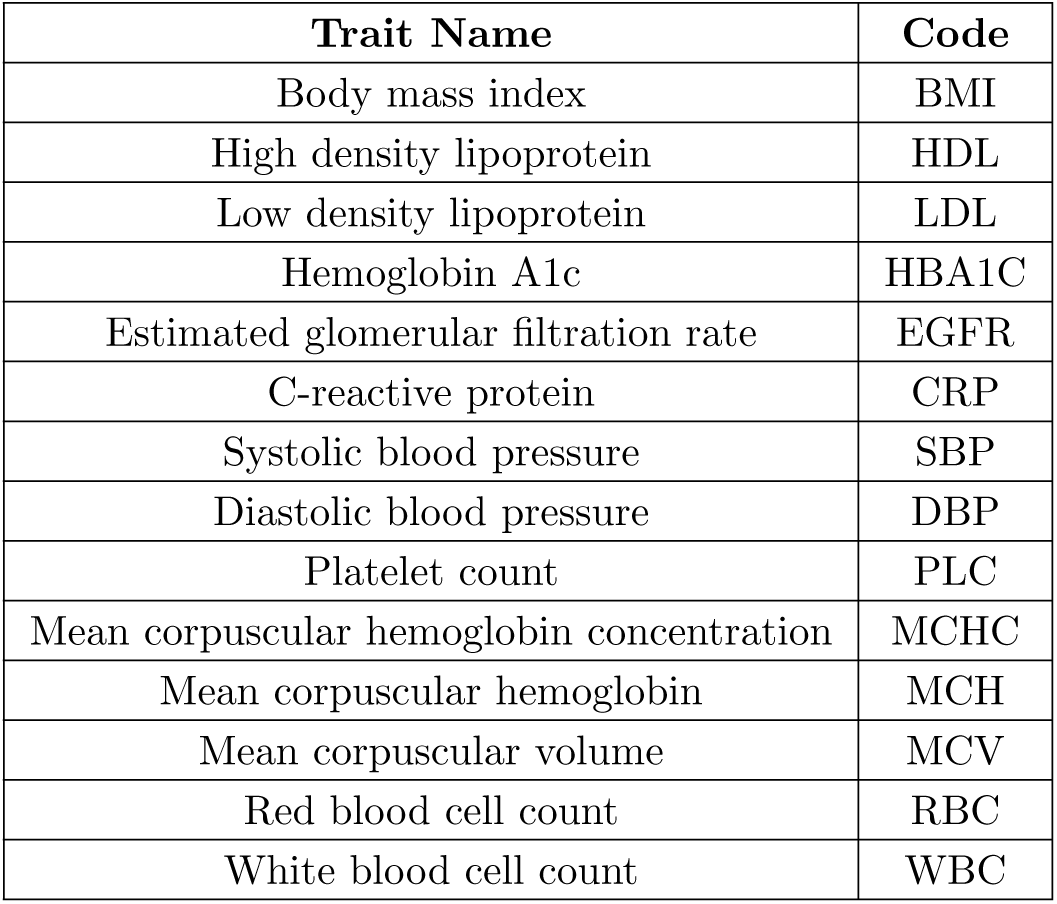
Abbreviations used throughout this study for 14 quantitative traits analyzed in this study. The remaining 11 traits analyzed were Basophil count, Cholesterol, Eosinophil count, Height, Hematocrit, Hemoglobin, Lymphocyte count, Monocyte count, Neutrophil count, and Triglyceride levels, respectively. These are not abbreviated in the main text.

| Trait Name | Code |
| --- | --- |
| Body mass index | BMI |
| High density lipoprotein | HDL |
| Low density lipoprotein | LDL |
| Hemoglobin A1c | HBA1C |
| Estimated glomerular filtration rate | EGFR |
| C-reactive protein | CRP |
| Systolic blood pressure | SBP |
| Diastolic blood pressure | DBP |
| Platelet count | PLC |
| Mean corpuscular hemoglobin concentration | MCHC |
| Mean corpuscular hemoglobin | MCH |
| Mean corpuscular volume | MCV |
| Red blood cell count | RBC |
| White blood cell count | WBC |

**Table S3:** Traits assayed in the PAGE study data by ancestry cohort. Data were available for each of the 25 listed traits in the UK Biobank European, South Asian, and African cohorts, as well as, the East Asian cohort from the Biobank Japan. Thus, each trait was analyzed in a minimum of four ancestries and a maximum of seven ancestries.

| **Trait Name or Code** | **AIAN** | **Native Hawaiian** | **Hispanic** |
| --- | --- | --- | --- |
| BMI | Yes | Yes | Yes |
| Basophil count | No | No | No |
| CRP | Yes | Yes | Yes |
| Cholesterol | Yes | Yes | Yes |
| DBP | Yes | No | Yes |
| EGFR | Yes | No | Yes |
| Eosinophil count | No | No | No |
| HBA1C | Yes | Yes | Yes |
| HDL | Yes | Yes | Yes |
| Height | Yes | Yes | Yes |
| Hematocrit | No | No | No |
| Hemoglobin | No | No | No |
| LDL | Yes | Yes | Yes |
| Lymphocyte count | No | No | No |
| MCH | Yes | No | Yes |
| MCHC | No | No | No |
| MCV | No | No | No |
| Monocyte count | No | No | No |
| Neutrophil count | No | No | No |
| PLC | Yes | No | Yes |
| RBC | No | No | No |
| SBP | Yes | No | Yes |
| Triglyceride | No | Yes | Yes |
| Urate | No | No | No |
| WBC | Yes | No | Yes |

**Table S4:** Number of individuals, total SNPs, pruned SNPs used for gene-*ε*, and genes and transcription factors (regions) included in the analysis for each trait in Biobank Japan data. Regions were defined using the hg19 list provided in Gusev et al. ^42^.

| Trait Name or Code | Sample Size | Total SNPs | Pruned SNPs | Regions | Citations |
| --- | --- | --- | --- | --- | --- |
| Basophil count | 62,076 | 5,653,566 | 410,465 | 23,106 | 93 |
| BMI | 158,284 | 5,653,566 | 410,465 | 23,085 | 94 |
| CRP | 75,391 | 5,608,701 | 408,166 | 23,108 | 93 |
| DBP | 136,615 | 5,653,566 | 410,465 | 23,085 | 93 |
| eGFR | 143,658 | 5,608,701 | 408,166 | 23,108 | 93 |
| Eosinophil count | 62,076 | 5,653,566 | 410,465 | 23,106 | 93 |
| HDL | 70,657 | 5,608,701 | 408,166 | 23,108 | 93 |
| Height | 159,095 | 6,296,332 | 354,647 | 23,679 | 95 |
| Hematocrit | 108,757 | 5,653,566 | 410,465 | 23,106 | 93 |
| Hemoglobin | 108,769 | 5,653,566 | 410,465 | 23,085 | 93 |
| HbA1c | 75,391 | 5,608,701 | 408,166 | 23,108 | 93 |
| LDL | 72,866 | 5,608,701 | 408,166 | 23,108 | 93 |
| Lymphocyte count | 62,076 | 5,653,566 | 410,465 | 23,106 | 93 |
| MCH | 108,054 | 5,653,566 | 410,465 | 23,106 | 93 |
| MCHC | 108,738 | 5,653,566 | 410,465 | 23,106 | 93 |
| MCV | 108,526 | 5,653,566 | 410,465 | 23,085 | 93 |
| Monocyte count | 62,076 | 5,653,566 | 410,465 | 23,106 | 93 |
| Neutrophil count | 62,076 | 5,653,566 | 410,465 | 23,106 | 93 |
| PLC | 108,208 | 5,653,566 | 410,465 | 23,085 | 93 |
| RBC | 108,794 | 5,653,566 | 410,465 | 23,085 | 93 |
| SBP | 136,597 | 5,653,566 | 410,465 | 23,085 | 93 |
| Cholesterol | 128,305 | 5,608,701 | 408,166 | 23,108 | 93 |
| Triglyceride | 105,597 | 5,608,701 | 410,465 | 23,108 | 93 |
| Urate | 109,029 | 5,608,701 | 408,166 | 23,108 | 93 |
| WBC | 107,694 | 5,653,566 | 408,166 | 23,085 | 93 |

**Table S5:** Number of individuals, total SNPs, pruned SNPs used for gene-*ε*, and genes and transcription factors (regions) included in the analysis for each trait in the African-American cohort of the PAGE study data.

| Trait Name or Code | Sample Size | Total SNPs | Pruned SNPs | Regions |
| --- | --- | --- | --- | --- |
| BMI | 17,127 | 12,139,115 | 404,401 | 24,216 |
| CRP | 8,349 | 12,274,126 | 404,572 | 24,206 |
| DBP | 11,380 | 12,148,801 | 405,188 | 24,218 |
| eGFR | 8,261 | 12,128,273 | 403,371 | 24,207 |
| Hemoglobin A1c | 17,127 | 12,139,115 | 404,401 | 24,215 |
| HDL | 10,085 | 12,114,827 | 404,089 | 24,201 |
| Height | 17,280 | 12,139,907 | 404,522 | 24,201 |
| LDL | 9,720 | 12,107,344 | 403,740 | 24,218 |
| MCHC | 3,750 | 12,132,232 | 405,558 | 24,217 |
| PLC | 8,850 | 12,131,935 | 404,497 | 24,193 |
| SBP | 11,380 | 12,148,801 | 405,188 | 24,218 |
| Cholesterol | 10,137 | 12,110,337 | 403,674 | 24,222 |
| Triglyceride | 9,980 | 12,110,879 | 403,455 | 24,206 |
| WBC | 8,802 | 12,126,732 | 404,579 | 24,219 |

**Table S6:** Number of individuals, total SNPs, pruned SNPs used for gene-*ε*, and genes and transcription factors (regions) included in the analysis for each trait in the Hispanic and Latin American cohort of the PAGE study data.

| Trait Name or Code | Sample Size | Total SNPs | Pruned SNPs | Region Count |
| --- | --- | --- | --- | --- |
| BMI | 21,955 | 8,812,436 | 432,762 | 24,138 |
| CRP | 15,912 | 8,576,621 | 397,941 | 24,118 |
| DBP | 21,549 | 8,784,112 | 430,360 | 24,126 |
| Estimated glomerular filtration rate | 18,548 | 8,702,426 | 422,598 | 24,123 |
| HbA1c | 21,955 | 8,812,436 | 432,762 | 24,138 |
| HDL | 17,751 | 8,583,603 | 412,771 | 24,122 |
| Height | 22,187 | 8,822,606 | 433,604 | 24,132 |
| LDL | 17,373 | 8,588,800 | 413,074 | 24,116 |
| MCHC | 15,522 | 8,763,739 | 427,208 | 24,132 |
| PLC | 18,949 | 8,612,804 | 415,201 | 24,115 |
| SBP | 21,549 | 8,784,112 | 430,360 | 24,126 |
| Cholesterol | 17,802 | 8,586,887 | 412,830 | 24,115 |
| Triglyceride | 17,856 | 8,594,121 | 413,546 | 24,104 |
| WBC | 18,206 | 8,603,503 | 414,462 | 24,123 |

**Table S7:** Number of individuals, total SNPs, pruned SNPs used for gene-ε, and genes and transcription factors (regions) included in the analysis for each trait in the AIAN cohort of the PAGE study data.

| Trait Name or Code | Sample Size | Total SNPs | Pruned SNPs | Regions |
| --- | --- | --- | --- | --- |
| BMI | 645 | 8,374,976 | 421,826 | 24,124 |
| CRP | 574 | 8,504,922 | 417,287 | 24,136 |
| DBP | 636 | 8,376,521 | 421,528 | 24,136 |
| eGFR | 602 | 8,336,044 | 417,540 | 24,132 |
| Hemoglobin A1c | 645 | 8,374,976 | 421,826 | 24,124 |
| HDL | 604 | 8,315,912 | 415,939 | 24,121 |
| Height | 645 | 8,375,624 | 421,750 | 24,117 |
| LDL | 591 | 8,360,719 | 419,544 | 24,123 |
| MCHC | 20 | 3,970,246 | 62,339 | 17,381 |
| PLC | 603 | 8,294,302 | 414,530 | 24,133 |
| Systolic blood pressure | 636 | 8,376,521 | 421,528 | 24,136 |
| Cholesterol | 604 | 8,586,887 | 415,939 | 24,121 |
| WBC | 602 | 8,289,567 | 414,462 | 24,133 |

**Table S8:** Number of individuals, total SNPs, pruned SNPs used for gene-*ε*, and genes and transcription factors (regions) included in the analysis for each trait in the Native Hawaiian (Native Hawaiian) cohort of the PAGE study data.

| Trait Name or Code | Sample Size | Total SNPs | Pruned SNPs | Regions |
| --- | --- | --- | --- | --- |
| BMI | 3,936 | 6,664,738 | 415,221 | 23,885 |
| CRP | 1,777 | 6,966,169 | 428,517 | 23,834 |
| Hemoglobin A1c | 3,936 | 6,664,738 | 415,221 | 23,885 |
| HDL | 1,912 | 6,656,996 | 416,255 | 23,894 |
| Height | 3,938 | 6,660,920 | 415,172 | 23,878 |
| LDL | 1,900 | 6,662,802 | 416,810 | 23,895 |
| Cholesterol | 1,915 | 6,660,807 | 416,425 | 23,899 |
| Triglycerides | 1,915 | 6,660,807 | 416,425 | 23,899 |

**Table S9:** Number of individuals, total SNPs, pruned SNPs used for gene-ε, and genes and transcription factors (regions) included in the analysis for each trait in the Asian cohort of the PAGE study data.

| Trait Name or Code | Sample Size | Total SNPs | Pruned SNPs | Regions |
| --- | --- | --- | --- | --- |
| BMI | 4,647 | 15,362,633 | 433,356 | 24,085 |
| CRP | 1,811 | 14,374,461 | 428,656 | 24,116 |
| DBP | 1,086 | 12,470,507 | 416,273 | 24,112 |
| eGFR | 150 | 8,314,417 | 337,167 | 24,017 |
| HbA1c | 4,647 | 15,362,633 | 433,356 | 24,085 |
| HDL | 2,378 | 13,413,244 | 428,598 | 24,072 |
| Height | 4,679 | 15,366,710 | 433,005 | 24,103 |
| LDL | 2,316 | 13,327,313 | 428,741 | 24,075 |
| MCHC | 128 | 8,089,136 | 315,583 | 23,946 |
| PLC | 541 | 10,528,072 | 421,929 | 24,098 |
| SBP | 1,086 | 12,470,507 | 416,273 | 24,112 |
| Cholesterol | 2,387 | 13,436,190 | 428,656 | 24,078 |
| Triglyceride | 2,381 | 13,423,953 | 429,246 | 24,073 |
| WBC | 543 | 10,570,051 | 421,776 | 24,095 |

**Table S10:**
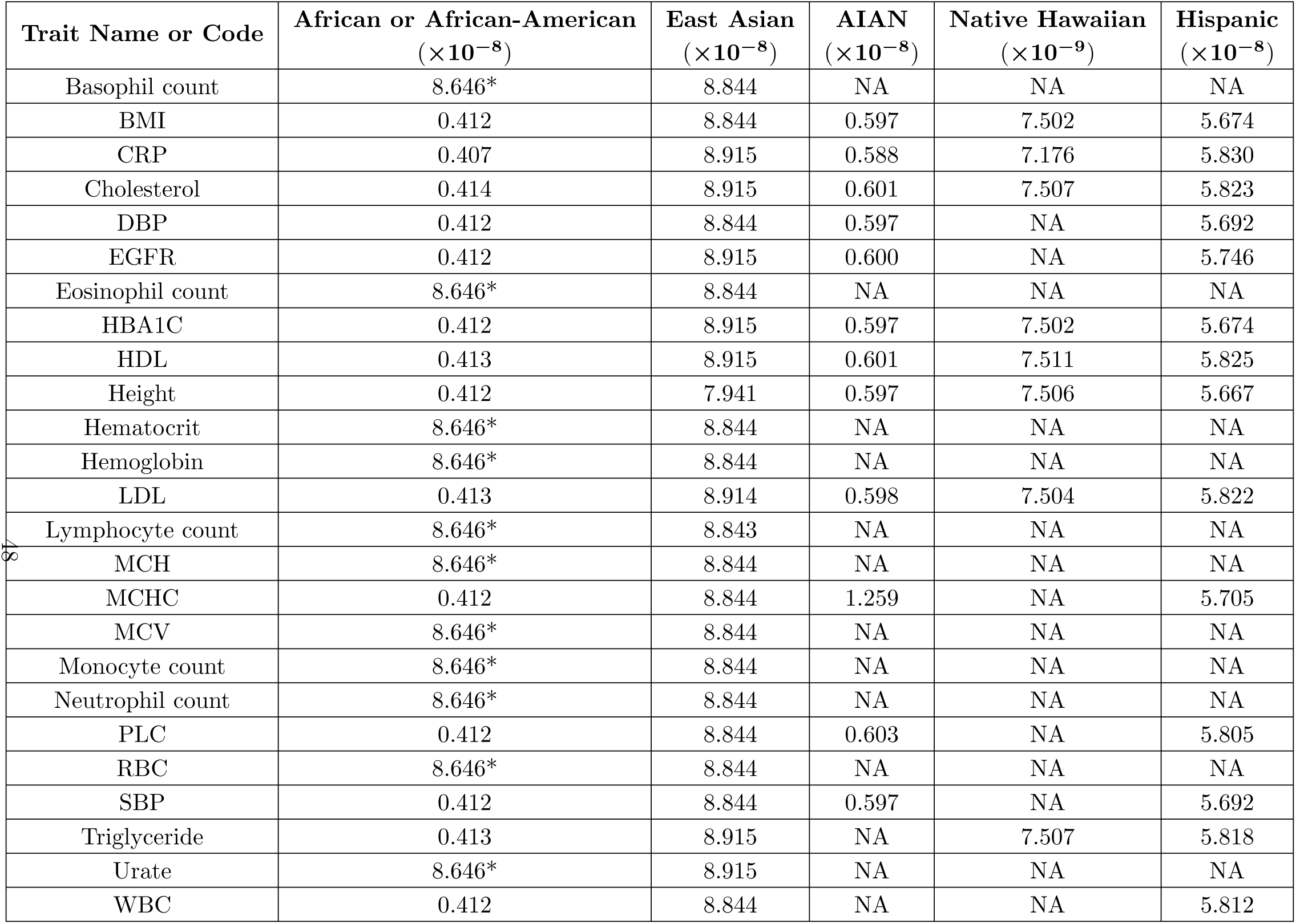
Bonferonni *p*-value threshold corrected for number of SNP-level association tests performed for each ancestry-trait pair. Thresholds are calculated as 0.05 divided by the number of SNPs tested in each ancestry-trait pair. The term“NA” indicates that there was no data for that ancestry-trait pair. The threshold for Bonferroni-corrected significance was the same for every trait in the European (*p*-value *<* 2.587 10*^−^*^8^) and South Asian (*p*-value *<* 5.217 10*^−^*^8^) cohorts from the UK Biobank. Traits for which the UK Biobank African cohort was used are denoted with a *; otherwise, the African-American cohort from the PAGE study data was used. See Table S2 for expansion of trait codes.

| Trait Name or Code | African or African-American<br>($\times 10^{-8}$) | East Asian<br>($\times 10^{-8}$) | AIAN<br>($\times 10^{-8}$) | Native Hawaiian<br>($\times 10^{-9}$) | Hispanic<br>($\times 10^{-8}$) |
| --- | --- | --- | --- | --- | --- |
| Basophil count | 8.646* | 8.844 | NA | NA | NA |
| BMI | 0.412 | 8.844 | 0.597 | 7.502 | 5.674 |
| CRP | 0.407 | 8.915 | 0.588 | 7.176 | 5.830 |
| Cholesterol | 0.414 | 8.915 | 0.601 | 7.507 | 5.823 |
| DBP | 0.412 | 8.844 | 0.597 | NA | 5.692 |
| EGFR | 0.412 | 8.915 | 0.600 | NA | 5.746 |
| Eosinophil count | 8.646* | 8.844 | NA | NA | NA |
| HBA1C | 0.412 | 8.915 | 0.597 | 7.502 | 5.674 |
| HDL | 0.413 | 8.915 | 0.601 | 7.511 | 5.825 |
| Height | 0.412 | 7.941 | 0.597 | 7.506 | 5.667 |
| Hematocrit | 8.646* | 8.844 | NA | NA | NA |
| Hemoglobin | 8.646* | 8.844 | NA | NA | NA |
| LDL | 0.413 | 8.914 | 0.598 | 7.504 | 5.822 |
| Lymphocyte count | 8.646* | 8.843 | NA | NA | NA |
| MCH | 8.646* | 8.844 | NA | NA | NA |
| MCHC | 0.412 | 8.844 | 1.259 | NA | 5.705 |
| MCV | 8.646* | 8.844 | NA | NA | NA |
| Monocyte count | 8.646* | 8.844 | NA | NA | NA |
| Neutrophil count | 8.646* | 8.844 | NA | NA | NA |
| PLC | 0.412 | 8.844 | 0.603 | NA | 5.805 |
| RBC | 8.646* | 8.844 | NA | NA | NA |
| SBP | 0.412 | 8.844 | 0.597 | NA | 5.692 |
| Triglyceride | 0.413 | 8.915 | NA | 7.507 | 5.818 |
| Urate | 8.646* | 8.915 | NA | NA | NA |
| WBC | 0.412 | 8.844 | NA | NA | 5.812 |

**Table S11:** Replication of effect sizes and posterior inclusion probabilities (PIPs) among ten independent subsamples of the UK Biobank European ancestry cohort using SuSiE ^34^ for fine-mapping. The sample size of the ten independent, non-overlapping subsamples of the UK Biobank European ancestry cohort was set to 10,000. For the 1,895,051 SNPs that were analyzed in every European ancestry cohort subsample (Table S1) and the effect sizes and PIPs (columns 2 and 4, respectively) generated using the SuSiE method ^34^, we calculated the median correlation coefficient between all possible pairwise comparisons (10 choose 2) of the European ancestry cohort subsamples. Column 3 reports the mean number of SNPs with effect sizes greater than 0.1 across all ten European ancestry cohort subsamples for each trait. Column 5 reports the mean number of SNPs with a posterior inclusion probability greater than 0.01 across the ten European ancestry cohort subsamples for each trait.

| Trait Name or Code | Median Effect Size Correlation | Effect Sizes > 0.1 (Mean) | Median PIP Correlation | PIPs > 0.01 (Mean) |
| --- | --- | --- | --- | --- |
| BMI | 0.0143 | 0 | 0.073 | 67.3 |
| Basophil | $3.69 \times 10^{-6}$ | 0 | 0.002 | 211.8 |
| CRP | 0.096 | 0.7 | 0.159 | 142.6 |
| Cholesterol | 0.966 | 1 | 0.232 | 53.1 |
| DBP | 0.001 | 0.6 | 0.012 | 173.1 |
| EGFR | $4.71 \times 10^{-6}$ | 0 | 0.005 | 149 |
| Eosinophil | 0.0179 | 0.2 | 0.042 | 325.8 |
| HBA1C | 0.012 | 0.2 | 0.025 | 26.8 |
| HDL | 0.325 | 0.1 | 0.219 | 57.6 |
| Height | $9.20 \times 10^{-5}$ | 1.6 | 0.010 | 187.7 |
| Hematocrit | $-4.16 \times 10^{-6}$ | 0.5 | 0.015 | 40.3 |
| Hemoglobin | $-1.13 \times 10^{-5}$ | 0.6 | 0.014 | 45.8 |
| LDL | 0.949 | 1 | 0.331 | 60.9 |
| Lymphocyte | $-1.01 \times 10^{-6}$ | 1.5 | 0.001 | 1368.5 |
| MCH | $7.098 \times 10^{-6}$ | 0.6 | 0.007 | 60.9 |
| MCHC | $-2.00 \times 10^{-6}$ | 0.9 | 0.01 | 58.7 |
| MCV | $1.71 \times 10^{-5}$ | 0.6 | 0.01 | 70 |
| Monocyte | 0.002 | 0.1 | 0.017 | 349.4 |
| Neutrophil | 0.038 | 0.1 | 0.071 | 66.5 |
| PLC | 0.609 | 0.9 | 0.154 | 112.5 |
| RBC | $9.650 \times 10^{-5}$ | 0.9 | 0.028 | 49.7 |
| SBP | 0.001 | 0.6 | 0.019 | 166.8 |
| Triglyceride | 0.348 | 0.0 | 0.247 | 164.7 |
| Urate | 0.252 | 0.5 | 0.24 | 39.9 |
| WBC | 0.0002 | 0.1 | 0.01 | 347.8 |

**Table S12:** Replication of effect sizes and posterior inclusion probabilities (PIPs) between the UK Biobank African ancestry cohort and ten independent subsamples of the UK Biobank European ancestry cohort using SuSiE ^34^ for fine-mapping. Each of the ten independent, non-overlapping subsamples of the UK Biobank European ancestry cohort was set to be equal in size to the sample size of the African ancestry cohort (*N* = 4,967), Table S1. Column headers containing ”(mean)” indicate a mean is generated averaging over the ten independent European ancestry cohort subsamples. For the 496,997 SNPs that were analyzed in both the African ancestry cohort and every European ancestry cohort subsample, we compared the SuSiE ^34^ effect size estimates and PIPs. For both effect sizes and PIPs, the median correlation coefficient between the African ancestry cohort and the pairwise comparison to each European ancestry cohort subsample is reported in the third and seventh columns, respectively. Column 3 reports the total number of SNPs with effect sizes greater than 0.1 in the African cohort. Column four gives the mean number of effect sizes greater than 0.1 in the European ancestry cohort subsamples for each trait. We performed the same comparison for the PIPs using a threshold of 0.01. Column 2 reports the number of SNPs that surpassed an effect size of 0.1 in both the African ancestry cohort and at least one of the European ancestry cohort subsamples. Column 6 reports the number of SNPs that surpasses a PIP of 0.01 in the African ancestry cohort and at least one European ancestry cohort subsample.

| Trait Name<br>or Code | Number of<br>Shared<br>Effect Sizes<br>> 0.1 | Median<br>Effect Size<br>Correlation | African Effect<br>Sizes > 0.1 | European<br>Effect Sizes<br>> 0.1<br>(Mean) | Number of<br>Shared PIPs<br>> 0.01 | Median PIP<br>Correlation | African<br>PIPs > 0.01 | European PIPs<br>> 0.01 (Mean) |
| --- | --- | --- | --- | --- | --- | --- | --- | --- |
| BMI | 0 | $5.12 \times 10^{-5}$ | 2 | 0 | 1 | 0.027 | 1206 | 8.9 |
| Basophil | 0 | $-3.01 \times 10^{-5}$ | 0 | 0 | 0 | 0.001 | 403 | 40.3 |
| CRP | 0 | $-3.68 \times 10^{-6}$ | 0 | 0.4 | 0 | $5.88 \times 10^{-5}$ | 95 | 33.6 |
| Cholesterol | 1 | 0.5 | 1 | 0.9 | 5 | 0.467 | 1144 | 8.4 |
| DBP | 0 | $1.58 \times 10^{-5}$ | 10 | 0.3 | 0 | 0.008 | 1249 | 20.3 |
| EGFR | 0 | $2.41 \times 10^{-6}$ | 1 | 0 | 0 | 0.001 | 984 | 24.7 |
| Eosinophil | 0 | $-1.97 \times 10^{-5}$ | 0 | 0 | 0 | 0.003 | 979 | 29.3 |
| HBA1C | 0 | $-1.15 \times 10^{-6}$ | 5 | 0.2 | 0 | 0.004 | 959 | 6.5 |
| HDL | 0 | 0.014 | 0 | 0.1 | 2 | 0.014 | 31 | 11.6 |
| Height | 0 | $-1.22 \times 10^{-5}$ | 19 | 0.7 | 0 | 0.003 | 208 | 33.7 |
| Hematocrit | 0 | $-1.14 \times 10^{-5}$ | 1 | 0 | 0 | 0.016 | 199 | 12.2 |
| Hemoglobin | 0 | $-6.31 \times 10^{-6}$ | 1 | 0 | 0 | 0.022 | 568 | 12.5 |
| LDL | 1 | 0.674 | 1 | 1 | 8 | 0.497 | 995 | 13.5 |
| Lymphocyte | 0 | $2.71 \times 10^{-5}$ | 0 | 0 | 1 | 0.001 | 60 | 115.5 |
| MCH | 0 | $1.14 \times 10^{-5}$ | 1 | 0.2 | 0 | 0.005 | 1332 | 11.8 |
| MCHC | 0 | $-1.56 \times 10^{-6}$ | 1 | 0.2 | 0 | 0.008 | 646 | 13.6 |
| MCV | 0 | $2.58 \times 10^{-6}$ | 1 | 0.2 | 0 | 0.005 | 917 | 11.8 |
| Monocyte | 0 | $-2.44 \times 10^{-5}$ | 0 | 0 | 2 | 0.002 | 2215 | 27.7 |
| Neutrophil | 0 | -0.0001 | 1 | 0 | 0 | 0.001 | 2386 | 10.2 |
| PLC | 0 | -0.017 | 1 | 0.6 | 3 | 0.035 | 2088 | 13.7 |
| RBC | 0 | $-6.35 \times 10^{-6}$ | 1 | 0.1 | 0 | -0.002 | 3 | 12.7 |
| SBP | 0 | $4.31 \times 10^{-5}$ | 13 | 0.1 | 0 | 0.008 | 1133 | 20 |
| Triglyceride | 0 | 0.004 | 0 | 0.1 | 8 | 0.009 | 2231 | 25.6 |
| Urate | 1 | 0.01 | 2 | 0.3 | 3 | 0.051 | 334 | 9.8 |
| WBC | 0 | -0.0001 | 1 | 0 | 5 | 0.0006 | 2862 | 48.3 |

**Table S13:** Replication of effect sizes and posterior inclusion probabilities (PIPs) between the UK Biobank South Asian ancestry cohort and ten independent subsamples of the UK Biobank European ancestry cohort using SuSiE ^34^ for fine-mapping. Each of the ten independent, non-overlapping subsamples of the UK Biobank European ancestry cohort was set to be equal in size to the sample size of the South Asian ancestry cohort (*N* = 5,660), Table S1. Column headers containing ”(mean)” indicate a mean is generated averaging over the ten independent European ancestry cohort subsamples. For the 863,569 SNPs that were analyzed in both the South Asian ancestry cohort and every European ancestry cohort subsample, we compared the SuSiE ^34^ effect size estimates and PIPs. For both effect sizes and PIPs, the median correlation coefficient between the South Asian ancestry cohort and the pairwise comparison to each European ancestry cohort subsample is reported in the third and seventh columns, respectively. Column 3 reports the total number of SNPs with effect sizes greater than 0.1 in the South Asian cohort. Column four gives the mean number of effect sizes greater than 0.1 in the European ancestry cohort subsamples for each trait. We performed the same comparison for the PIPs using a threshold of 0.01. Column 2 reports the number of SNPs that surpassed an effect size of 0.1 in both the South Asian ancestry cohort and at least one of the European ancestry cohort subsamples. Column 6 reports the number of SNPs that surpasses a PIP of 0.01 in the South Asian ancestry cohort and at least one European ancestry cohort subsample.

| Trait Name or Code | Number of Shared Effect Sizes $> 0.1$ | Median Effect Size Correlation | South Asian Effect Sizes $> 0.1$ | European Effect Sizes $> 0.1$ (Mean) | Number of Shared PIPs $> 0.01$ | Median PIP Correlation | South Asian PIPs $> 0.01$ | European PIPs $> 0.01$ (Mean) |
| --- | --- | --- | --- | --- | --- | --- | --- | --- |
| BMI | 0 | $7.07 \times 10^{-5}$ | 1 | 0 | 0 | 0.0173 | 180 | 18 |
| Basophil | 0 | $-1.10 \times 10^{-5}$ | 0 | 0 | 0 | 0.0008 | 210 | 78.4 |
| CRP | 0 | 0.0309 | 0 | 0.4 | 17 | 0.121 | 181 | 64.7 |
| Cholesterol | 0 | 0.004 | 0 | 0.9 | 2 | 0.0127 | 20 | 9 |
| DBP | 0 | $-2.12 \times 10^{-5}$ | 0 | 0.6 | 0 | 0.013 | 115 | 40 |
| EGFR | 0 | $-1.26 \times 10^{-5}$ | 0 | 0 | 0 | $5.12 \times 10^{-5}$ | 452 | 44.2 |
| Eosinophil | 0 | $2.16 \times 10^{-5}$ | 0 | 0 | 0 | 0.006 | 63 | 69.1 |
| HBA1C | 0 | $-3.33 \times 10^{-5}$ | 2 | 0.3 | 0 | 0.011 | 133 | 13.3 |
| HDL | 0 | 0.525 | 0 | 0.1 | 3 | 0.505 | 78 | 12.2 |
| Height | 0 | $-2.13 \times 10^{-5}$ | 12 | 1.1 | 1 | 0.001 | 754 | 66.8 |
| Hematocrit | 0 | $-4.78 \times 10^{-6}$ | 1 | 0.3 | 0 | $6.24 \times 10^{-5}$ | 24 | 17.4 |
| Hemoglobin | 0 | $-4.19 \times 10^{-6}$ | 1 | 0.3 | 0 | 0.011 | 19 | 16.1 |
| LDL | 0 | 0.491 | 1 | 1 | 14 | 0.386 | 44 | 15.6 |
| Lymphocyte | 0 | $-4.40 \times 10^{-6}$ | 0 | 1.1 | 0 | 0.0001 | 113 | 343.1 |
| MCH | 0 | $5.29 \times 10^{-6}$ | 2 | 0.5 | 0 | 0.008 | 78 | 22.3 |
| MCHC | 0 | $-5.77 \times 10^{-6}$ | 1 | 0.9 | 0 | 0.007 | 45 | 20.3 |
| MCV | 0 | $1.74 \times 10^{-5}$ | 3 | 0.5 | 0 | 0.005 | 81 | 26.3 |
| Monocyte | 0 | $-4.05 \times 10^{-6}$ | 0 | 0.1 | 0 | 0.001 | 156 | 76.8 |
| Neutrophil | 0 | 0.221 | 0 | 0 | 16 | 0.172 | 70 | 25.7 |
| PLC | 0 | 0.577 | 0 | 0.8 | 1 | 0.679 | 126 | 22.7 |
| RBC | 0 | $2.79 \times 10^{-6}$ | 0 | 0.3 | 0 | -0.011 | 41 | 20.9 |
| SBP | 0 | -0.0004 | 0 | 0.3 | 0 | 0.037 | 116 | 43.2 |
| Triglyceride | 0 | 0.158 | 0 | 0 | 53 | 0.173 | 189 | 46.4 |
| Urate | 0 | 0.137 | 1 | 0.4 | 6 | 0.150 | 52 | 15.8 |
| WBC | 0 | 0.001 | 0 | 0 | 18 | 0.008 | 52 | 93 |

**Table S14:** Overlap between SNPs identified by PESCA and GWA analyses in the European ancestry cohort from the UK Biobank and the East Asian ancestry cohort from the Biobank Japan. For seven continuous traits, we compared SNP-level association *p*-values from our analysis to the posterior probabilities calculated in Shi et al. ^51^ using the PESCA framework. For each of the seven traits, there were SNPs that had a posterior probability *>* 0.8 of being causal in both ancestries and were also nominally significant (*p*-value 5 *<* 10*^−^*^6^) using the standard GWA SNP-level framework.

| Trait | PESCA EUR<br>only and EUR<br>GWA nominal<br>significance | PESCA EAS<br>only and EAS<br>GWA nominal<br>significance | PESCA both<br>and EUR GWA<br>nominal<br>significance | PESCA both<br>and EAS GWA<br>nominal<br>significance | PESCA both<br>and GWA<br>nominal<br>significance in<br>both |
| --- | --- | --- | --- | --- | --- |
| BMI | 0 | 0 | 97 | 539 | 94 |
| Cholesterol | 2 | 13 | 14 | 54 | 11 |
| HDL | 5 | 8 | 29 | 74 | 27 |
| LDL | 0 | 6 | 16 | 43 | 13 |
| MCH | 3 | 14 | 39 | 180 | 39 |
| MCV | 1 | 11 | 44 | 215 | 44 |
| Triglyceride | 4 | 8 | 15 | 49 | 15 |

**Table S15:** Bonferonni p-value threshold corrected for number of gene-level association tests performed for each ancestry-trait pair. Thresholds are calculated as 0.05 divided by the number of SNPs tested in each ancestry-trait pair. The term“NA” indicates that there was no data for that ancestry-trait pair. The threshold for Bonferroni-corrected significance was the same for every trait in the European (*p*-value *<* 2.092 10*^−^*^6^) and South Asian (*p*-value *<* 1.085 10*^−^*^6^) cohorts from the UK Biobank. Traits for which the UK Biobank African cohort was used are denoted with a *; otherwise, the African-American cohort from the PAGE study data was used. See Table S2 for expansion of trait codes.

| Trait | African or African-American<br>($\times 10^{-6}$) | East Asian<br>($\times 10^{-6}$) | AIAN<br>($\times 10^{-6}$) | Native Hawaiian<br>($\times 10^{-6}$) | Hispanic and Latin American<br>($\times 10^{-6}$) |
| --- | --- | --- | --- | --- | --- |
| Basophil count | 1.121 | 2.197 | NA | NA | NA |
| BMI | 2.096 | 2.197 | 2.073 | 2.093 | 2.072 |
| CRP | 2.097 | 2.085 | 2.072 | 2.100 | 2.073 |
| Cholesterol | 2.096 | 2.198 | 2.073 | 2.092 | 2.073 |
| DBP | 2.096 | 2.197 | 2.072 | NA | 2.073 |
| EGFR | 2.097 | 2.198 | 2.073 | NA | 2.073 |
| Eosinophil count | 2.121* | 2.197 | NA | NA | NA |
| HBA1C | 2.096 | 2.198 | 2.073 | 2.093 | 2.072 |
| HDL | 2.097 | 2.198 | 2.073 | 2.093 | 2.073 |
| Height | 2.097 | 2.131 | 2.073 | 2.094 | 2.072 |
| Hematocrit | 2.121* | 2.197 | NA | NA | NA |
| Hemoglobin | 2.121* | 2.197 | NA | NA | NA |
| LDL | 2.096 | 2.198 | 2.073 | 2.093 | 2.073 |
| Lymphocyte count | 2.121* | 2.197 | NA | NA | NA |
| MCH | 2.121* | 2.197 | NA | NA | NA |
| MCHC | 2.096 | 2.197 | 2.877 | NA | 2.072 |
| MCV | 2.121* | 2.197 | NA | NA | NA |
| Monocyte count | 2.121* | 2.197 | NA | NA | NA |
| Neutrophil count | 2.121* | 2.197 | NA | NA | NA |
| PLC | 2.097 | 2.197 | 2.072 | NA | 2.073 |
| RBC | 2.121* | 2.197 | NA | NA | NA |
| SBP | 0.412 | 8.844 | 0.597 | NA | 5.692 |
| Triglyceride | 2.096 | 2.197 | NA | 2.072 | 2.073 |
| Urate | 2.121* | 2.197 | NA | NA | NA |
| WBC | 2.092 | 2.197 | 2.072 | NA | 2.073 |

**Table S16:** Genes shown in Figure 3 as associated with triglyceride levels are supported by publications in the GWAS Catalog. Each of the genes listed is present in the significantly mutated subnetworks identified using Hierarchical HotNet ^31^ as enriched for associations with triglyceride levels in the European, East Asian, or Native Hawaiian ancestry cohorts. We mapped SNP-level associations from the GWAS Catalog to the 29 genes present in the significantly mutated subnetworks shown in Figure 3 (using the gene list provided by Gusev et al. ^42^) to generate the results for the 16 genes shown here.

| Gene | African or African-American | European | East Asian | Hispanic |
| --- | --- | --- | --- | --- |
| *ANGPTL4* | 96 | 97 | NA | 97 |
| *APOA1* | 98 | 98 | NA | 98 |
| *APOA4* | NA | 99 | NA | NA |
| *APOA5* | 100 | 100–103 | 104 | 100 |
| *APOB* | 105 | 105 | NA | 105 |
| *APOC1* | 98 | 99 | NA | 98 |
| *APOC2* | 100 | 100 | NA | 100 |
| *APOC3* | 98 | 98,99 | NA | 98 |
| *APOC4* | 99 | 99 | NA | 99 |
| *APOE* | 99 | 99,100 | NA | 99 |
| *CETP* | 98 | 99 | NA | 98 |
| *LMF1* | NA | 99 | NA | NA |
| *LPL* | 98 | 99 | 104 | 98 |
| *PCSK6* | 96 | 99 | NA | 96 |
| *PCSK7* | 100 | 100 | 104 | 100 |
| *PLTP* | 97,98,100,105 | 97–99,105–107 | NA | 97,98,100,105 |

**Table S17:** Gene-level association p-values for seven genes associated with platelet count in at last two ancestry cohorts. Of the 65 genes that were associated with platelet count in at least two ancestry cohorts, these seven contained previously identified SNP-level associations in studies submitted to the GWAS Catalog. Previous associations in the GWAS Catalog are discussed in the Supplemental Information. Ancestry-specific Bonferroni corrected significance thresholds for gene-level association analysis of platelet count are shown in Table S15.

| Gene | African-American | European | South Asian | East Asian | AIAN | Hispanic and Latin American |
| --- | --- | --- | --- | --- | --- | --- |
| *RDH13* | $4.14 \times 10^{-10}$ | $9.95 \times 10^{-1}$ | $8.80 \times 10^{-2}$ | $1.76 \times 10^{-6}$ | 1 | $7.88 \times 10^{-1}$ |
| *AGPAT5* | 1 | $1.30 \times 10^{-6}$ | $7.83 \times 10^{-1}$ | $5.00 \times 10^{-1}$ | $7.33 \times 10^{-8}$ | 1 |
| *GP6* | $7.20 \times 10^{-10}$ | $9.93 \times 10^{-1}$ | $1.47 \times 10^{-1}$ | $9.07 \times 10^{-7}$ | 1 | $6.92 \times 10^{-1}$ |
| *ALDH2* | 1 | $1.00 \times 10^{-20}$ | $1.13 \times 10^{-2}$ | $1.00 \times 10^{-20}$ | 1 | 1 |
| *RAB8A* | $9.57 \times 10^{-1}$ | $1.00 \times 10^{-20}$ | 1 | $5.76 \times 10^{-6}$ | 1 | $9.97 \times 10^{-1}$ |
| *CUX2* | 1 | $5.13 \times 10^{-7}$ | $1.16 \times 10^{-1}$ | $3.44 \times 10^{-11}$ | 1 | 1 |
| *ACAD10* | 1 | $1.47 \times 10^{-10}$ | $1.10 \times 10^{-2}$ | $2.00 \times 10^{-10}$ | 1 | 1 |

**Table S18:** Gene-*ε p*-values for the 28 genes present in the significantly mutated sunnetworks associated with triglyceride level in the European, East Asian, and Native Hawaiian cohorts. Each of these genes is present in Figure 3 which depicts the overlapping significantly mutated subnetworks identified using Hierarchical HoNet ^31^ identified in an analysis of triglyceride levels in the European, East Asian, and Native Hawaiian cohorts. Known SNP-level associations identified within the bounds of these genes in previous studies submitted to the GWAS Catalog are discussed in the Supplemental Information. Ancestry-specific Bonferroni corrected significance thresholds for gene-level association analysis of triglyceride levels are shown in Table S15.

| Gene | African-American<br>(PAGE) | European | South Asian | East Asian | Native<br>Hawaiian | Hispanic and<br>Latin American |
| --- | --- | --- | --- | --- | --- | --- |
| *APOA1* | $4.99 \times 10^{-1}$ | $1.00 \times 10^{-20}$ | $9.91 \times 10^{-1}$ | $1.00 \times 10^{-20}$ | $7.52 \times 10^{-1}$ | $4.99 \times 10^{-1}$ |
| *APOA4* | $4.99 \times 10^{-1}$ | $1.00 \times 10^{-20}$ | $2.51 \times 10^{-5}$ | $1.00 \times 10^{-20}$ | $9.15 \times 10^{-1}$ | $4.99 \times 10^{-1}$ |
| *APOA5* | $4.99 \times 10^{-1}$ | $1.42 \times 10^{-11}$ | $1.60 \times 10^{-6}$ | $9.95 \times 10^{-1}$ | $3.67 \times 10^{-12}$ | $4.99 \times 10^{-1}$ |
| *APOC3* | $4.99 \times 10^{-1}$ | $1.00 \times 10^{-20}$ | $9.82 \times 10^{-1}$ | $9.83 \times 10^{-1}$ | $3.05 \times 10^{-15}$ | $4.99 \times 10^{-1}$ |
| *APOE* | $4.99 \times 10^{-1}$ | $1.00 \times 10^{-20}$ | $8.65 \times 10^{-1}$ | $1.00 \times 10^{-20}$ | 1 | 1 |
| *PLTP* | $4.99 \times 10^{-1}$ | $4.29 \times 10^{-9}$ | $9.66 \times 10^{-1}$ | $6.66 \times 10^{-15}$ | $1.00 \times 10^{-2}$ | $4.99 \times 10^{-1}$ |
| *LPL* | $4.99 \times 10^{-1}$ | $4.08 \times 10^{-13}$ | $3.00 \times 10^{-3}$ | $1.00 \times 10^{-20}$ | $6.59 \times 10^{-1}$ | $4.99 \times 10^{-1}$ |
| *ANGPTL3* | $4.99 \times 10^{-1}$ | $8.86 \times 10^{-8}$ | $2.00 \times 10^{-3}$ | $1.00 \times 10^{-20}$ | $4.00 \times 10^{-3}$ | 1 |
| *ANGPTL4* | $4.99 \times 10^{-1}$ | $1.00 \times 10^{-20}$ | $9.78 \times 10^{-1}$ | $9.99 \times 10^{-1}$ | $9.89 \times 10^{-1}$ | 1 |
| *APOC1* | $4.99 \times 10^{-1}$ | $1.67 \times 10^{-16}$ | $4.99 \times 10^{-1}$ | $1.00 \times 10^{-20}$ | $9.81 \times 10^{-1}$ | $4.99 \times 10^{-1}$ |
| *APOC2* | $4.99 \times 10^{-1}$ | $3.57 \times 10^{-13}$ | $7.71 \times 10^{-1}$ | $1.11 \times 10^{-1}$ | $9.11 \times 10^{-1}$ | $4.99 \times 10^{-1}$ |
| *APOC4* | $4.99 \times 10^{-1}$ | $3.72 \times 10^{-13}$ | $7.36 \times 10^{-1}$ | $2.58 \times 10^{-14}$ | $9.73 \times 10^{-1}$ | $4.99 \times 10^{-1}$ |
| *APOB* | $4.99 \times 10^{-1}$ | $1.00 \times 10^{-20}$ | $9.99 \times 10^{-1}$ | $7.32 \times 10^{-12}$ | $1.00 \times 10^{-3}$ | 1 |
| *LMF1* | $9.98 \times 10^{-1}$ | $8.03 \times 10^{-7}$ | 1 | $3.21 \times 10^{-2}$ | $3.79 \times 10^{-5}$ | $9.98 \times 10^{-1}$ |
| *APOL1* | $4.99 \times 10^{-1}$ | $5.30 \times 10^{-2}$ | $6.40 \times 10^{-2}$ | 1 | $8.89 \times 10^{-11}$ | $4.99 \times 10^{-1}$ |
| *HBA1* | $4.99 \times 10^{-1}$ | $3.75 \times 10^{-5}$ | $9.99 \times 10^{-1}$ | $4.51 \times 10^{-1}$ | $2.46 \times 10^{-10}$ | 1 |
| *HBA2* | $4.99 \times 10^{-1}$ | $1.30 \times 10^{-5}$ | $9.99 \times 10^{-1}$ | $4.51 \times 10^{-1}$ | $3.93 \times 10^{-10}$ | $4.99 \times 10^{-1}$ |
| *B4GALT3* | $4.99 \times 10^{-1}$ | $6.80 \times 10^{-2}$ | $7.21 \times 10^{-1}$ | $4.99 \times 10^{-1}$ | $1.23 \times 10^{-6}$ | 1 |
| *KLK8* | 1 | 1 | $1.62 \times 10^{-6}$ | $9.89 \times 10^{-1}$ | $1.00 \times 10^{-3}$ | 1 |
| *PNLIP* | $4.99 \times 10^{-1}$ | $9.99 \times 10^{-1}$ | $9.26 \times 10^{-1}$ | $7.75 \times 10^{-1}$ | $1.00 \times 10^{-3}$ | $4.99 \times 10^{-1}$ |
| *WNT4* | $4.99 \times 10^{-1}$ | $9.61 \times 10^{-1}$ | $9.99 \times 10^{-1}$ | $4.99 \times 10^{-1}$ | $3.29 \times 10^{-5}$ | $4.99 \times 10^{-1}$ |
| *BACE1* | $4.99 \times 10^{-1}$ | $5.55 \times 10^{-17}$ | $2.20 \times 10^{-2}$ | $9.99 \times 10^{-16}$ | $6.69 \times 10^{-1}$ | $4.99 \times 10^{-1}$ |
| *CETP* | $4.99 \times 10^{-1}$ | $1.00 \times 10^{-3}$ | $9.99 \times 10^{-1}$ | $1.41 \times 10^{-6}$ | $9.99 \times 10^{-1}$ | $4.99 \times 10^{-1}$ |
| *PCSK6* | $4.99 \times 10^{-1}$ | 1 | $9.99 \times 10^{-1}$ | $1.83 \times 10^{-5}$ | $1.00 \times 10^{-3}$ | $4.99 \times 10^{-1}$ |
| *PCSK7* | $4.99 \times 10^{-1}$ | $1.66 \times 10^{-8}$ | $9.97 \times 10^{-1}$ | $1.00 \times 10^{-20}$ | $9.99 \times 10^{-1}$ | $4.99 \times 10^{-1}$ |
| *LCAT* | $4.99 \times 10^{-1}$ | $5.00 \times 10^{-1}$ | 1 | $6.24 \times 10^{-3}$ | $4.38 \times 10^{-1}$ | $4.99 \times 10^{-1}$ |
| *APOF* | $4.99 \times 10^{-1}$ | $5.78 \times 10^{-1}$ | $7.79 \times 10^{-1}$ | $4.10 \times 10^{-3}$ | $9.64 \times 10^{-1}$ | $4.99 \times 10^{-1}$ |
| *TYRO3* | $4.99 \times 10^{-1}$ | $9.28 \times 10^{-1}$ | $9.99 \times 10^{-1}$ | $1.20 \times 10^{-2}$ | $8.57 \times 10^{-1}$ | $4.99 \times 10^{-1}$ |

## S1 Supplemental Information

### Supplemental Subjects and Methods

#### UK Biobank Data

We downloaded individual genotype data using the UK Biobank’s (UKB) ukbgene resource, https://biobank.ctsu.ox.ac.uk/crystal/download.cgi. European individuals from the UK Biobank data were selected using the self-identified ancestry (data field 21000) using values outlined at https://biobank.ctsu.ox.ac.uk/crystal/field.cgi?id=21000. Using the relatedness file provided by the UK Biobank, one individual from each related pair was then randomly removed. This process was repeated for individuals whose self-identified ancestry was South Asian.

We performed unsupervised genome-wide ancestry estimation using ADMIXTURE by setting *K* = 3^91^ on the self-identified African ancestry cohort. We also included YRI and CEU individuals in the ADMIXTURE runs from the 1000 Genomes Project, to identify the ancestry components corresponding to African and European ancestry. We removed individuals containing less than 5% membership in the African ancestry component and more than 5% membership in the third component, which corresponds to American Indian/Alaskan Native (AIAN) ancestry (Figure S18). We downloaded imputed SNP data from the UK Biobank for all remaining individuals and removed SNPs with an information score below 0.8. Information scores for each SNP are provided by the UK Biobank (http://biobank.ctsu.ox.ac.uk/crystal/refer.cgi?id=1967). The remaining genotype and high-quality imputed SNPs were put through a stringent quality control pipeline in each ancestry cohort to obtain cohort-specific SNPs to be used for further analysis as detailed in the main text (detailed below).

We performed the following quality control filters in the European, South Asian, and African cohorts from the UK Biobank (Application number 22419). Genotype data for 488,377 individuals in the UK Biobank were downloaded using the UK Biobank’s ukbgene (https://biobank.ctsu.ox.ac.uk/crystal/download.cgi) tool and converted using the provided ukbconv tool (https://biobank.ctsu.ox.ac.uk/crystal/refer.cgi?id=149660). Phenotype data was also downloaded for those same individuals using the ukbgene tool. Individuals identified by the UK Biobank to have high heterozygosity, excessive relatedness, or aneuploidy were removed (1,550 individuals). After then separating individuals into self-identified ancestral groups using data field 21000. Within these cohorts, unrelated individuals were then selected by randomly selecting an individual from each pair of related individuals. This resulted in 349,469 European individuals, 5,716 South Asian individuals, and 4,967 African individuals.

Genotype quality control was then performed on each cohort separately using the following steps. All structural variants were first removed, leaving only single nucleotide polymorphisms in the genotype data. Next, all AT/CG SNPs were removed to avoid possible confounding due to sequencing errors. Then, SNPs with minor allele frequency less than 1% were removed using the plink2^32^ --maf 0.01. We then removed all SNPs found to be in Hardy-Weinberg equilibrium, using the plink --hwe 0.000001 flag to remove all SNPs with a Fisher’s exact test *p*-value *>* 10*^−^*^6^. Finally, all SNPs with missingness greater than 1% were removed using the plink --mind 0.01 flag.

#### Biobank Japan Data

We downloaded summary statistics for 25 quantitative traits from the Biobank Japan (BBJ) website (http://jenger.riken.jp/en/result) ^1, 93–95^. The sample descriptions and number of SNPs included in our analyses are given in Table S4. The number of SNPs included in the analysis of each trait represent those SNPs that: (*i*) contained an rsid number that could be mapped to the hg19 genome build, (*ii*) overlapped with SNPs contained within the 1000 Genomes Project phase 3 genotype data, and (*iii*) had a minor allele frequency greater than 0.01 in Japanese (JPT) individuals in the 1000 Genomes Project. We used the 1000 Genomes phase 3 data from 93 JPT individuals to estimate the linkage disequilibrium (LD) between SNPs in BioBank Japan for which we had the summary statistic data; LD was estimated separately for each of the 25 quantitative traits using the trait specific SNP arrays. LD estimates were used in the calculation of regional association statistics.

#### Population Architecture using Genomics and Epidemiology (PAGE) Study Data

Summary statistics for genotyped and imputed SNPs in five admixed populations were downloaded from the Population Architecture using Genomics and Epidemiology (PAGE) ^6^ with permission granted via approval of manuscript proposal. We included summary statistics for up to 14 quantitative traits for African-American, Hispanic and Latin American, Native Hawaiian, American Indian/Alaska Native, and Asian ancestry cohorts when available. All AT/CG SNPs were omitted, and SNPs with an IMPUTE2 information score greater than 0.8 were included in this analysis. Number of individuals and SNPs varied across ancestry-trait combinations and are given in Table S4 - Table S9.

Individuals from the 1000 Genomes Project phase 3^92^ and the Human Genome Diversity Panel (HGDP) ^108^ were used to obtain LD estimates between SNPs for each ancestry cohort. To construct the LD reference panel for PAGE summary statistics from the African-American ancestry cohort, unrelated individuals from the 1000 Genomes Americans of African Ancestry in SW USA (ASW) and African Caribbeans in Barbados (ACB) were used. Only SNPs found in both the 1000 Genomes imputed data and PAGE summary statistics files were used in gene-level association and heritability analyses. We used the same approach to compute reference LD estimates between SNPs for the Hispanic and Latin American, AIAN, and Asian ancestry cohorts, with the following 1000 Genomes reference population, respectively: Mexican Ancestry from Los Angeles USA (MXL) and Puerto Ricans from Puerto Rico (PUR); Colombians from Medellin, Colombia (CLM) and Peruvians from Lima, Peru (PEL); and the East Asian superpopulation (EAS). For the Native Hawaiian individuals from the PAGE study, there were no appropriate reference populations in the 1000 Genomes data. In order to construct a reference LD matrix for the Native Hawaiian ancestry cohort, we randomly sampled individuals from populations in the most recent release of the HGDP proportional to the global ancestry of the Native Hawaiian cohort. The Native Hawaiian cohort’s global ancestry proportions were determined using ADMIXTURE runs to be 47.89% Oceanian, 25.16% East Asian, 25.51% European, 0.90% African, and 0.54% AIAN in a separate publication (Wojcik preprint - in prep.). We did not sample from populations with less than 1% of the total ancestry in the admixture analysis referenced above. The resulting sample from which LD was estimated included 39 individuals from the Papuan Sepik in New Guinea and Melanesian in Bougainville, 14 individuals from the French in France, and 14 individuals from the Yoruba in Nigeria.

#### WHI study cohort description

The Women’s Health Initiative (WHI) is a long-term, prospective, multi-center cohort study investigating post-menopausal women’s health in the US. WHI was funded by the National Institutes of Health and the National Heart, Lung, and Blood Institute to study strategies to prevent heart disease, breast 124 cancer, colon cancer, and osteoporotic fractures in women 50-79 years of age. WHI involves 161,808 women recruited between 1993 and 1998 at 40 centers across the US. The study consists of two parts: the WHI Clinical Trial which was a randomized clinical trial of hormone therapy, dietary modification, and calcium/Vitamin D supplementation, and the WHI Observational Study, which focused on many of the inequities in women’s health research and provided practical information about incidence, risk factors, and interventions related to heart disease, cancer, and osteoporotic fractures. For this project, women who self identified as European were excluded from the study sample (dbGaP study accession number: phs000227).

#### HCHC/SOL study cohort description

The Hispanic Community Health Study / Study of Latinos (HCHS/SOL) is a multi center study of Hispanic/Latino populations with the goal of determining the role of acculturation in the prevalence and development of diseases, and to identify other traits that impact Hispanic/Latino health. The study is sponsored by the National Heart, Lung, and Blood Institute (NHLBI) and other institutes, centers, and offices of the National Institutes of Health (NIH). Recruitment began in 2006 with a target population of 16,000 persons of Cuban, Puerto Rican, Dominican, Mexican or Central/South American origin. Household sampling was employed as part of the study design. Participants were recruited through four sites affiliated with San Diego State University, Northwestern University in Chicago, Albert Einstein College of Medicine in Bronx, New York, and the University of Miami. Researchers from seven academic centers provided scientific and logistical support. Study participants who were self-identified Hispanic/Latino and aged 18-74 years underwent extensive psycho-social and clinical assessments during 2008-2011. A re-examination of the HCHS/SOL cohort is conducted during 2015-2017. Annual telephone follow-up interviews are ongoing since study inception to determine health outcomes of interest. (dbGaP study accession number: phs000555).

#### BioMe Biobank study cohort description

The Charles Bronfman Institute for Personalized Medicine at Mount Sinai Medical Center (MSMC), BioMeTM 54 BioBank (BioMe) is an EMR-linked bio-repository drawing from Mount Sinai Medical Center consented patients which were drawn from a population of over 70,000 inpatients and 800,000 outpatients annually. The MSMC serves diverse local communities of upper Manhattan, including Central Harlem (86% African American), East Harlem (88% Hispanic/Latino), and Upper East Side (88% Caucasian/White) with broad health disparities. BioMeTM 58 enrolled over 26,500 participants from September 2007 through August 2013, with 25% African American, 36% Hispanic/Latino (primarily of Caribbean origin), 30% Caucasian, and 9% of Other ancestry. The BioMeTM 60 population reflects community-level disease burdens and health disparities with broad public health impact. Biobank operations are fully integrated in clinical care processes, including direct recruitment from clinical sites waiting areas and phlebotomy stations by dedicate Biobank recruiters independent of clinical care providers, prior to or following a clinician standard of care visit. Recruitment currently occurs at a broad spectrum of over 30 clinical care sites. Study participants of self-reported European ancestry were not included in this analysis. (dbGaP study accession number: phs000925).

#### MEC study cohort description

The Multiethnic Cohort (MEC) is a population-based prospective cohort study including approximately 215,000 men and women from Hawaii and California. All participants were 45-75 years of age at baseline, and primarily of 5 ancestries: Japanese Americans, African Americans, European Americans, Hispanic/Latinos, and Native Hawaiians. MEC was funded by the National Cancer Institute in 1993 to examine lifestyle risk factors and genetic susceptibility to cancer. All eligible cohort members completed baseline and follow-up questionnaires. Within the PAGE II investigation, MEC proposes to study: 1) diseases for which we have DNA available for large numbers of cases and controls (breast, prostate, and colorectal cancer, diabetes, and obesity); 2) common traits that are risk factors for these diseases (e.g., body mass index / weight, waist-to-hip ratio, height), and 3) relevant disease-associated biomarkers (e.g., fasting insulin and lipids, steroid hormones). The specific aims are: 1) to determine the population-based epidemiologic profile (allele frequency, main effect, heterogeneity by disease characteristics) of putative causal SNPs in the five racial/ethnic groups in MEC; 2) for SNPs displaying effect heterogeneity across ethnic/racial groups, we will utilize differences in LD to identify a more complete spectrum of associated SNPs at these loci; 3) investigate gene x gene and gene x environment interactions to identify modifiers; 4) examine the associations of putative causal SNPs with already measured intermediate phenotypes (e.g., plasma insulin, lipids, steroid hormones); and 5) for SNPs that do not fall within known genes, start to investigate their relationships with gene expression and epigenetic patterns in small genomic studies. For this project, MEC contributed African American, Japanese American, and Native Hawaiian samples.(dbGaP study accession number: phs000220).

#### Fine-mapping analyses using the SuSiE framework

To perform SNP-level fine mapping analyses on a given quantitative trait, we applied Sum of Single Effects (SuSiE) variable selection software ^34^. SuSiE implements a Bayesian linear regression model on individual level data where sparse prior distributions are placed on the effect size of SNP and posterior inclusion probabilities (PIPs) are used to summarize their statistical relevance to the trait of interest. The software for SuSiE requires an input *£* which fixes the maximum number of causal SNPs to include in the model. In this work, we consider results when this parameter is chosen conservatively (*£* = 3000). We used the three cohorts for which we had genotype data from the UK Biobank (African, European, and South Asian) to test whether there was effect size heterogeneity among ancestries in the 25 traits analyzed in this study. We first selected ten independent, non-overlapping subsamples of 10,000 individuals from the European ancestry cohort and filtered out any SNPs that had a minor allele frequency of less than 0.01. For each subsample, we then applied SuSiE to each of the 25 traits and compared the effect sizes and posterior inclusion probabilities. The average number of SNPs with an effect size greater than 0.01 and average number of SNPs with a PIP greater than 0.01 for each trait across the ten cohorts are reported in Table S11. Table S11 also reports the median correlation coefficient of effect sizes and PIPs among the 45 pairwise comparisons between the 10 subsample cohorts.

We then applied SuSiE to the African and South Asian ancestry cohorts and compared their resulting effect sizes and PIPs to ten independent, non-overlapping subsamples of the European ancestry cohort. The number of SNPs with an effect size greater than 0.1 and PIPs greater than 0.01 in both the focal cohort (either African or South Asian) and at least one of the ten European ancestry subsamples of the same size are reported in Table S12 and Table S13. Also reported in these tables, are the mean number of effect sizes greater than 0.01 and PIPs greater than 0.01 across the European ancestry subsamples for each trait and the number of unique effect sizes greater than 0.01 and PIPs greater than 0.01 that were only identified in the African or South Asian ancestry cohorts. Finally, Table S12 and Table S13 report the median correlation coefficient of the African or South Asian ancestry cohort effect sizes and PIPs with the ten European ancestry subsample cohorts of the same size.

#### S1 Description of the gene-***ε*** framework

A unique feature of gene-*ε* is that it treats SNPs with spuriously associated nonzero effects as non-associated. gene-*ε* assumes a reformulated null distribution of SNP-level effects 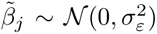, where 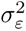 is the SNP-level null threshold and represents the maximum proportion of phenotypic variance explained (PVE) by a spurious or non-associated SNP. This leads to the reformulated SNP-level null hypothesis 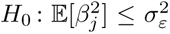. To infer an appropriate 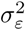, gene-*ε* fits a *K*-mixture of normal distributions over the regularized effect sizes with successively smaller variances (i.e., 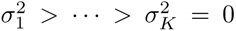). In this study as in Cheng et al. ^38^, we assume that associated SNPs will appear in the first set, while spurious and non-associated SNPs appear in the latter sets. As a final step, gene-*ε* computes its gene-level association test statistic for the *g*-th gene by conformably partitioning the regularized GWA effect size estimates and computing the quadratic form 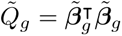. Corresponding *p*-values are then derived using Imhof’s method. This assumes the common gene-level null *H*_0_ : *Q_g_* = 0, where the null distribution of *Q_g_* is dependent upon the eigenvalues from the scaled LD matrix 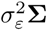. For details on implementation, validation and performance comparison with simulations, and empirical application to UK Biobank white British individuals in six traits, see Cheng et al. ^38^.

#### S2 Regression with Summary Statistics (RSS) Enrichment

Consider a GWA study with *N* individuals typed on *P* SNPs. For the *j*-th SNP, assume that we are given corresponding effect sizes 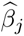 and standard error 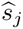 via a single-SNP linear model fit using OLS. RSS then implements the following likelihood to model the GWA summary statistics ^60^

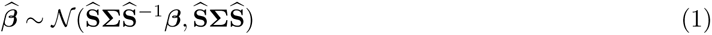

where 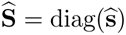 is a *J × J* diagonal matrix of standard errors, **Σ** is again used to represent some empirical estimate of the LD matrix (i.e., using some external reference panel with ancestry matching the cohort of interest), and ***β*** are the true (unobserved) SNP-level effect sizes. To model gene-level enrichment, RSS assumes the following hierarchical prior structure on the true effect sizes

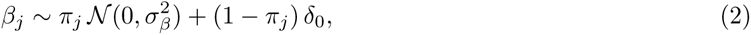

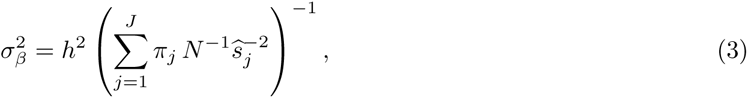

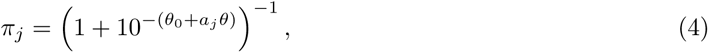

where *δ*_0_ is point mass centered at zero, *h*^2^ denotes the narrow-sense heritability of the trait, *a_j_* is an indicator detailing whether the *j*-th SNP is inside a particular gene, *θ*_0_ is the background proportion of trait-associated SNPs, and *θ* reflects the increase in probability (on the log_10_-odds scale) when a SNP within a gene has non-zero effect. Here, the authors follow earlier works ^109^ and place independent uniform grid priors on the hyper-parameters *{h*^2^*, θ*_0_*, θ}*. Note that, unlike other methods, RSS does not calculate a *P* -value for assessing gene-level association. Instead, RSS produces a posterior enrichment probability that at least one SNP in a given gene boundary is associated with the trait

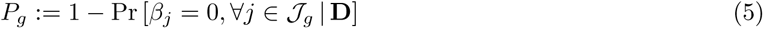

where **D** represents all of the input data including the GWA summary statistics 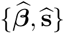, the estimated LD matrix **Σ**, and any applicable SNP annotations or weights **a** = (*a*_1_*, …, a_J_*). See ^60, 110^ for more details on preferred hyper-parameter settings. As noted in the main text, RSS is relies on a Markov chain Monte Carlo (MCMC) scheme for sampling posterior distributions and estimating model parameters. As a result, its algorithm can be subject to convergence issues if these (or the random seed) are not chosen properly.

#### S3 SNP-set (Sequence) Kernel Association Test (SKAT)

The implementation of SKAT required access to raw phenotype **y** and genotype **X** information for *N* individuals typed on *J* SNPs. To assess enrichment of the *|J_g_|* variants within gene *g*, consider the linear model with sub-matrix **X***_g_*

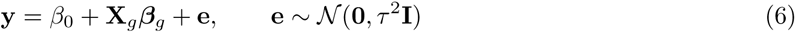

where *β*_0_ is an intercept term, ***β****_g_* = (*β*_1_*, …, β_|𝒥g|_*) is a vector of regression coefficients for the SNPs within the gene of interest, and **e** is a normally distributed error term with mean zero and scaled variance *τ* ^2^. For model flexibility, gene-specific SNP effects *β_j_* are assumed to follow an arbitrary distribution with mean zero and marginal variances 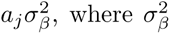 is a variance component and *a_j_* is a pre-specified weight for the *j*-th SNP. To this end, SKAT uses a variance component scoring approach and tests the null hypothesis *H*_0_ : ***β*** = **0**, or equivalently *H*_0_ : 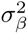 = 0. The corresponding gene-level test statistic 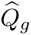 then takes on the familiar quadratic form

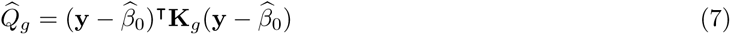

where 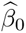 is the predicted mean of trait under the null hypothesis, and is computed by projecting **y** onto the column space of the intercept (i.e., a vector of ones). The term **K***_g_* = **X***_g_* **A***_g_* **A***_g_* **X**^T^ is commonly referred to as an *N × N* kernel matrix, where **A***_g_* = diag(*a*_1_*, …, a_|𝒥g|_*) is used to denote a diagonal weight matrix that changes for each gene *g*. Each element of **K***_g_* is computed via the linear kernel function

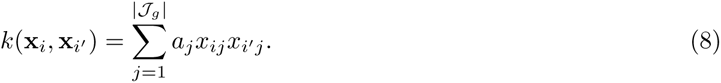

While implementing SKAT in this work, we follow previous works and set each weight to be 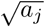 = Beta(MAF*_j_,* 1, 25) — the beta distribution density function with pre-specified parameters evaluated at the sample minor allele frequency (MAF) for the *j*-th SNP in the gene region. For more details, see ^40, 111–113^.

#### Clustering traits sharing a core set of associated genes using the WINGS algorithm

We used the WINGS algorithm ^114^ to identify clusters of traits sharing a core set of genes enriched for associated mutations. WINGS takes as input a gene (*M*) by trait (*N*) matrix and uses the Ward distance metric to find the distance among vectors of gene scores for different phenotypes; in this study, we used gene-*ε* gene-level association statistics as the input to WINGS. The more significantly associated genes that two traits share, the closer they will be in the gene-dimensional space. Applying WINGS to a matrix of gene scores for each ancestry separately, we examined whether the same traits clustered together, separately in each ancestry. We constructed matrices of gene-*ε* gene-level association statistics for the UK Biobank European, African, South Asian (from the UK Biobank) and East Asian (Biobank Japan) ancestry cohorts. Each of these matrices contained gene-level association statistics for all 25 quantitative traits of interest. The total number of genes and regulatory regions included were: European (23,603), African (23,575), South Asian(23,671), and East Asian (21,435). For the East Asian ancestry cohort, we limited the genes to the intersection of genes with gene-*ε* gene-level association statistics across all 25 traits. The number of gene scores calculated for each trait in the East Asian ancestry cohort varies due to the heterogeneity in imputed and genotype SNP arrays in the Biobank Japan studies (Table S4 and Table S15). Figure S19 shows the resulting dendrograms displaying prioritized phenotypes identified using the WINGS algorithm on each cohort’s gene score matrix. The WINGS algorithm is designed to run on 25 phenotypes or more (see McGuirl et al. ^114^ for details), and we therefore did not apply the WINGS algorithm to the AIAN, Native Hawaiian, or Hispanic and Latin American cohorts as there was not data for enough phenotypes (Table S5-Table S9).

#### Analysis of GWAS Catalog Metadata and Previous GWA Publications

We cross-referenced our results from association testing at multiple genomic scales against previously published results in the GWAS catalog (https://www.ebi.ac.uk/gwas/) and in PubMed using the following processes.

In order to collect PubMed IDs (PMIDs) for publications associated with the UK Biobank, a two-part data collection process was used. The first process was to directly search for publications with variations of the term “UK Biobank” (e.g., U.K. Biobank, United Kingdom Biobank) from PubMed using the Entrez Programming Utilities (E-Utilities) API. The E-Utilities API is the public API to the NCBI Entrez system and allows direct access to all Entrez databases including PubMed. Search queries were formulated by narrowing publications using year published and then further narrowing to those publications with variations of the search term “UK Biobank” in either the title or abstract. The open-source Python package Entrez (https://biopython.org/DIST/docs/api/Bio.Entrez-module.html) from the Biopython Project was used to facilitate interaction with the E-Utilities API.

The second data collection process was to gather information from publications listed directly on the UK Biobank website (https://www.ukbiobank.ac.uk/). Since the majority of publications on the website did not have an easily accessible PMID, identifying information including publication title and year was scraped and used to retrieve a publication’s corresponding PMID (again using the E-Utilities API). The HTML/XML document parsing Python library Beautiful Soup (https://www.crummy.com/software/BeautifulSoup/bs4/doc/) was used to parse the HTML of the various UK Biobank webpages, and the Python Requests library (https://requests.readthedocs.io/en/master/) was used to programatically send HTTP calls to the server hosting the website. PMIDs were retrieved directly from the XML output of the E-Utilities API calls.

The PMIDs retrieved from both processes were aggregated into a single set of unique PMIDs, as some publications were identified by both processes. Publications that could not get associated PMIDs from the second data collection process were flagged for manual processing. The PMIDs that were retrieved from PubMed directly but could not be found based on the publication information provided on the UK Biobank website were noted. Conversely, the PMIDs that could be retrieved from publication information found on the UK Biobank website but not directly from PubMed were also noted.

Using the compiled list of PMIDs, analyses of the UK Biobank data set reported in the GWAS catalog association data were compiled. Previous genotype-to-phenotype association data and sample ancestry descriptions were downloaded from https://www.ebi.ac.uk/gwas/docs/file-downloads. Unique genotype-to-phenotype associations were parsed using a set of custom python scripts. All scripts used in the curation of PMIDs, parsing of GWAS catalog summary data, and determination of previously published genotype-to-phenotype associations from UK Biobank studies are available on GitHub (https://github.com/ramachandran-lab/redefining_replication).

#### Simulation design to test the power and false discovery rate of GWA and gene-level association analyses

##### Simulations of a single population

In our simulation studies, we used the following general simulation scheme to generate quantitative traits using real genotype data on chromosome 1 from *N* randomly sampled individuals of European ancestry in the UK Biobank. This pipeline follows from previous studies ^38, 78^. We will use **X** to denote the *N × J* genotype matrix, with *J* denoting the number of single nucleotide polymorphisms (SNPs) encoded as 0, 1, 2 copies of a reference allele at each locus and **x***_j_* representing the genotypic vector for the *j*-th SNP. Following quality control procedures detailed in the Supplemental Information, our simulations included *J* = 36,518 SNPs distributed across genome. We used the NCBI’s RefSeq database in the UCSC Genome Browser to assign SNPs to genes which resulted in *G* = 1,408 genes in the simulation studies.

After the annotation step, we simulated phenotypes by first assuming that the total phenotypic variance V[**y**] = 1, and that all observed genetic effects explained a fixed proportion of this value (i.e., narrow-sense heritability, *h*^2^). Next, we randomly selected a certain percentage of genes to be enriched fotr associations and denoted the sets of SNPs that they contained as *C*. Within *C*, we selected causal SNPs in a way such that each associated gene at least contains one SNP with non-zero effect size. Quantitative continuous traits were then generated under the following two general linear models:

*•* Standard Model: 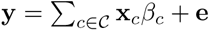
*•* Population Structure Model: 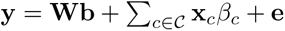

where **y** is an *N* -dimensional vector containing all the phenotype states; **x***_c_* is the genotype for the *c*-th causal SNP; *β_c_* is the additive effect size for the *c*-th SNP; and **e** *∼ 𝒩* (0*, τ* ^2^**I**) is an *N* -dimensional vector of normally distributed environmental noise. Additionally, in the model with population structure, **W** is an *N × M* matrix of the top *M* = 10 principal components (PCs) from the genotype matrix and represents additional population structure with corresponding fixed effects **b**. The effect sizes of SNPs in genes enriched for associations are randomly drawn from standard normal distributions and then rescaled so they explain a fixed proportion of the narrow-sense heritability V[ **x***_c_β_c_*] = *h*^2^. The coefficients for the genotype PCs are also drawn from standard normal distributions and rescaled such that V[**Wb**] = 10% of the total phenotypic variance, with the variance of all non-genetic effects contributing V[**Wb**] + V[**e**] = (1 *− h*^2^). For any simulations conducted under the population structure model, genotype PCs are not included in any of the model fitting procedures, and no other preprocessing normalizations were carried out to account for the additional population structure. More specifically, GWA summary statistics are then computed by fitting a single-SNP univariate linear model via ordinary least squares (OLS):

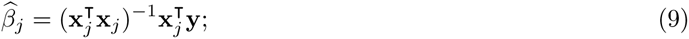

for every SNP in the data *j* = 1*, … J*. These OLS effect size estimates, along with an empirically LD matrix **Σ** computed directly from the full *N × J* genotype matrix **X**, are given to gene-*ε*. We also retain standard errors and *p*-values for the implementation of competing methods: RSS ^60^, SKAT^40^, and the standard GWA SNP-level association test. Given the simulation procedure above, we simulate a wide range of scenarios for comparing the performance of gene-level association approaches by varying the following parameters:

*•* Number of individuals: *N* = 5,000 and 10,000;
*•* Narrow-sense heritability: *h*^2^ = 0.2 and 0.6;
*•* Percentage of enriched genes: 1% (sparse) and 10% (polygenic);

Furthermore, we set the number of causal SNPs with non-zero effects to be some fixed percentage of all SNPs located within the designated genes enriched for associations. We set this percentage to be 0.125% in the 1% associated SNP-set case, and 3% in the 10% associated SNP-set case. All performance comparisons are based on 100 different simulated runs for each parameter combination. Lastly, for each simulated dataset, we also selected some number of intergenic SNPs (i.e., SNPs not mapped to any gene) to have non-zero effect sizes. This was done to mimic genetic associations in unannotated regulatory elements. Specifically, five randomly selected intergenic SNPs were given non-zero contributions to the trait heritability in the 1% enriched genes case, and 30 intergenic SNPs were selected in the 10% enriched genes case.

All performance comparisons are based on 100 different simulated runs for each parameter combination. We computed gene-level *p*-values for gene-*ε*, SKAT, and the single-SNP GWAS. For evaluating the performance of RSS, we compute posterior enrichment probabilities. For all approaches, we assessed the power and false discovery rates when identifying enriched genes at either a Bonferroni-corrected threshold (*p* = 0.05*/*1, 408 genes = 3.55 *×* 10*^−^*^5^) or according to the median probability model (posterior enrichment probability *>* 0.5) ^115^. Figure S4 and Figure S5 show the mean performances (and standard errors) across all simulated replicates.

###### Simulations of genetic trait architecture in two populations

We used the African (UKB) cohort and a subset of the European cohort and simulation studies to test the ability of GWAS and gene-*ε* to detect shared causal SNPs (in the case of gene-*ε*, genes containing causal SNPs) in a multi-ancestry study. Using the same simulation protocol as that described for testing power of different enrichment analysis methods, described in *Simulations in a single population*, we labeled all genes containing at least one causal SNP as ”causal”. We first determined the power of gene-*ε* to identify SNPs or genes that are causal in each cohort under a variety of genomic architectures. The total amount of variance explained in the phenotype by the causal SNPs (i.e. the narrow-sense heritability) to be equal to 0.2 or 0.6. In each of these contexts, the sparsity of causal variants as a function of the total number of variants was set to either 0.1 or 0.5. These values of causal SNP sparsity were selected in order to ensure that an ample number of SNPs were associated with the phenotype in both cohorts. Finally, the overlap in causal SNPs between the two cohorts was tested at proportions equal to 0 (no overlap in causal between SNPs cohorts) 0.25, 0.5, and 1 (complete overlap in causal SNPs between cohorts). For each of these parameter sets, 50 replicate simulations were performed of two cohorts derived from 10,000 European individuals and 4,967 African individuals, respectively. We summarize the performance of the standard GWA framework and gene-*ε* across the parameter space. Generally, gene-*ε* performs better on the European cohort than it does in the African cohort, but is better powered in the African cohort when the causal SNPs are the same in both cohorts (Figure S6 and Figure S7). Additionally, gene-*ε* performs better when identifying causal genes that are shared between the two cohorts - particularly when traits have high heritability Figure S8 - Figure S9.

## Appendix

### S4 SNP-level results for height and C-reactive protein

In Figure S3a and Figure S3d, we found that, across 25 traits analyzed, height had the greatest number of genome-wide significant SNP-level associations (76,910 unique associations) in at least one ancestry. Of these SNP-level associations, 8.90% (7,377 SNPs) replicate based off of rsID in at least two ancestry cohorts. Height is not the only trait in which the standard GWA SNP-level association test detects associations that replicate extensively across ancestries. In fact, SNP-level associations replicate in each of the 25 continuous traits that we analyze in this study.

We analyzed SNP-level associations with C-reactive protein in six ancestry cohorts: African-American (PAGE), European, South Asian, East Asian, Native Hawaiian, and Hispanic and Latin American cohorts. C-reactive protein is an example of a trait with a sparse and highly conserved genetic architecture across ancestries, as shown in Figure 2. Many SNPs within the *CRP* gene have been previously associated with C-reactive protein plasma levels ^116–118^. In our analysis, rs3091244 is genome-wide significant in only the European ancestry cohort, and has been functionally validated as influencing C-reactive protein levels ^52, 53^. The SNP rs3091244 is located in a promoter region slightly upstream of *CRP*, and it has clinical implications for both atrial fibrillation ^119^ and lupus erythematosus ^120^ (European *p* = 1.54 *×* 10*^−^*^116^; East Asian *p* = 1.15 *×* 10*^−^*^9^).

We expanded our search for replicated GWA SNP-level association signals across ancestry cohorts by scanning for 1 Mb regions that contained associations to the same phenotype in two or more ancestries— a process often referred to as “clumping”. These windows were centered at every unique genome-wide significant SNP in any ancestry for a given trait (we refer the 1Mb window around the significant SNP as a “clump”, Figure S3b and Figure S3e). In addition to the largest number of unique SNP-level associations, height also had the largest proportion of clumps containing a significant SNP-level GWA association signal that replicated in at least two ancestry cohorts (see Figure S3b and Figure S3e). The three traits with the greatest proportion of clumps containing SNP-level GWA signals that replicate in multiple ancestry cohorts were height (77.09% of clumps), urate (65.89%), and low density lipoprotein (54.40%).

In addition to the SNP-level associations on chromosome 1 surrounding the *CRP* gene across all six ancestry cohorts (displayed in Figure 2), there are other regions of the genome that contain significant GWA associations with C-reactive protein that replicate in multiple ancestry cohorts. On chromosome 2, there is a cluster of four SNPs significantly associated with C-reactive protein levels in the European, East Asian, and Hispanic and Latin American ancestry cohorts. Of these, rs1260326 (European *p* = 1.01 *×* 10*^−^*^55^; East Asian *p* = 1.70 *×* 10*^−^*^9^; Hispanic and Latin American *p* = 1.24 *×* 10*^−^*^20^), rs780094 (European *p* = 9.95 *×* 10*^−^*^51^; East Asian *p* = 1.70 *×* 10*^−^*^9^; Hispanic and Latin American *p* = 1.14 *×* 10*^−^*^16^), and rs6734238 (African-American (PAGE) *p* = 3.04 *×* 10*^−^*^10^; European *p* = 8.38 *×* 10*^−^*^34^; South Asian *p* = 2.17 *×* 10*^−^*^9^) were statistically significant in three of the six ancestry cohorts that we analyzed. Each of these three SNPs has been previously associated with C-reactive protein levels in a European ancestry cohort ^121–123^. Of these three SNPs, only one (rs6734238) had previously been replicated in other ancestries (in African-American, and Hispanic and Latin American cohorts ^124^).

On chromosome 19 there are 23 SNPs that are associated with CRP in the African-American PAGE, European, and Hispanic and Latin American ancestry cohorts. Two other SNPs are associated with C-reactive protein in the African-American (PAGE), European, and Hispanic and Latin American cohorts, as well as the East Asian ancestry cohort. One of these two SNPs, rs7310409 (African-American (PAGE) *p* = 8.57 *×* 10*^−^*^9^; European *p* = 3.57 *×* 10*^−^*^210^; East Asian *p* = 2.72 *×* 10*^−^*^27^; Hispanic and Latin American *p* = 5.35*×*10*^−^*^29^) located in the HNF1 homeobox A (*HNF1A*) gene, has been previously associated with C-reactive protein levels in only a European ancestry cohort ^122, 123^. Three additional significant SNPs in our analysis have been previously associated with European ancestry cohorts in previous studies, including: rs1169310^124^ (European *p* = 1.52 *×* 10*^−^*^172^; East Asian *p* = 1.28 *×* 10*^−^*^18^; Hispanic and Latin American *p* = 1.17 *×* 10*^−^*^27^), rs1183910^121, 125^ (European *p* = 5.50 *×* 10*^−^*^177^; East Asian *p* = 3.16 *×* 10*^−^*^29^; Hispanic and Latin American *p* = 7.47 *×* 10*^−^*^29^), and rs7953249^126^ (European *p* = 1.19 *×* 10*^−^*^177^; East Asian *p* = 1.10 *×* 10*^−^*^19^; Hispanic and Latin American *p* = 4.80 *×* 10*^−^*^29^). Two SNPs, rs2259816 (European *p* = 2.77 *×* 10*^−^*^172^; East Asian *p* = 9.33 *×* 10*^−^*^18^; Hispanic and Latin American *p* = 1.90 *×* 10*^−^*^27^) and rs7979473 (African *p* = 1.49 *×* 10*^−^*^9^; East Asian *p* = 6.06*×*10*^−^*^29^; Hispanic and Latin American *p* = 1.56*×*10*^−^*^30^), have been previously associated with C-reactive protein in both African-American and Hispanic and Latin American ancestry cohorts ^124^. There is one final group of three SNPs associated with C-reactive protein in the African-American (PAGE), European, East Asian, and Hispanic and Latin American ancestry cohorts on chromosome 19. One of them, rs4420638 (East Asian *p* = 9.93 *×* 10*^−^*^29^; Hispanic and Latin American *p* = 2.03 *×* 10*^−^*^30^), has been previously associated in a European ancestry cohort ^121, 123, 125^. These four regions indicate a highly conserved SNP-level architecture of C-reactive protein across six ancestry cohorts. Interestingly, we were unable to replicate associations with C-reactive protein across ancestries at the gene or pathway levels.

### Gene and pathway association results

Three genes, *GP6*, *RDH13*, and *AGPAT5*, were significantly associated with platelet count (PLC) in the African-American (PAGE) ancestry cohort and the East Asian ancestry cohort (Figure S10. Of these, no significant SNPs in the glycoprotein VI platelet (*GP6*) gene have been reported in the GWAS catalog for either ancestry cohort. However, a single SNP within *GP6*, rs1613662, has previously been associated with mean platelet volume in a GWA study analyzing a European ancestry cohort ^127^. *GP6* plays a critical role in platelet aggregation, and mutations have been previously associated with fetal loss ^128^. The retinol dehydrogenase 13 (*RDH13*) gene has no reported GWAS catalog associations with platelet count, but is within 60kb of a SNP significantly associated with platelet aggregation ^129^. Of the three genes significantly associated with PLC in both the European and AIAN cohorts, 1-Acylglycerol-3-Phosphate O-Acyltransferase 5 (*AGPAT5*) is a member of a gene family known to play a role in immunity and inflammation response ^130^. Alcohol dehydrogenase 2 (*ALDH2*) has additionally been associated with hypertension in an elderly Japanese cohort ^131^. A member of the RAS oncogene family (*RAB8A*) has been shown to play a role in the inhibition of inflammatory response. In contrast, the cut like homeobox 2 *CUX2* gene contains a significantly associated SNP in the array used in this study for the East Asian ancestry cohort, but it has no previous associations in a European ancestry cohort. However, *CUX2* is significantly associated at the gene-level in both the European and East Asian ancestry cohorts. Although not reported as being associated with PLC in the GWAS Catalog, a single SNP, rs61745424 which encodes a missense mutation, has been previously identified as being related to the trait ^132^. The gene-*ε* association statistics for the seven genes significantly associated with PLC are available in Table S17.

Finally, a single gene, acyl-CoA dehydrogenase family member 10 (*ACAD10*) associated in our gene-level analysis of PLC, was significant in both the European and East Asian ancestry cohorts (European gene- *ε p* = 1.47 *×* 10*^−^*^10^; East Asian gene-*ε p* = 2.00 *×* 10*^−^*^10^) but contained no previous associations in the GWAS catalog. The African-American and Hispanic and Latin American ancestry cohorts analyzed in Qayyum et al. ^133^ both contain SNPs within *ACAD10* that are significantly associated with PLC.

In our analysis of triglyceride levels in six ancestry cohorts (African-American (PAGE), European, East Asian, South Asian, Hispanic and Latin American, and Native Hawaiian), we identified shared genetic architecture at the SNP, gene, and subnetwork level. Replicated SNPs and genes between the six ancestry cohorts are shown in Figure S12-Figure S13. We focus our discussion of results at the network level in the European, East Asian, and Native Hawaiian ancestry cohorts (Figure 3). In the European and East Asian ancestry cohorts, we identified 55 shared genome-wide significant associations at the gene-level. Of these results, eight genes lie in the same significantly mutated subnetwork (Hierarchical HotNet *p <* 10*^−^*^3^) when analyzing each ancestry cohort independently. Five of those eight genes belong to the apolipoprotein family of genes, including: apolipoprotein A1 (*APOA1*), apolipoprotein A4 (*APOA4*), apolipoprotein A5 (*APOA5*), apollipoprotein C3 (*APOC3*), apolipoprotein E (*APOE*). Specifically, the apolipoprotein play a central role in lipoprotein biosynthesis and transport. All of these genes contain SNPs previously associated with triglyceride levels in a European ancestry cohort ^98–103^. All five genes also contain SNPs previously associated with triglyceride levels in non-European ancestry cohorts. Specifically, *APOA1*, *APOC3*, and *APOE* each contain SNPs previously associated with triglyceride levels in African-American and Hispanic and Latin American ancestry cohorts^98, 99^. The *APOA5* gene has previously been associated to triglyceride levels in an East Asian, African-American, and Hispanic and Latin American ancestry cohorts^100, 104^.

The other three genes that were significantly associated with triglyceride levels in the European and East Asian ancestry cohorts are members of the largest significantly mutated subnetwork including phospholipid transfer protein (*PLTP* ; European gene-*ε p* = 4.29 *×* 10*^−^*^9^; East Asian gene-*ε p* = 6.66 *×* 10*^−^*^15^), lipoprotein lipase (*LPL*; European gene-*ε p* = 4.08 *×* 10*^−^*^13^; East Asian gene-*ε p* = 1.00 *×* 10*^−^*^20^), and angiopoietin like 3 (*ANGPTL3* ; European gene-*ε p* = 8.86 *×* 10*^−^*^8^; East Asian gene-*ε p* = 1.00 *×* 10*^−^*^20^). *PLTP* has previously been associated with triglyceride levels in European, African-American, and Hispanic and Latin American ancestry cohorts^97–100, 105–107, 134^. *LPL* is one of the most well-studied genes in the regulation of triglyceride levels. It has previously been associated with triglyceride levels in European ancestry cohorts^96–103, 105–107^, East Asian ancestry cohorts^104, 147^, and African ancestry cohorts as well as Hispanic and Latin American ancestry cohorts^96–98, 100, 105, 145, 146, 148, 149^. The final gene that was genome-wide significant in both the European and East Asian ancestry cohorts, *ANGPTL3*, has no previous associations in the GWAS catalog and presents a novel candidate gene within the network. While not significant in any gene-level analysis, the gene *ANGPTL4* (European gene-*ε p* = 1.00 *×* 10*^−^*^20^; East Asian gene-*ε p* = 9.99 *×* 10*^−^*^1^) from the same family is present in the largest subnetwork in the European cohort and also has also been previously identified as having associations in European, African, and Hispanic and Latin American ancestry cohorts^96, 97, 100, 145, 146, 150^.

In our analysis of the European ancestry cohort from the UK Biobank, we additionally identified a set of eight genes that are connected to the core network discussed above. One of these genes is *ANGPTL4*, which we discussed above. Five of these genes were significant at the gene-level in the European ancestry cohort, including four apoliprotein genes (*APOC1* ; European gene-*ε p* = 1.67 *×* 10*^−^*^16^, *APOC2* ; European gene- *ε p* = 3.57*×*10*^−^*^13^, *APOC4* ; European gene-*ε p* = 3.72*×*10*^−^*^13^, and *APOB* ; European gene-*ε p* = 1.00*×*10*^−^*^20^) and lipase maturation factor 1 (*LMF1* ; European gene-*ε p* = 8.03 *×* 10*^−^*^7^). Each of these genes have been previously associated with triglyceride levels in a European ancestry cohort^100^. Additional associations were also found in that same study which conducted a meta-analysis of European, African-American, and Hispanic and Latin American ancestry cohorts. The final two genes included in the significantly mutated subnetwork of the European ancestral cohort, *APOL1* and *HBA1*, were not were not identified as genome-wide significant by gene-*ε* and have no previous SNP-level associations with triglyceride levels in the GWAS Catalog. Interestingly, both *APOL1* (Native Hawaiian gene-*ε p* = 8.89*×*10*^−^*^11^) and *HBA1* (Native Hawaiian gene-*ε p* = 2.46*×*10*^−^*^10^) were both identified as genome-wide significant by gene-*ε* in our analysis of the Native Hawaiian ancestry cohort and the interaction between them was identified in our Hierarchical HotNet ^31^ analysis as present in both the European and Native Hawaiian ancestry cohorts.

In addition to *APOL1* and *HBA1*, six more genes are connected to the core network of genes that overlap in the East Asian and European significantly mutated subnetworks. Of these, both *HBA2* and *B4GALT3* are significant at the gene-level in the Native Hawaiian ancestry cohort alone. They are each connected to genes identified in both the European and Native Hawaiian ancestry cohorts as members of the largest significantly mutated subnetwork. The final three genes include kallikrein related peptidase 8 (*KLK8*), pancreatic lipase *PNLIP*, and wnt family member 4 (*WNT4*) which were not significant at the gene-level and did not contain previous SNP-level associations in the GWAS catalog.

In the largest significantly mutated subnetwork identified in our analysis of the East Asian ancestry cohort, we identified seven genes that were not shared by the networks in other ancestry cohorts. One of these genes, beta-secretase 1 (*BACE1* ; East Asian gene-*ε p* = 3.57 *×* 10*^−^*^13^; European gene-*ε p* = 5.55 *×* 10*^−^*17), was significant at the gene-level but contained no previously associated SNPs in any cohort in the GWAS catalog. *BACE1* plays a role in the metabolism of amyloid beta precursor protein ^151^. Three of the genes within the network identified in the East Asian ancestry cohort contain previously associated SNPs in both European and non-European ancestry cohorts, including: cholesteryl ester transfer protein (*CETP*), proprotein convertase subtilisin/kexin type 6 (*PCSK6*), and proprotein convertase subtilisin/kexin type 7 (*PCSK7*) ^98, 105^. The final three genes in the significantly mutated subnetwork identified in the East Asian ancestry cohort were not significant at the gene-level and do not contain previously associated SNPs in the GWAS catalog in any ancestral cohort. Lecithin-cholesterol acyltransferase (*LCAT*) is involved in cholesterol biosynthesis and apolipoprotein F (*APOF*) encodes one of the minor apolipoprotein genes present in plasma. Finally, tyrosine-protein kinase receptor 3 (*TYRO3*) plays a role in ligand recognition and cell metabolism ^152^. The gene-*ε p*-values in each ancestry cohort for each of the 28 genes discussed here are shown in Table S18.

## Notes

### Competing Interest Statement

C.G. owns stock in 23andMe. E.K. and C.G. are members of the scientific advisory board for Encompass Bioscience. E.K. consults for Illumina.

